# The phylogenetic position of the Yunxian cranium elucidates the origin of *Homo longi* and the Denisovans

**DOI:** 10.1101/2024.05.16.594603

**Authors:** Xiaobo Feng, Qiyu Yin, Feng Gao, Dan Lu, Qin Fang, Yilu Feng, Xuchu Huang, Chen Tan, Hanwen Zhou, Qiang Li, Chi Zhang, Chris Stringer, Xijun Ni

**Affiliations:** School of History and Culture, Shanxi University, Taiyuan, 030006, China; Hubei Polytechnic University, Huangshi, 435003, China; Institute of Yunxian Man Site, Hanjiang Normal University, Shiyan, 442000, China; School of History Culture and Tourism, Hanjiang Normal University, Shiyan, 442000, China; Institute of Vertebrate Paleontology and Paleoanthropology, Chinese Academy of Sciences, Beijing, 100044, China; College of Earth and Planetary Sciences, University of Chinese Academy of Sciences, Beijing, 100049, China; Yunnan Institute of Cultural Relics and Archeology, Kunming 650118, China; Hubei Provincial Institute of Cultural Relics and Archaeology, Wuhan, 430077, China; School of History of Wuhan University, Wuhan, 430072, China; Museum of Shiyan, Shiyan, 442000, China; Poly Art Research Institute, Beijing, 100010, China; School of Earth Sciences, China University of Geosciences, Wuhan 430074, China; Centre for Human Evolution Research, Natural History Museum, London, UK; PaleoAnthropological Research Center, Fudan University, Shanghai, 200433, China

## Abstract

Diverse Middle Pleistocene forms of *Homo* coexisted in Africa, Europe, and Asia, and it’s controversial whether these fossil humans represent different species or clades. The ∼1 Ma old Yunxian 2 fossil from China is crucial for understanding the cladogenesis of *Homo* and the origin of *Homo sapiens*. Here, we restored and reconstructed the distorted Yunxian 2 cranium using new technology. The results show that this cranium displays mosaic features of plesiomorphy and apomorphy. Geometric-morphometric, phylogenetic and Bayesian tip-dating analyses including the reconstructed Yunxian 2 suggest that it is an early member of the Asian *Homo longi* clade, which probably includes the Denisovans, and is the sister group to the *Homo sapiens* clade. Both the *H. sapiens* and *H. longi* clades had deep roots extending beyond the Middle Pleistocene, and the basal position of the Yunxian fossil cranium suggests that it represents a population lying close to the last common ancestor of the two clades.

**One-Sentence Summary:** The newly-reconstructed Yunxian 2 cranium represents a basal member of the *Homo longi* and Denisovan clade, and probably lies close to the last common ancestor of that clade and the clade of *H. sapiens*.

## Main Text

Middle Pleistocene (Chibanian) human fossils show high morphological diversity. Some of those discovered recently, such as from Callao Cave in the Philippines, Rising Star Cave in South Africa, and Harbin in China, are so distinctly different from other fossil humans that they have been proposed as new species (*1-3*), additional to previously known taxa such as *H. sapiens, H. neanderthalensis, H. heidelbergensis, H. erectus*, and *H. habilis*. It remains a matter of intense debate whether these morphologically diverse archaic humans represent multiple evolutionary clades, or whether some are transitional variants leading to *H. sapiens*. Chibanian human fossils from China, sometimes called “archaic *Homo sapiens*” have been at the center of fierce debates about their relationship to *H. sapiens* over many years. Because of their even greater antiquity (*4, 5*) and apparent possession of both primitive features of *H. erectus* and arguably derived features of *H. sapiens* (*6, 7*), the two fossil crania from Yunxian, China (Yunxian 1 and 2) are crucial for interpreting the Chibanian record in China and reconstructing the evolution of *Homo* more generally. The best-preserved of the two crania, Yunxian 2, is nevertheless distorted, which has hampered attempts to infer its phylogenetic position. Here we reconstructed Yunxian 2, also using elements from Yunxian 1, and investigated its phylogenetic position. Given its geological age of 0.94-1.10 Ma (*5*) and the presence of mosaic features of plesiomorphy and apomorphy, Yunxian 2 probably lies close to the last common ancestor of the clades containing *H. sapiens* and *H. longi* (hereafter called the sapiens and longi clades), and can also elucidate the evolutionary history of the Denisovans.

Our understanding of human evolution is largely based on fossil cranial specimens, but many of these are incompletely preserved and/or distorted. Proper reconstruction of these imperfect fossils is therefore critical to studying their phylogenetic relationships. Reconstructions of material such as *Sahelanthropus, Ardipithecus, Paranthropus, H. habilis*, and the Chibanian Steinheim and Ceprano *Homo* crania have already provided essential evidence regarding the early stages of human evolution (*8-15*).

The Yunxian human fossils were discovered at a site on a terrace of the Hanjiang River in Yunxian County, Hubei Province, in central China. The site has yielded three crania so far (including a recently discovered and still unprepared cranium (*16*)). Two previously known Yunxian crania, Yunxian 1 and 2, are both distorted. Yunxian 1 is crushed, with obvious plastic deformation. The distortion of Yunxian 2 is much less severe, and evaluation of high resolution CT scans indicates that the primary distortion is due to fragmentation and displacement rather than plastic deformation (Supplementary Materials). In a previous study, this cranium was virtually reconstructed mainly through keystone and symmetry corrections (*17*). The new reconstruction reported here is based on the least distorted cranium Yunxian 2, but also uses some data from Yunxian 1. The CT scans showed that the distortion of Yunxian 2 consisted mainly of small fractures and displacement of large fragments. We used methods developed in recent years (*8-13*), including virtual segmentation, separation and repositioning of the fragments, to correct the distortion (see Supplementary Material for detailed reconstruction procedure and evaluation of the variations of different versions of reconstructions).

As reconstructed, the Yunxian 2 cranium is large and long, with a platycephalic braincase in lateral view. It is smaller than the Harbin cranium (*H. longi*) and Xuchang, about the same size as Kabwe, Jinniushan, Petralona and Bodo, and larger than the Sangiran 17 (*H. erectus*), Irhoud 1 and Maba specimens in overall dimensions. After reconstruction, the fossil lacks only small parts of the zygomatic arches and the missing central incisors.

The cranium is clearly plesiomorphic in overall form, with a broad supraorbital torus, basicranium and palate, a long and low vault in lateral view, a receding frontal, and rather flat parietal contour. However, it lacks both the strongly angulated occipital with a prominent transverse torus found in *H. erectus* and some Chibanian African and European *Homo* crania, and the protruding occipital region (occipital bun) with a central suprainiac fossa typical of Neanderthals. In posterior view, the slightly keeled cranium is widest in the supramastoid region, below which the large mastoid processes slope inward. Additionally, the temporals and parietals converge superiorly, although not as strongly as in *H. erectus* fossils, and there is neither the upper parietal expansion found in recent *H. sapiens*, nor the “en bombe” shape typical of Neanderthals. In lateral view the face is high and projects anteriorly as in *H. erectus* and Chibanian African and European *Homo* fossils, but to a slightly smaller degree. The nasal bones also project strongly anteriorly, but the midface is not pulled forwards as in Neanderthals. The upper face and nasal aperture are wide. The zygomaxillary region is transversely flat and faces anteriorly, more comparable to the condition in European *H. antecessor* and the Asian Harbin, Dali, Jinniushan and Hualongdong fossils, as well as in *H. sapiens*. However, the cheek bones are large and high.

The matrix enclosed in the endocranial cavity of Yunxian 2 is very dense and the bone-matrix contrast of the CT scan is low. As a result, anatomical details such as the cranial vessel impressions on the inner surface of the cranial bones of Yunxian 2 cannot yet be reconstructed. Nevertheless, by setting 28 landmarks and measuring bone thickness, we were able to reconstruct the gross endocranial impressions and show that Yunxian 2 has a moderate endocranial capacity of about 1143 cc, similar to Dali, Hualongdong, Arago 21, Petralona and Ceprano. Its frontal lobe is low and narrow, barely more expanded than those of *H. erectus* and Chibanian African and European *Homo* crania, and significantly less expanded than those of Neanderthals (including the Sima de los Huesos fossils (*18*)) and *H. sapiens*. The posterior parts of the parietal and occipital lobes are laterally and posteriorly expanded to a degree similar to Kabwe, Petralona, Dali, Jinniushan, Xuchang, and Neanderthals. The dorsal expansion of the parietal lobe of Yunxian 2 is much less than that of Neanderthals and *H. sapiens*.

Geometric morphometric analysis based on 276 landmarks and semilandmarks (SI) shows that the 24 most complete hominin fossils and 153 recent human specimens compared with the reconstructed Yunxian 2 clustered into 4 main groups in the morphospace of bgPCA 1 and bgPCA 2 (Fig. 3). Yunxian 2 is close to the group containing Dali, Jinniushan and Harbin, and further from the *H. erectus* group. As the shape extremes indicate, lower bgPCA 1 values represent a more retracted and smaller face and more globular neurocranium, while higher bgPCA 1 values represent a larger and more projecting face, low and small neurocranium, stronger supraorbital torus, and stronger occipital torus. Lower bgPCA 2 values represent a narrower face, higher neurocranium and more vertical mastoid process, while higher bgPCA values represent a broader and lower face, stronger supraorbital torus, lower but more elongated neurocranium, more medially inclined mastoid process, and more protruding and angled occipital region. Dali, Jinniushan, Harbin and Yunxian 2 have moderate bgPCA 1 values, but higher bgPCA 2 values, grouping them in a distinctive position.

**Fig. 1.**
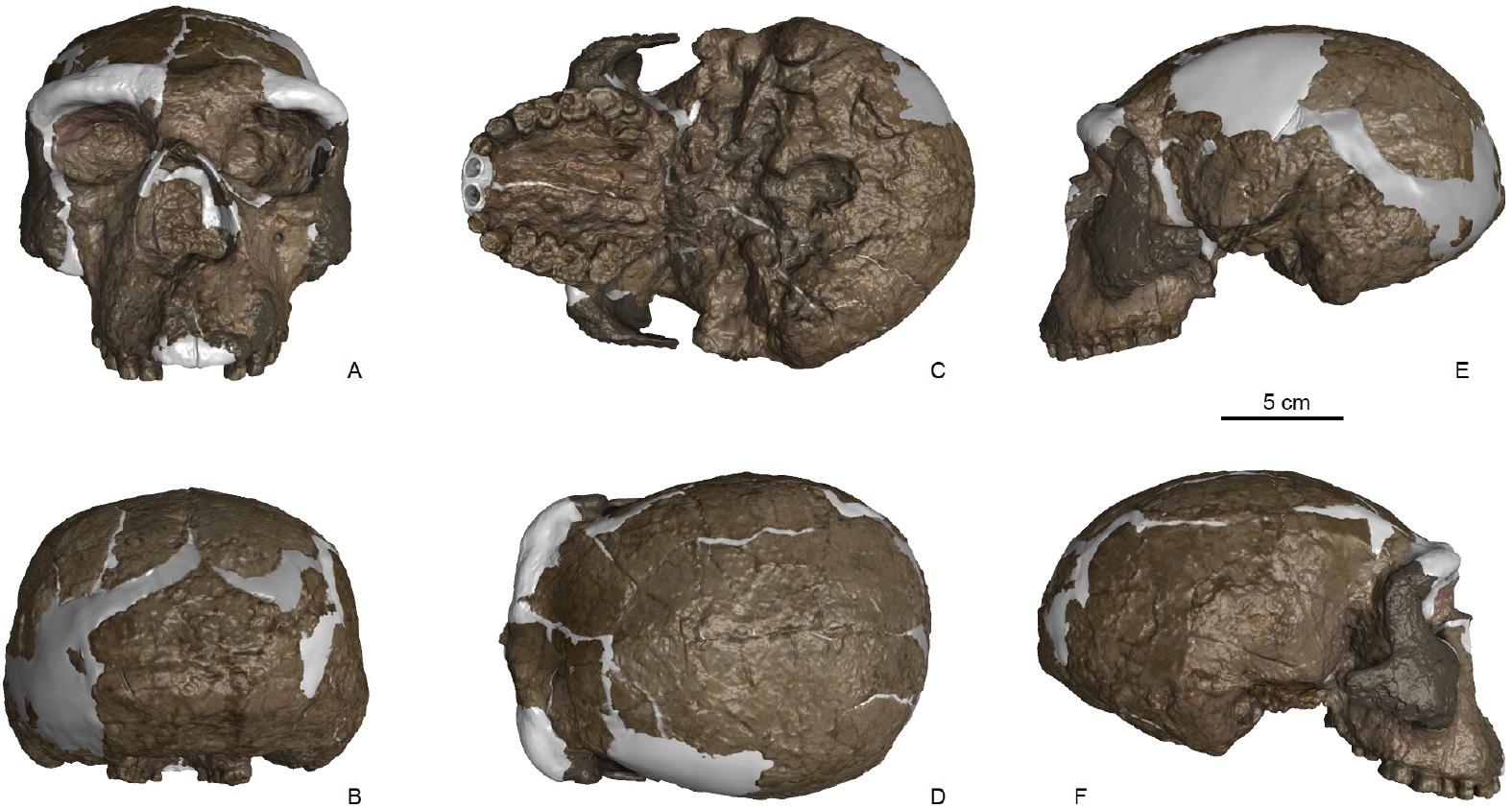
Reconstruction of the Yunxian 2 cranium in standard views. (**A-F**) Anterior, posterior, inferior, superior, left and right views. Brown color indicates the fossil bone. White color indicates the reconstructed parts. Light grey indicates the bones crushed and covered by other bones and matrix. Scale bar indicates 5 cm.

**Fig. 2.**
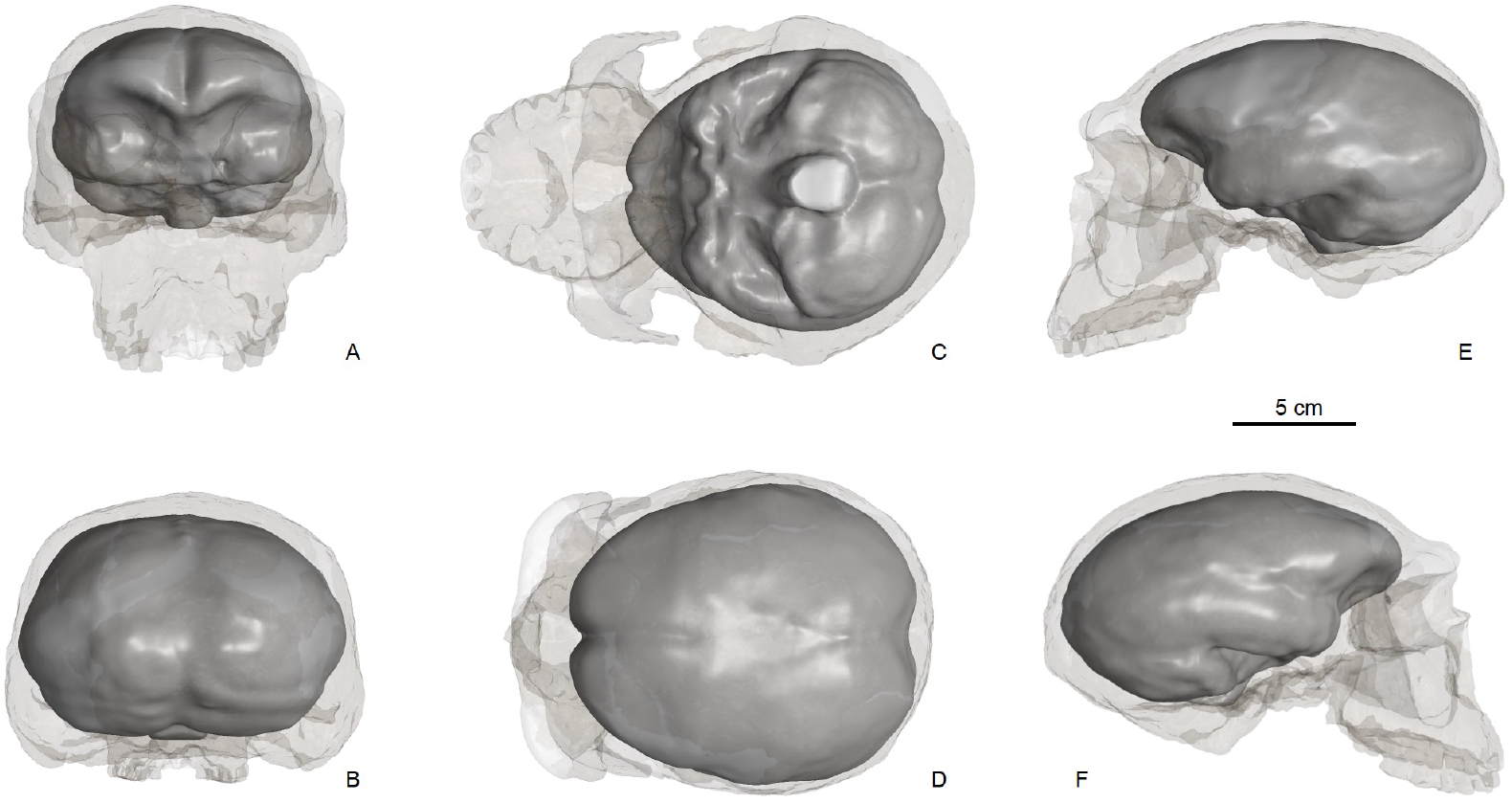
Reconstruction of the endocranial cast of Yunxian 2. (**A-F**) Anterior, posterior, inferior, superior, left and right views. Scale bar indicates 5 cm.

**Fig. 3.**
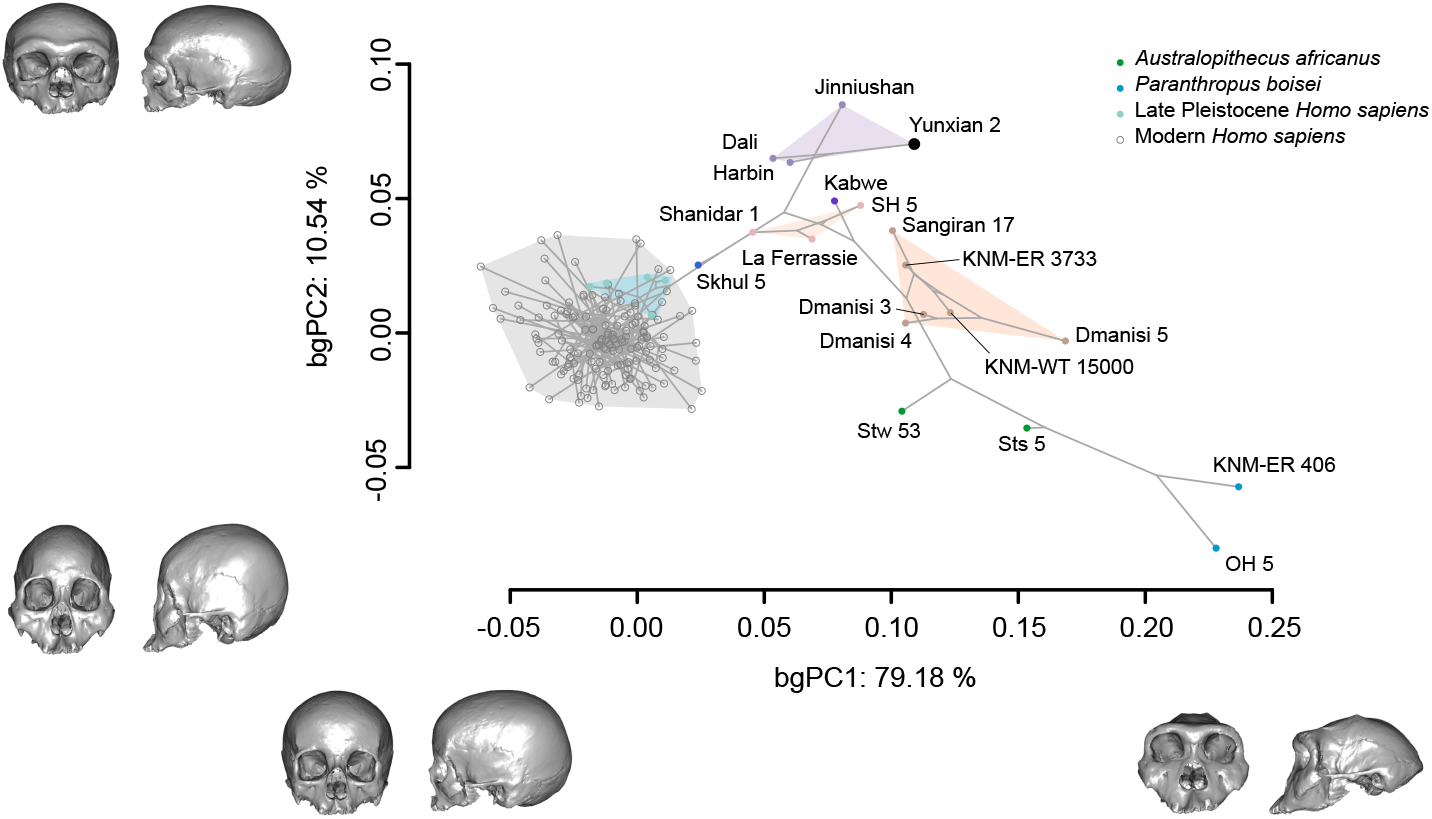
Between-group principal component analysis (bgPCA) of the Procrustes superimposed 276 landmarks and semilandmarks for 177 fossil and recent *Homo* specimens. The first two bgPCs are shown, with the reconstruction of Yunxian 2 projected onto the morphospace. The anterior and lateral views of the crania shown at the bottom and left of the bgPC axis represent the shape extremes of bgPC1 and bgPC2. These shape extremes were created based on the mean shape (specimen YNO227) of all specimens analyzed (see Supplementary Materials for details). Grey lines show the phylogenetic relationships between fossil and recent *Homo* specimens. The relationship between fossils is based on the phylogenetic analyses in this study. The relationship between recent specimens is a random tree and based on the assumption that the recent population is monophyletic.

**Fig. 4.**
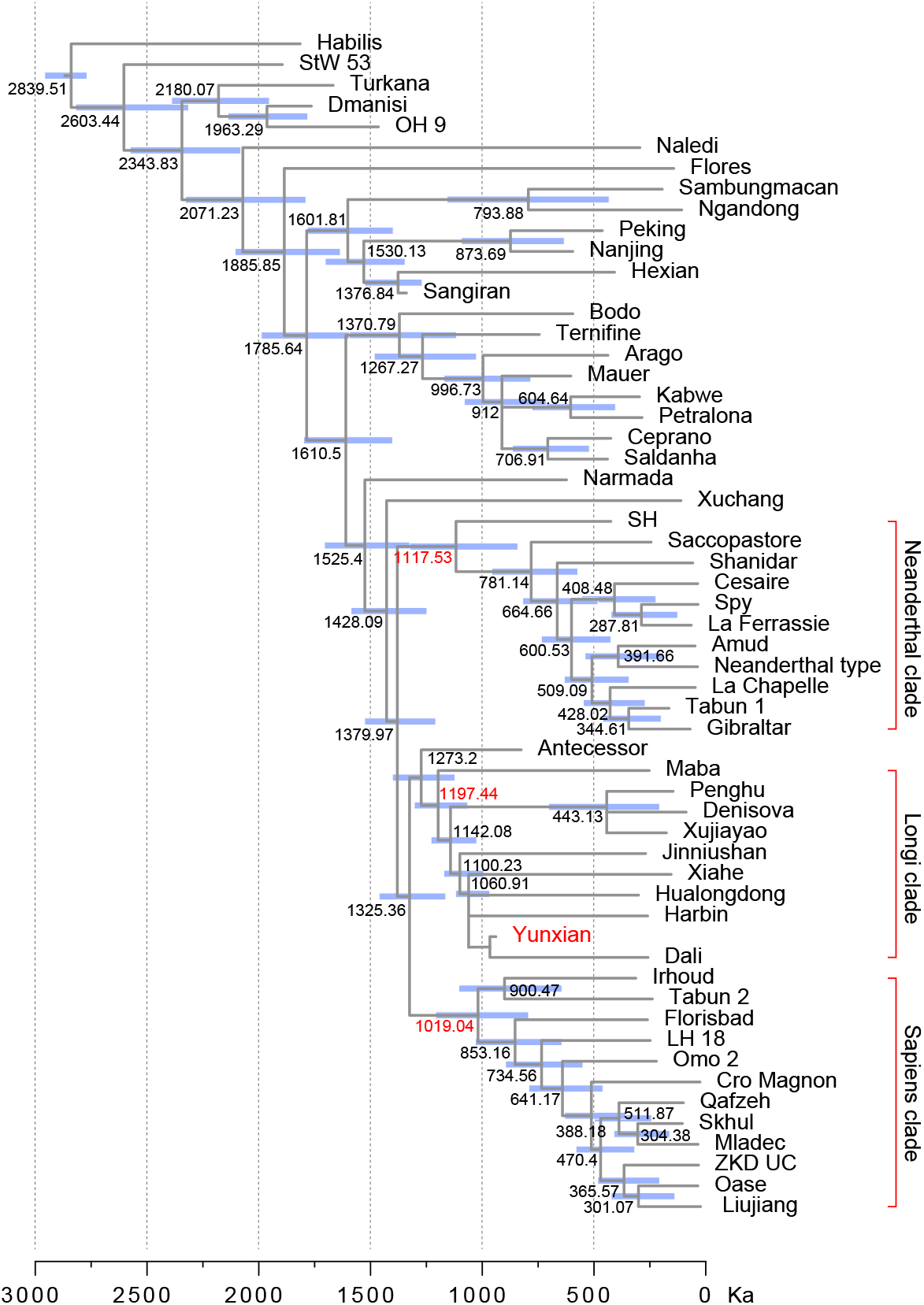
Phylogeny and divergence time of the 57 selected fossil OTUs from the genus *Homo*. The topology of the tree was the majority consensus of the most parsimonious trees from the parsimony analysis in TNT (*31*). The divergence time was inferred from the Bayesian tip-dating analysis in MrBayes 3.2 (*32*). Branch lengths are proportional to the division age in thousand years. Numbers at the internal nodes are the median ages, and the blue bars indicate the 95% highest posterior density interval of the node ages.

Overall, the Yunxian cranium shows a special combination of traits. The reconstructed Yunxian 2 specimen provides new anatomical details for systematic comparison, inferring phylogenetic position, and testing phylogenetic models for the genus *Homo* in general. Our phylogenetic analyses are based on an updated data matrix containing 521 discrete and continuous characters and 57 *Homo* OTUs, which are at the population level by combining multiple specimens from the same locality with the same age range into a paleodeme. To assess the robustness of the phylogenetic inferences based on our restoration and reconstruction of Yunxian 2, we used the bootstrap method to resample its character scoring (see details in Supplementary Materials). The results suggest that potential errors in our discrete and metric character scoring have very little influence on the phylogenetic inferences we have made.

Phylogenetic analyses using the parsimony criterion and Bayesian tip-dating to estimate the divergence dates show that *H. sapiens* Neanderthals, Asian *H. erectus* and Chibanian hominins from Africa and Europe (such as Bodo, Kabwe, Mauer and Arago) are all monophyletic groups. Most Asian Chibanian hominins, often previously referred to as “archaic *Homo sapiens*” including Dali, Jinniushan, Xujiayao, Hualongdong, are grouped together with the Xiahe and Penghu mandibles to form a monophyletic group, the longi clade for convenience. Recently, Xujiayao, Xuchang, Xiahe, Penghu and Denisovans were proposed as a new species of *Homo* (*19*). Our phylogenetic analysis shows that these *Homo* samples (except for Xuchang) belong to the monophyletic longi clade. Yunxian has the oldest age within the longi clade, but is not the most basal fossil in that clade.

The Denisovan Cave in the Altai Mountains has yielded fragmentary fossil humans that have been genetically identified as representing a distinct clade from those of *H. sapiens* and Neanderthals (*20, 21*). Analyses of mitochondrial DNA place Denisovans with the Sima de los Huesos fossils, and outside the divergence between *H. sapiens* and Neanderthals (*22-24*), while nuclear genome sequences suggest that Denisovans are a sister group to Neanderthals (*24, 25*). With only 3 clades (sapiens, Neanderthals and Denisovans), both possible phylogenetic relationships, as reflected in the form of a tree topology, are equally logical and depend on the choice of the rooting point of the tree, and it is also possible that analyses of separately inherited mitochondrial and autosomal DNA will give different results. Our parsimony analysis, based on the limited number of informative characters scored for the Denisovans, suggests that they most likely belong to the longi clade.

The longi clade shared 9 synapomorphies (S-Table 5), including characters such as larger cranial capacity, lower and longer frontal squama, narrower interorbital breadth, deeper glabellar inflexion etc. These characters are evident in the reconstructed morphology of Yunxian 2. The monophyly of the longi + *antecessor* and *sapiens* clade is also well supported. The synapomorphies of this larger group include 31 characters, such as narrower supraorbital torus breadth, narrower interorbital breadth, smaller palate, smaller malar height, presence of maxillary flexion, coronally oriented infraorbital plate, and gracile phalanx without ungual tuberosity expansion.

The Denisovans are known from very few specimens. The apomorphies shared with other fossil humans of the longi clade and shown in the preserved specimens include reduction of the metacone of the Upper M3, strengthening the Carabelli’s cusp of M2, reduction of the M2 width, increases of M2 hypocone and cusp 5, and development of M3 crista obliqua. Morphological study of the Denisovan phalanx indicated that its dimensions and shape are within the range of *H. sapiens* (*26*). A gracile phalanx without expansion of the ungual tuberosity is also present in Jinniushan, a member of the longi clade.

Previous estimates of the divergence time between Neanderthals and *H. sapiens* are about 0.5-0.7 Ma (*27-29*). However, a recent study based on the tree sequence of a large number of ancient genomes with 3589 samples to constrain the dating relationships revealed a much deeper ancestry in sapiens (*30*). The oldest ancestral haplotypes are about 2 Ma and geographically located in Africa. Our Bayesian tip-dating analysis also revealed that the diversification of *Homo* and the origin of *H. sapiens* have much greater time depths. The origin of the longi clade can be inferred to be about 1.2 Ma, probably slightly older than the Yunxian fossils. The origin of the sapiens clade is estimated to be about 1.02 Ma, also close to the age of Yunxian. The divergence between the longi clade and the sapiens clade is at about 1.32 Ma. The monophyletic Neanderthal clade, widely thought to be sister to *H. sapiens*, diverged from the longi and sapiens clades at about 1.38 Ma in our analysis.

Given its geological age of 0.94-1.10 Ma (*5*), Yunxian is close to the theoretical origin time of the clades of longi and sapiens. Our reconstruction of Yunxian 2 shows that this fossil human has mosaic features of plesiomorphies, as seen in *H. erectus*, and apomorphies, as seen in longi and sapiens. It is reasonable to conclude that Yunxian is morphologically and chronologically close to the last common ancestor of the longi and sapiens clades.

## Acknowledgments

We thank Drs. Hou and Ma for help with CT scanning and surface modeling. Mr. Zhang helped with cluster computing. We thank Drs. Venditti, White, and Puschel for helpful discussions and reexamination of the Bayesian analysis. Images and CT scans of specimens from the National Museum of Kenya (NMK) are courtesy of NMK and human-fossil-record.org. CT scans of Steinheim were provided by the State Museum of Natural History Stuttgart and the Department of Human Evolution, Max Planck Institute for Evolutionary Anthropology. We thank Drs. Manzi, Buzi, Di Vincenzo, Profico and their team members for providing digital reconstructions of Steinheim and Ceprano. We thank Anthropology staff at the Natural History Museum London, for the provision of comparative material.

## Funding

National Natural Science Foundation of China No. 41988101 (XN)

National Key Research and Development Program of China No. 2023YFF0804502 (XN, QL, CZ)

Key Project of Beijing Social Science Foundation No.22LSA022 (XF)

The research of CS is supported by the Calleva Foundation No. SDV17014 and the Human Origins Research Fund

## Author contributions

Conceptualization: XN, CS

Methodology: XN, DL, QY, CZ, CS

Investigation: XF, DL, QY, FG, QL, CZ, CS, XN

Visualization: DL, QY, XN

Funding acquisition: XN

Project administration: XN

Supervision: XN

Writing – original draft: XN, CS

Writing – review & editing: XN, CS, XF, DL, FG, QY, QF, YF, XH, CT, HZ, QL, CZ

## Data and materials availability

All data are available in the main text or the supplementary materials.

## Supplementary Materials

Materials and Methods

Supplementary Text

Figures S1 to S30

Tables S1 to S7

References (*33–129*)

Appendix 1-7

