## Supplementary materials for "The phylogenetic position of the Yunxian cranium elucidates the origin of *Homo longi* and the Denisovans"

Science
AAAS

### Supplementary Materials for

#### The phylogenetic position of the Yunxian cranium elucidates the origin of *Homo longi* and the Denisovans

Xiaobo Feng[1,2,3,4]†, Qiyu Yin[5,6]†, Feng Gao[7]†, Dan Lu[5,6]†, Qin Fang[8], Yilu Feng[9], Xuchu Huang[10], Chen Tan[11], Hanwen Zhou[12], Qiang Li[5,6], Chi Zhang[5,6], Chris Stringer[13]*, Xijun Ni[5,6,14]*

†Co-first authors

**Affiliations:**

1. School of History and Culture, Shanxi University, Taiyuan, 030006, China
2. Hubei Polytechnic University, Huangshi, 435003, China
3. Institute of Yunxian Man Site, Hanjiang Normal University, Shiyan, 442000, China
4. School of History Culture and Tourism, Hanjiang Normal University, Shiyan, 442000, China
5. Institute of Vertebrate Paleontology and Paleoanthropology, Chinese Academy of Sciences, Beijing, 100044, China
6. College of Earth and Planetary Sciences, University of Chinese Academy of Sciences, Beijing, 100049, China
7. Yunnan Institute of Cultural Relics and Archeology, Kunming 650118, China
8. Hubei Provincial Institute of Cultural Relics and Archaeology, Wuhan, 430077, China
9. School of History of Wuhan University, Wuhan, 430072, China
10. Museum of Shiyan, Shiyan, 442000, China
11. Poly Art Research Institute, Beijing, 100010, China
12. School of Earth Sciences, China University of Geosciences, Wuhan 430074, China
13. Centre for Human Evolution Research, Natural History Museum, London, UK
14. PaleoAnthropological Research Center, Fudan University, Shanghai, 200433, China

**The PDF file includes:**

Materials and Methods
Supplementary Text
Figures S1 to S30
Tables S1 to S7
References 1-129

**Other Supplementary Materials for this manuscript include the following:**

Appendix 1 to 7

**Discovery backgrounds** 7
S-Figure 1. The Yunxian Site 7
S-Figure 2. Aerial photography of the Yunxian Site 8
**Reconstruction of Yunxian 2** 8
**Surface model split and reinstatement** 10
S-Figure 4. Surface model of Yunxian 2 in standard views. Different colors indicate segments that were digitally sectioned along cracks. Segments without plastic deformation were used for reinstatement and reconstruction. A. anterior view; B. posterior view; C. inferior view; D. superior view; E. left view; F. right view. 10
S-Figure 5. Comparison of unreconstructed and reconstructed Yunxian 2 in anterior views. A. Deformed skull without reinstatement and reconstruction. B, C. reinstated and reconstructed skull. In A and B, different colors indicate segments that were digitally sectioned along cracks. Segments without plastic deformation were used for reinstatement and reconstruction. C. Reconstructed skull with natural fossil color. In B and C, white indicates the reconstructed parts inferred from the fracture edge and Yunxian 1; light gray indicates the missing parts and severe plastic deformation; medium gray indicates the bone crushed into the cranial cavity and covered by other bones; dark gray indicates the parts reconstructed from Yunxian 1. 11
S-Figure 6. Comparison of unreconstructed and reconstructed Yunxian 2 in posterior views. A. Deformed skull without reinstatement and reconstruction. B, C. reinstated and reconstructed skull. In A and B, different colors indicate segments that were digitally sectioned along cracks. Segments without plastic deformation were used for reinstatement and reconstruction. C. Reconstructed skull with natural fossil color. In B and C, white indicates the reconstructed parts inferred from the fracture edge and Yunxian 1; light gray indicates the missing parts and severe plastic deformation; medium gray indicates the bone crushed into the cranial cavity and covered by other bones; dark gray indicates the parts reconstructed from Yunxian 1. 12
S-Figure 8. Comparison of unreconstructed and reconstructed Yunxian 2 in superior views. A. Deformed skull without reinstatement and reconstruction. B, C. reinstated and reconstructed skull. In A and B, different colors indicate segments that were digitally sectioned along cracks. Segments without plastic deformation were used for reinstatement and reconstruction. C. Reconstructed skull with natural fossil color. In B and C, white indicates the reconstructed parts inferred from the fracture edge and Yunxian 1; light gray indicates the missing parts and severe plastic deformation; medium gray indicates the bone crushed into the cranial cavity and covered by other bones; dark gray indicates the parts reconstructed from Yunxian 1. 12
S-Figure 9. Comparison of unreconstructed and reconstructed Yunxian 2 in left views. A. Deformed skull without reinstatement and reconstruction. B, C. reinstated and reconstructed skull. In A and B, different colors indicate segments that were digitally sectioned along cracks. Segments without plastic deformation were used for reinstatement and reconstruction. C. Reconstructed skull with natural fossil color. In B and C, white indicates the reconstructed parts inferred from the fracture edge and Yunxian 1; light gray indicates the missing parts and severe plastic deformation; medium gray indicates the bone crushed into the cranial cavity and covered by other bones; dark gray indicates the parts reconstructed from Yunxian 1. 13
S-Figure 10. Comparison of unreconstructed and reconstructed Yunxian 2 in right views. A. Deformed skull without reinstatement and reconstruction. B, C. reinstated and reconstructed skull. In A and B, different colors indicate segments that were digitally sectioned along cracks. Segments

without plastic deformation were used for reinstatement and reconstruction. C. Reconstructed skull with natural fossil color. In B and C, white indicates the reconstructed parts inferred from the fracture edge and Yunxian 1; light gray indicates the missing parts and severe plastic deformation; medium gray indicates the bone crushed into the cranial cavity and covered by other bones; dark gray indicates the parts reconstructed from Yunxian 1. 13

**Removing cracking 13**

S-Figure 11. CT images of Yunxian 2 and 3D reconstruction showing cracks. A. Transverse CT slice through the posterior part of the skull. B. 3D reconstruction of the occipital region, the cracks have been highlighted and enhanced. 14

**Evaluating the variation of the reconstructions 14**

S-Table 1. List of anatomical landmarks 15

S-Table 2. List of curves 16

S-Figure 12. landmarks on the presented specimen (modern *H. sapiens* specimen curated in IVPP). Left: anterior view; middle: lateral view (left); right: inferior view. Red dots: fixed landmarks; black dots: curves; blue dots: patch landmarks. 17

S-Figure 13. Between-group principal component analysis (bgPCA) of the Procrustes superimposed landmarks for all specimens used in this study. The first two bgPCs are shown, with the seven Yunxian 2 reconstruction versions projected onto the morphospace. Variation of the reconstruction of Yunxian falls in a small range when compared with other *Homo* fossils and extant human skulls. 18

S-Figure 14. PCA on landmark configurations of the seven versions of Yunxian 2 reconstructions. Colored meshes illustrate the shape variations corresponding to the extreme values of each principal component. The Black dot indicates the preferred final version. 19

S-Figure 15. Local morphological differences between the first six versions and the final version of the reconstruction of Yunxian 2, implemented by the function 'meshDist' of R. Red areas indicate high variations and blue areas indicate low variations. a to f stands for the six versions of the reconstruction. In anterior view. 19

S-Figure 16. Local morphological differences between the first six versions and the final version of the reconstruction of Yunxian 2, implemented by the function 'meshDist' of R. Red areas indicate high variations and blue areas indicate low variations. a to f stands for the six versions of the reconstruction. In ventral view. 20

S-Figure 17. Local morphological differences between the first six versions and the final version of the reconstruction of Yunxian 2, implemented by the function 'meshDist' of R. Red areas indicate high variations and blue areas indicate low variations. a to f stands for the six versions of the reconstruction. In left lateral view. 20

S-Figure 18. Local morphological differences between the first six versions and the final version of the reconstruction of Yunxian 2, implemented by the function 'meshDist' of R. Red areas indicate high variations and blue areas indicate low variations. a to f stands for the six versions of the reconstruction. In posterior view. 21

S-Figure 19. Local morphological differences between the first six versions and the final version of the reconstruction of Yunxian 2, implemented by the function 'meshDist' of R. Red areas indicate high variations and blue areas indicate low variations. a to f stands for the six versions of the reconstruction. In right lateral view. 22

S-Figure 20. Local morphological differences between the first six versions and the final version of the reconstruction of Yunxian 2, implemented by the function 'meshDist' of R. Red areas indicate high variations and blue areas indicate low variations. a to f stands for the six versions of the reconstruction. In superior view. 22

**Reconstruction of the endocranial cast** **22**

**Morphology of the reconstructed Yunxian 2** **23**

**Comparative morphology of the Yunxian 2 cranium** **24**

S-Table 3. Measurements of the Yunxian 2 cranium and comparisons with other Middle-Late Pleistocene *Homo* cranial fossils. Linear measurements in millimeters, angles in degrees, ratios ranged from 0 to 100. 26

**Taxonomic scope for comparison and phylogenetic analyses** **34**

S-Table 4. Specimens used for comparison and phylogenetic analysis 34

**Morphological data collection** **39**

**Characters for phylogenetic analysis** **40**

**Test of potential correlations among continuous characters** **41**

S-Figure 21. Non-parameter Kendall correlation analysis among 267 continuous characters. Each color dot represents a pairwise Kendall Tau. 42

**Parsimony analysis** **42**

S-Figure 22. The preferred majority rule consensus tree of the 42 most parsimonious trees. Linear measurements of cranium and upper dentition were transformed into ratios by dividing the measurements by the 1/3 power of the cranial capacity. Numbers near internal nodes indicate majority consensus percentage. 43

S-Figure 23. Majority rule consensus tree of the 24 most parsimonious trees including Eliye Springs, Ndutu, Steinheim and Rabat. Linear measurements of cranium and upper dentition were transformed into ratios by dividing the measurements by the 1/3 power of the cranial capacity. Numbers near internal nodes indicate majority consensus percentage. The main topological feature of this tree is identical to the tree excluding the 4 fossils (as in S-figure 22). 44

S-Figure 24. Majority rule consensus tree of the 31 most parsimonious trees. Linear measurements of cranium and upper dentition were transformed into ratios by dividing the measurements by the maximum frontal width. Numbers near internal nodes indicate majority consensus percentage. The main topological feature of this tree is identical to the tree based on the linear measurements of cranium and upper dentitions were transformed into ratios by dividing the measurements by the 1/3 power of the cranial capacity (as in S-figure 22). 45

**Test of influence of potential correlations among characters on the phylogeny** **45**

S-Figure 25. Symmetric resampling analyses with different resampling probability. Resampling trees were mapped on to the majority consensus of the most parsimonious trees inferred from the original data matrix. Supports of different resampling strength are mainly positive. 47

S-Figure 26. Symmetric resampling analyses with different resampling probability. Resampling trees were mapped on to the majority consensus of the most parsimonious trees inferred from the original data matrix. The distribution of supports of different resampling strength shows a similar pattern. 48

S-Figure 27. Random duplication resampling analysis shows that major clades can be identified even with very high resampling strength. The topology of the phylogenetic tree is almost identical to the most parsimonious tree for 5% ~ 30% character duplication. 49

S-Figure 28. The preferred majority rule consensus tree of the 42 most parsimonious trees (as in S-Figure 22). Numbers before the slashes are the TBE (*118*) of 1 million trees of the bootstrap analysis (*100*). Numbers after slashes are the Bremer supports (*120*). Number in red color indicates that the bootstrap support TBE is greater than 70% and the node is strongly supported (*118, 119*). 50

**Effect of the equally-weighted character subgroups on phylogeny 50**

S-Figure 29. Majority rule consensus tree of the 41 most parsimonious trees, with neurocranial, visceral cranial, mandibular, dental, and postcranial character subgroups being equally weighted. The main topological feature of this tree is identical to the most parsimonious tree without reweighted the character subgroups (as in S-figure 22). The numbers near the internal nodes are the percentages of consensus. 51

**Bootstrap analysis of the phylogenetic position of Yunxian 2 51**

S-Figure 30. Bootstrap resampling analysis by randomly removing 50% of the total metric and discrete scores of Yunxian 2. Bootstrap trees were summarized on the preferred majority consensus tree of the most parsimonious trees (as in S-Figure 22). Numbers at the internal nodes are the TBE (*118*) of 1477 trees of the bootstrap analysis (*100*). 53

**Bayesian tip-dating analyses 53**

**Synapomorphies inferred from parsimony analysis 55**

S-Table 5. Nine synapomorphies shared by the *longi* clade (not including *Homo antecessor*) 55

S-Table 6. Forty-four synapomorphies shared by the sapiens clade 55

S-Table 7. Forty-eight synapomorphies shared by the Neanderthal clade 57

**Reference 59**

**Discovery backgrounds**

The Yunxian Site (S-Figure 1-2), also known as the Xuetangliangzi Site, is located on the left bank of the Han River (a tributary of the Yangtze River) in northern Hubei Province in central China. It is about 550 km northwest of Wuhan City and about 40 km west of Yunxian City. Several major excavation projects were carried out at the site in 1990, 1991, 1995, 1998, 1999, 2006 and 2021. Three *Homo* crania (Yunxian 1, Yunxian 2 and Yunxian 3) were discovered in Layer C3 at the Yunxian Site. Yunxian 1 was discovered in May 1989, Yunxian 2 was discovered in June 1990, and Yunxian 3 was discovered in 2022.

The Pleistocene sediments at the Yunxian site can be divided into 6 layers.

Layer C1: Surface soil, about 0.40 m thick.

Layer C2: Reddish brown clay and silts, about 4.50 m thick. This layer contains some stone tools but no fossils.

Layer C3: Yellowish brown sandy-clay, about 2 m thick. The layer is very fossiliferous, with many mammalian fossils. Yunxian 1, 2 and 3, and stone tools were discovered in this layer.

Layer C4: Medium and fine sandstones, about 1.10m thick. Clay pellets and some concretions of iron oxide and manganese are present. Stone tools and mammal bones are rare.

Layer C5: Medium and fine pinkish sands, 0.40m thick. A few millimeter grains of manganese oxide.

Layer C6: Gravel and river sands containing large pebbles and medium and fine sand lenses, 3 to 5m thick.

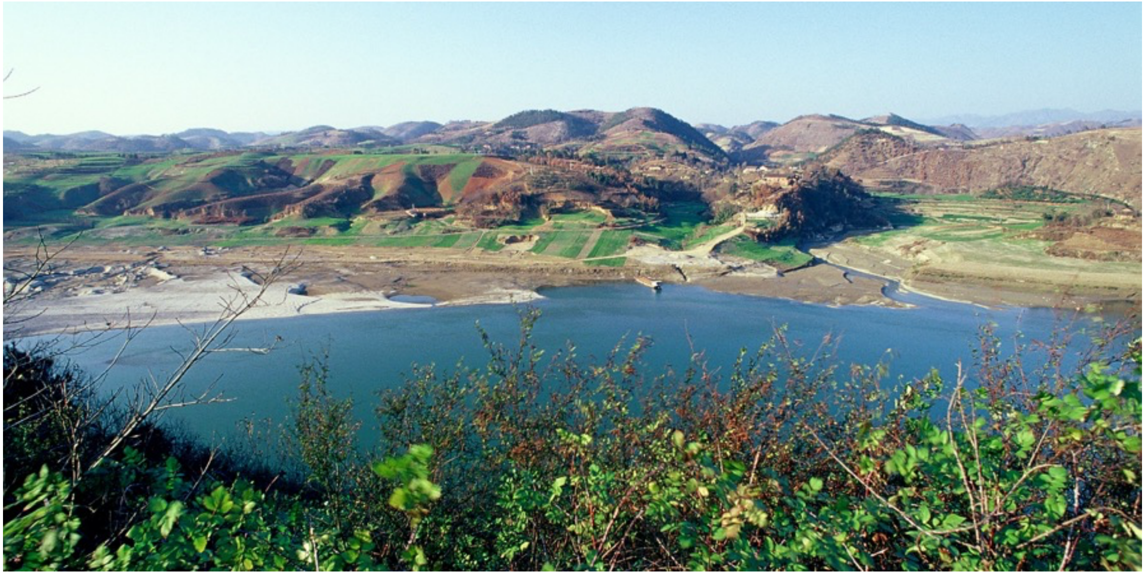

S-Figure 1. The Yunxian Site

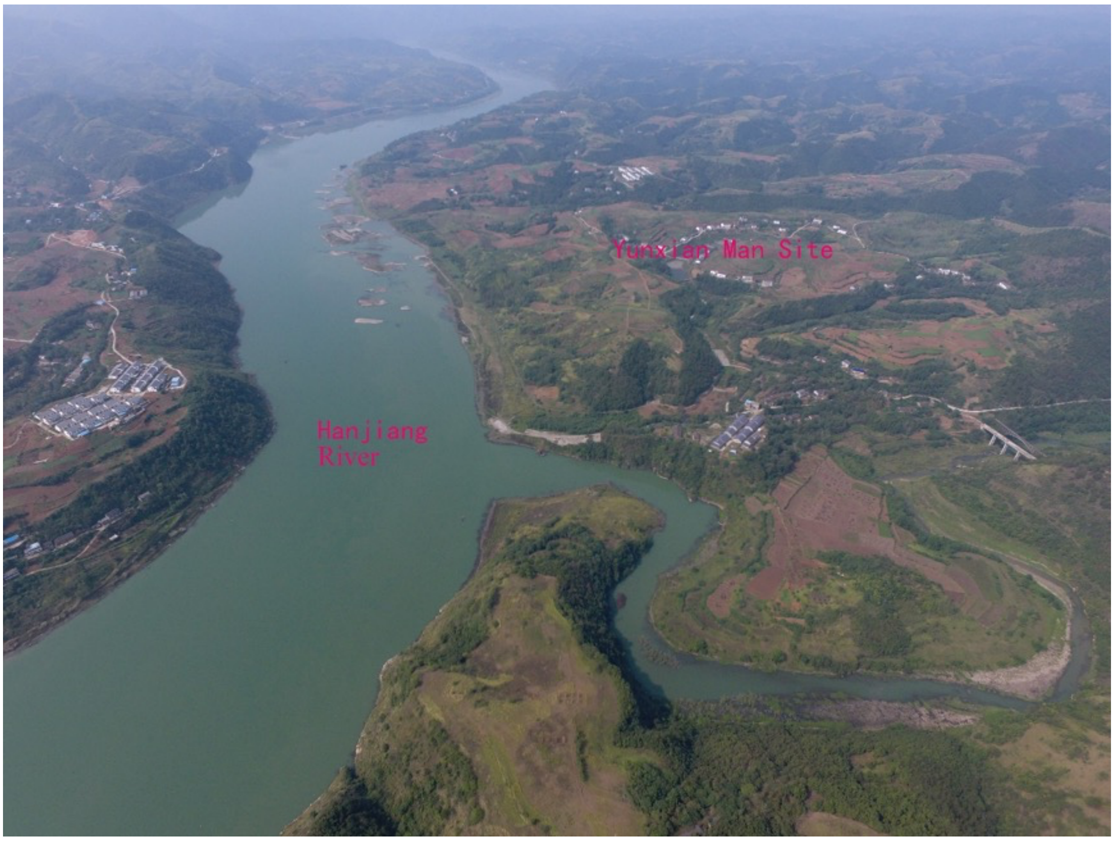


S-Figure 2. Aerial photography of the Yunxian Site

**Reconstruction of Yunxian 2**

We used the procedure depicted in S-Figure 3 to make the reconstruction.

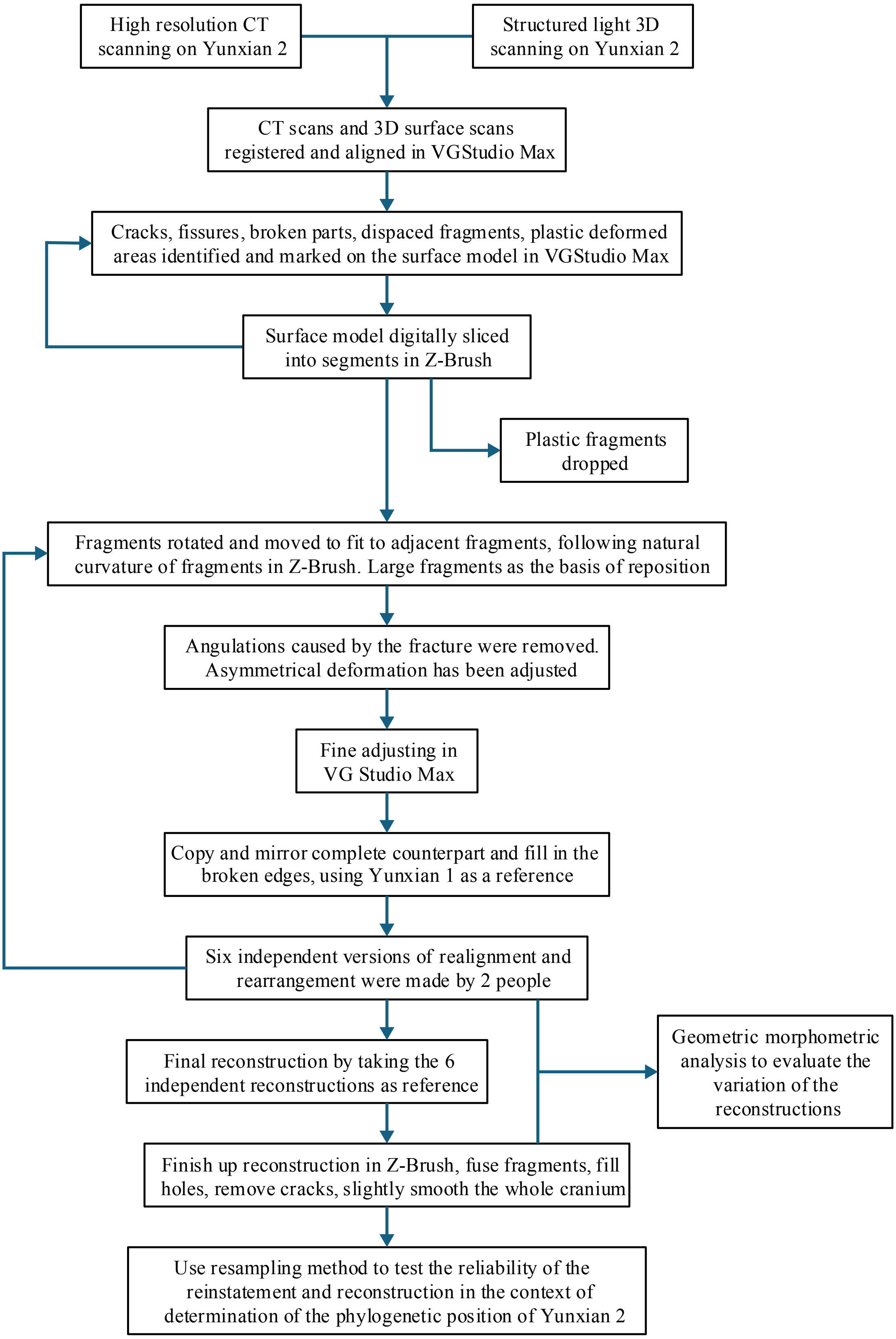


S-Figure 3. Procedure for reconstructing Yunxian 2.

Surface model split and reinstatement

Yunxian 1 and 2 were CT scanned using the High Resolution CT facilities at the Key Laboratory of Vertebrate Evolution and Human Origins of the Chinese Academy of Sciences. The resolution is 62.5 micrometers. X-ray CT images were exported to the VG Studio Max 3.2 to build thre dimensional models. Yunxian 1 and 2 were then surface-scanned by using the Arctec Space Spider surface scanner with a 3D point accuracy up to 50 micrometers.

The surface models and CT data were registered together in VG Studio Max 3.2. From the CT scans, cracks of fragmentation and displacement were identified and marked on the surface model, and registered on the CT data. Several regions of plastic deformation were also identified and marked. Along the cracks, the surface model was digitally sliced into 99 segments (S-Figure 4). Segments without plastic deformation were preserved for reinstatement and reconstruction. The preserved segments were then exported to the Z-Brush software for realignment and rearrangement. Each fragment was rotated and moved slightly to fit it to adjacent fragments. The natural curvature of the fragments was made to fit as smoothly as possible, and all angulations caused by the fracture were removed. Asymmetrical deformation has been adjusted. The large fragments, such as a part of frontal preserving part of the supraorbital ridges, the left and right maxillae, left and right ear region of the temporals and the basioccipital, allowed relatively reliable digital repositioning of these bones (S-Figure 5-10).

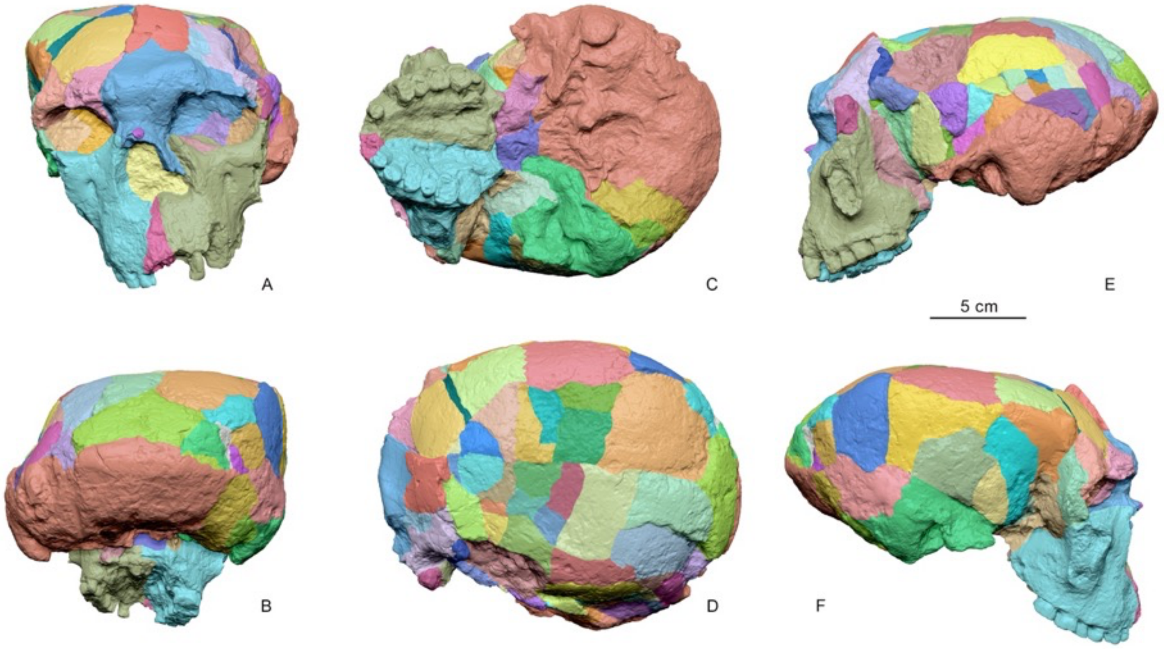


S-Figure 4. Surface model of Yunxian 2 in standard views. Different colors indicate segments that were digitally sectioned along cracks. Segments without plastic deformation were used for reinstatement and reconstruction. A. anterior view; B. posterior view; C. inferior view; D. superior view; E. left view; F. right view.

A small part of the left maxilla is broken. We copied and made a mirror of the complete right maxilla and composited the small piece to the broken left maxilla. The left and right zygomatic bones are broken. We copied the two bones from Yunxian 1 and did some minor rescaling to fit them to the broken edges of the left and right zygomatics of Yunxian 2. The supraorbital ridges of Yunxian 2 are also broken. The size of the broken edges suggests that they are quite thick. The supraorbital ridges of Yunxian 1 are completely preserved, but they are significantly thinner than those of Yunxian 2, as shown by the broken edges of Yunxian 2. We digitally reconstructed the supraorbital ridges of Yunxian 2 by filling in the broken edges and using Yunxian 1 as a reference (S-Figure 5-10).

Realignment and rearrangement of the undeformed fragments, and reconstruction of the broken parts were made 6 times by 2 people. Each person did this 3 times independently. Lastly, the 6 reinstatements and reconstructions of Yunxian 2 were fitted together to form a reference model, then the final reconstruction was made by taking this model as the key reference (S-Figure 5-10).

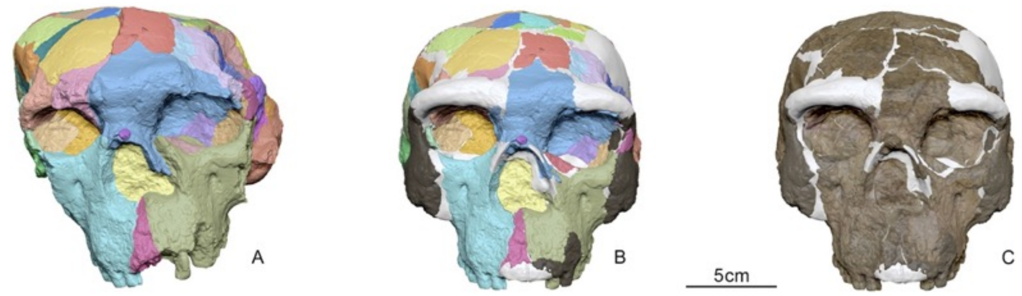


S-Figure 5. Comparison of unreconstructed and reconstructed Yunxian 2 in anterior views. A. Deformed skull without reinstatement and reconstruction. B, C. reinstated and reconstructed skull. In A and B, different colors indicate segments that were digitally sectioned along cracks. Segments without plastic deformation were used for reinstatement and reconstruction. C. Reconstructed skull with natural fossil color. In B and C, white indicates the reconstructed parts inferred from the fracture edge and Yunxian 1; light gray indicates the missing parts and severe plastic deformation; medium gray indicates the bone crushed into the cranial cavity and covered by other bones; dark gray indicates the parts reconstructed from Yunxian 1.

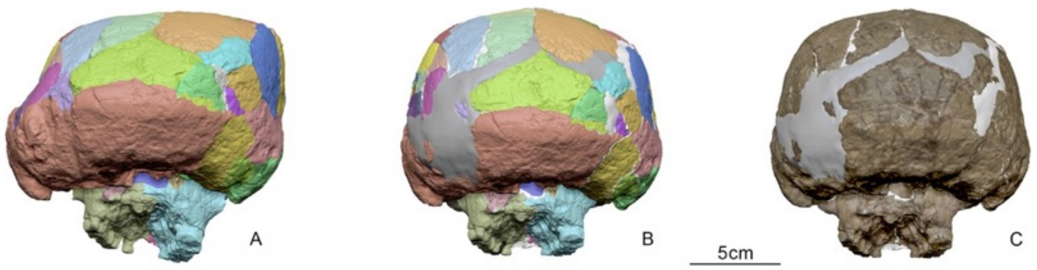

S-Figure 6. Comparison of unreconstructed and reconstructed Yunxian 2 in posterior views. A. Deformed skull without reinstatement and reconstruction. B, C. reinstated and reconstructed skull. In A and B, different colors indicate segments that were digitally sectioned along cracks. Segments without plastic deformation were used for reinstatement and reconstruction. C. Reconstructed skull with natural fossil color. In B and C, white indicates the reconstructed parts inferred from the fracture edge and Yunxian 1; light gray indicates the missing parts and severe plastic deformation; medium gray indicates the bone crushed into the cranial cavity and covered by other bones; dark gray indicates the parts reconstructed from Yunxian 1.

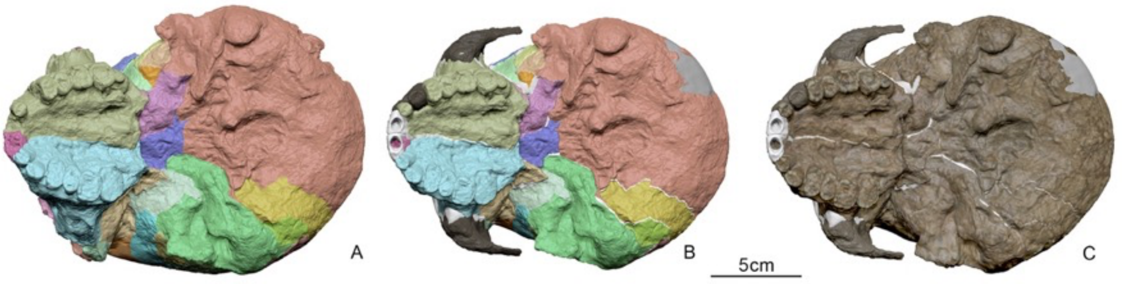


S-Figure 7. Comparison of unreconstructed and reconstructed Yunxian 2 in inferior views. A. Deformed skull without reinstatement and reconstruction. B, C. reinstated and reconstructed skull. In A and B, different colors indicate segments that were digitally sectioned along cracks. Segments without plastic deformation were used for reinstatement and reconstruction. C. Reconstructed skull with natural fossil color. In B and C, white indicates the reconstructed parts inferred from the fracture edge and Yunxian 1; light gray indicates the missing parts and severe plastic deformation; medium gray indicates the bone crushed into the cranial cavity and covered by other bones; dark gray indicates the parts reconstructed from Yunxian 1.

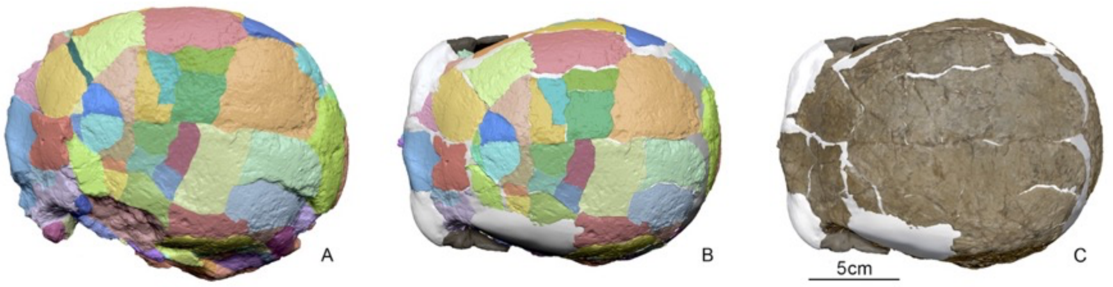


S-Figure 8. Comparison of unreconstructed and reconstructed Yunxian 2 in superior views. A. Deformed skull without reinstatement and reconstruction. B, C. reinstated and reconstructed skull. In A and B, different colors indicate segments that were digitally sectioned along cracks. Segments without plastic deformation were used for reinstatement and reconstruction. C. Reconstructed skull with natural fossil color. In B and C, white indicates the reconstructed parts inferred from the fracture edge and Yunxian 1; light gray indicates the missing parts and severe plastic deformation; medium gray indicates the bone crushed into the cranial cavity and covered by other bones; dark gray indicates the parts reconstructed from Yunxian 1.

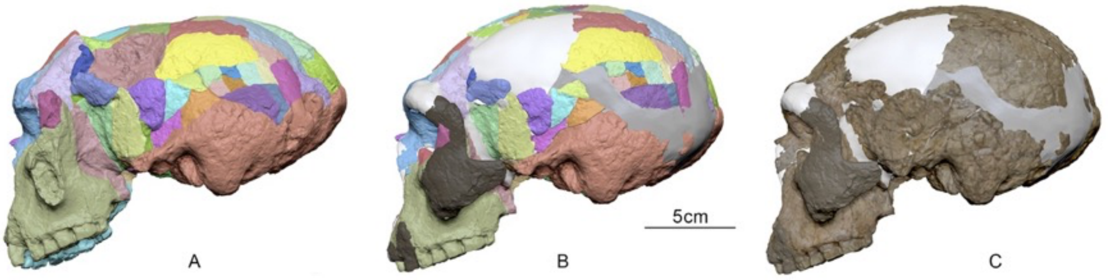


S-Figure 9. Comparison of unreconstructed and reconstructed Yunxian 2 in left views. A. Deformed skull without reinstatement and reconstruction. B, C. reinstated and reconstructed skull. In A and B, different colors indicate segments that were digitally sectioned along cracks. Segments without plastic deformation were used for reinstatement and reconstruction. C. Reconstructed skull with natural fossil color. In B and C, white indicates the reconstructed parts inferred from the fracture edge and Yunxian 1; light gray indicates the missing parts and severe plastic deformation; medium gray indicates the bone crushed into the cranial cavity and covered by other bones; dark gray indicates the parts reconstructed from Yunxian 1.

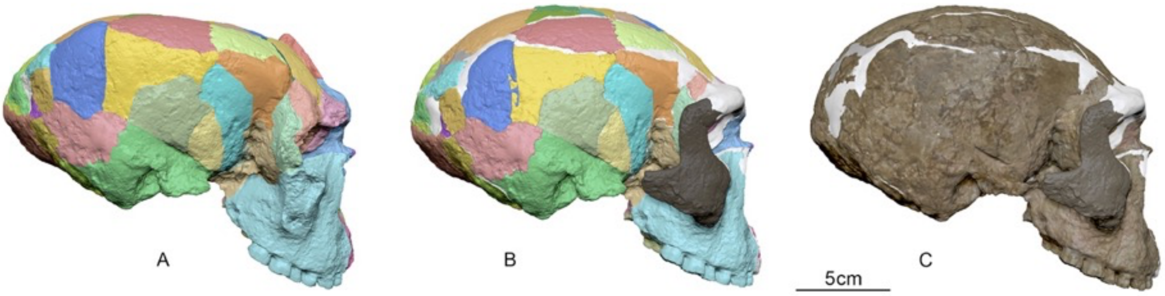


S-Figure 10. Comparison of unreconstructed and reconstructed Yunxian 2 in right views. A. Deformed skull without reinstatement and reconstruction. B, C. reinstated and reconstructed skull. In A and B, different colors indicate segments that were digitally sectioned along cracks. Segments without plastic deformation were used for reinstatement and reconstruction. C. Reconstructed skull with natural fossil color. In B and C, white indicates the reconstructed parts inferred from the fracture edge and Yunxian 1; light gray indicates the missing parts and severe plastic deformation; medium gray indicates the bone crushed into the cranial cavity and covered by other bones; dark gray indicates the parts reconstructed from Yunxian 1.

Removing cracking

Both Yunxian 1 and 2 have numerous small cracks (S-Figure 11). These cracks have enlarged the two skulls. Because the cracks are thin and numerous, slicing, rearranging, and reorienting large fragments as shown in S-Fig. 10 cannot reconstruct the actual size. We took random samples from 3 sections of the CT data and measured the width of the cracks and calculated the proportion of cracks relative to the bone length of each sample. Forty samples were taken from each section. A total of 120 samples were measured. From this ratio of crack width to bone length, we can calculate that on average, the cracks increased the size of the original specimen by 104.38%. Thus, the reinstated and reconstructed cranium was shrunk to 95.8% of the Yunxian 2 fossil.

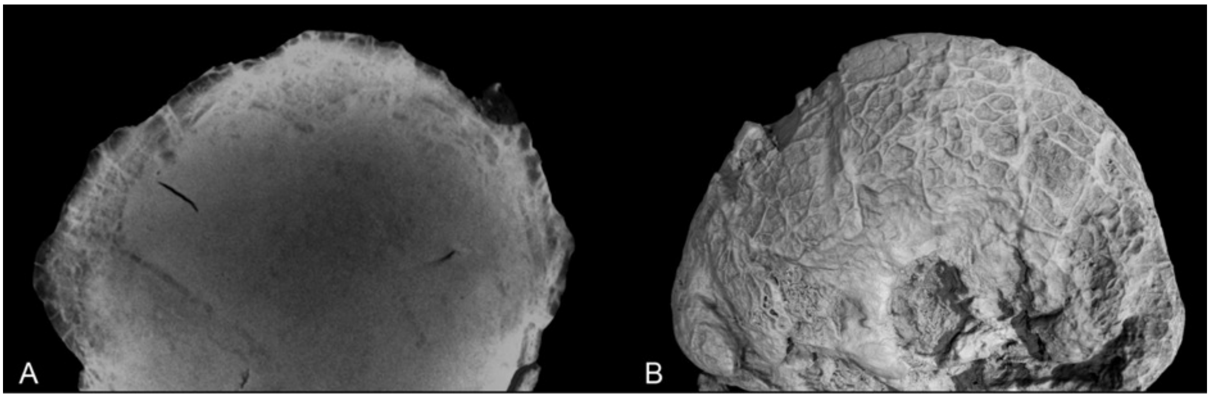


S-Figure 11. CT images of Yunxian 2 and 3D reconstruction showing cracks. A. Transverse CT slice through the posterior part of the skull. B. 3D reconstruction of the occipital region, the cracks have been highlighted and enhanced.

Evaluating the variation of the reconstructions

We used landmark-based morphological geometric analysis to evaluate the variation of the 7 reconstructions. This evaluation method was used by Stefano Benazzi and colleagues in their OH5 and KNM-ER 1813 reconstructions (*10, 11*). 276 landmarks in total were collected from 174 specimens (24 human fossils, 153 recent human specimens, and 7 versions of the Yunxian 2 reconstruction) using Stratovan Checkpoint software. We built a template consisting of 66 anchor points and 9 curves (S-Table 1, 2), with 210 semilandmarks evenly spaced between the landmarks on each curve (S-Figure 12). This template was then applied to new specimens using the Thin-Plate Spline (TPS) method. For surface landmarks, we used 'buildtemplate' function from the 'geomorph' package to generate a template of 300 surface slider semilandmarks based on the 66 landmarks in S-Table 1. This function automatically selected a specified number of semilandmarks from the mesh at approximately equal distances (*33, 34*). We used the 'findMeanSpec' function from the 'geomorph' package to identify YNO.227 as the specimen mesh most closely representing the mean morphology (*35*). As a result, we selected YNO.227 as the reference template for placing landmark points on the patches. Due to poor preservation of areas such as the orbits, zygomatic arches, and sphenoid bones in hominin fossils, we removed these regions from the specimen mesh in Meshmixer before placing the landmarks. This was done to avoid placing semilandmarks on these problematic areas. The 'placepatch' function from the Morpho package in R was then used to project the template onto each specimen, as it has been successfully applied in geometric morphometric studies elsewhere (*36-40*). This function projects surface points from an atlas onto the samples' surface using Thin-Plate Spline deformation, while minimizing distortions. Then the surface semilandmarks obtained after projection were combined with the original 276 fixed and curve landmarks to form a complete landmark set.

The following morphological analyses were performed in R 4.4.0 (*41*). Curve and surface semilandmarks were slid to minimize bending energy using the 'slider3d' function in the Morpho package using TPS (*40*). This procedure ensures that semilandmarks best match the positions of corresponding points in a reference configuration (*42, 43*). After sliding, all landmarks and semilandmark datasets were converted to shape variables, and factors like location, size, and orientation were removed using Generalized Procrustes Analysis

(GPA) method. This step was performed using the ‘gpagen’ function from the geomorph package. During the GPA process, the centroid size is calculated and separated as an independent factor to ensure it did not influence any subsequent analyses of shape variation.

S-Table 1. List of anatomical landmarks

| Number | Landmark names | Definition |
|---|---|---|
| 1 | n | nasion |
| 2 | g | glabella |
| 3 | sg | supraglabella |
| 4 | b | bregma |
| 5,6 | fmo | frontomalare orbitale |
| 7,8 | fmt | frontomalare temporale |
| 9 | rhi | rhinion |
| 10 | ns | nasospinale |
| 11,12 | sphn | sphenion |
| 13 | l | lambda |
| 14,15 | ast | asterion |
| 16,17 | ms | mastoidale |
| 18 | ba | basion |
| 19 | o | opisthion |
| 20 | i | inion |
| 21 | alv | alveolon |
| 22 | ids | infradentale superius/alveolare |
| 23,24 | po | porion |
| 25 | inl-m | midpoint of inferior nuchal line |
| 26,27 | d | dacryon |
| 28,29 | mo-h | highest point of margo orbitalis |
| 30,31 | or | the lowest point on the orbital margin |
| 32,33 | al | alare |
| 34,35 | sca-s | starting point of the supraciliary arch |
| 36,37 | sca-h | highest point of the supraciliary arch |
| 38,39 | sca-e | ending point of the supraciliary arch |
| 40,41 | cop | condyles occipitalis posterior |
| 42,43 | snl | ending points of superior nuchal line |
| 44,45 | or-li-tp | The turning point of lateral inferior orbital margin |
| 46,47 | zm | the most inferior point on the zygomaticomaxillary suture. |

| | | |
|---|---|---|
| 48,49 | iof | infra-orbital foramen |
| 50,51 | ft | The point of the superior temporal lobe located directly above the zygomatic process of the forehead, which looks most forward and inward. |
| 52,53 | zm-s | the most superior point of zygomaticomaxillary suture |
| 54,55 | pm-p | parietomastoid point: the most anterior point of parietomastoid suture |
| 56,57 | s-t-h | the highest point of the squamous part of the temporal bone |
| 58,59 | st | the point where the coronal suture crosses the temporal line |
| 60,61 | ms-l | the most lateral point of the mastoid process |
| 62,63 | mf | mandibular fossa |
| 64 | on | intersection of a line drawn horizontally across the frontal bone from the measuring points of the smallest forehead width and the median sagittal plane |
| 65,66 | max-tp-l | the turning point of the lateral margin of the maxilla |

S-Table 2. List of curves

| Number | Curve names | Numbers of semilandmarks |
|---|---|---|
| 1,2 | orbital margin | 25 each |
| 3 | along the midsagittal plane of the cranium (from n to o) | 52 |
| 4,5 | superciliary arch | 16 each |
| 6 | foramen magnum | 25 |
| 7 | piriform aperture | 25 |
| 8 | maxillary midline | 7 |
| 9 | median nuchal line | 19 |

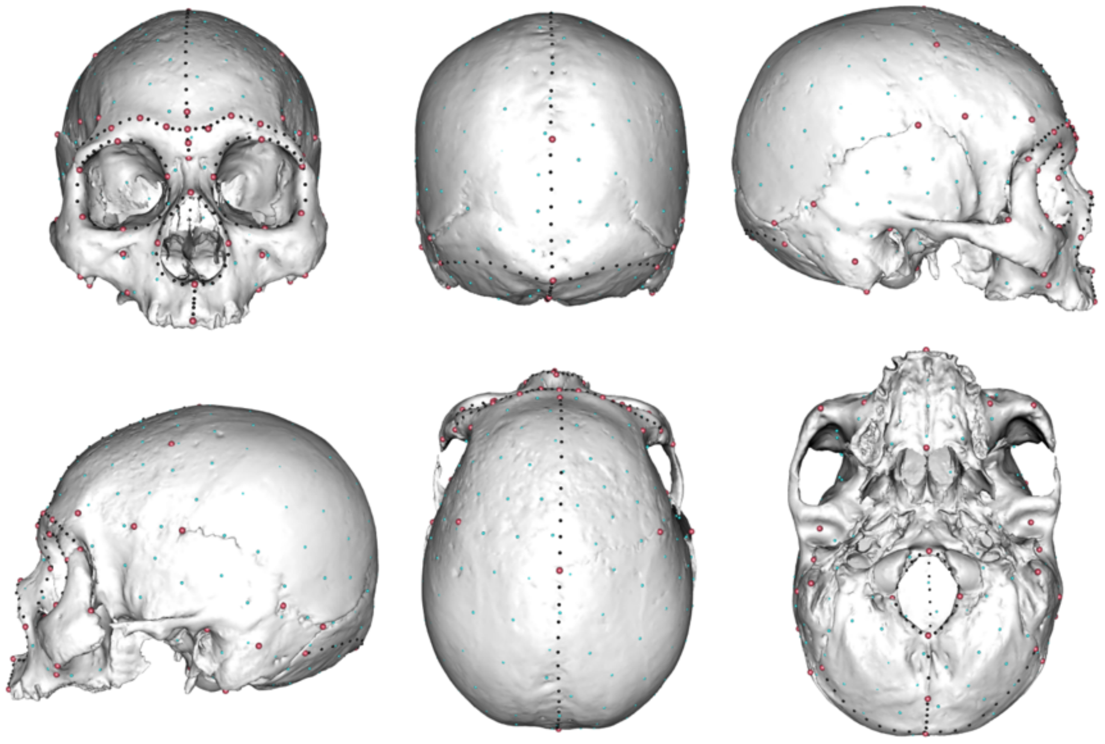

S-Figure 12. landmarks on the presented specimen (modern *H. sapiens* specimen curated in IVPP). Left: anterior view; middle: lateral view (left); right: inferior view. Red dots: fixed landmarks; black dots: curves; blue dots: patch landmarks.

After the GPA superimposition, Procrustes-registered shape coordinates of all specimens except the seven Yunxian2 versions were used to perform a Principal Component Analysis (PCA). PCA helps filter noise by reducing data dimensionality, focusing on the few components that explain the most variance. The seven versions of Yunxian 2 reconstructions were then projected into the PCA space, to reduce the influence of the reconstructed Yunxian 2 samples on the overall shape space. PCA was performed and visualized using the 'geomorph' and 'ggplot2' R packages (*44, 45*).

To further explore the morphological relationships between different *Homo* groups, we performed a between-group principal component analysis (bgPCA) on the GPA superimposed coordinates for all specimens, which accounted for 89.72% of the total variance. In the main text, Figure 3 provides a clearer distinction of the relative positions of different groups. The seven Yunxian 2 versions were projected into the morphospace constructed using human specimens, and the results indicate that their morphology occupies a unique position, not belonging to any known *Homo* species, but being closest to the *Homo longi*, *Homo neanderthalensis*, or *Homo erectus* groups. YX2 scores similarly to the *Homo erectus* group on bgPC1 axis and differs from the *Homo longi* group, indicating that it shares similarities with *H. erectus*, such as a low and flat cranial vault and a rounded piriform aperture, while lacking a prominent maxillary protrusion. On bgPC2 axis, however, Yunxian 2 is closer to the *H. longi* group and distinct from *H. erectus*, suggesting that it also shares characteristics with the *H. longi* group, such as a robust brow ridge, a lowered temporal line orientation, and a broad facial morphology. To better visualize the morphological differences represented by the principal components of our bgPCA analysis, we generated the morphologies of extreme values along the first two principal component axes. The anterior and lateral views of the crania shown on the bottom and left of Figure 3 in the main text represent the shape extremes of bgPC1 and bgPC2. These morphologies were created based on the mean shape (specimen

YNO227) of all specimens analyzed and were produced using the `shape.predictor()` function from the geomorph package and the `tps3d()` function from the Morpho package in R. Different from the Figure 3 in the main text, all the 7 reconstruction version of the Yunxian 2 were projected onto the bgPCA splace (S-Figure 13).

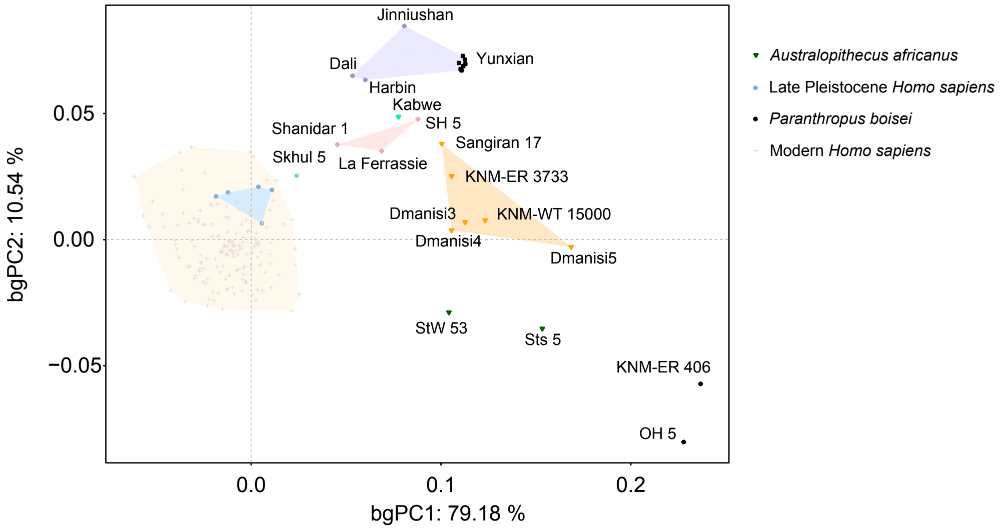


S-Figure 13. Between-group principal component analysis (bgPCA) of the Procrustes superimposed landmarks for all specimens used in this study. The first two bgPCs are shown, with the seven Yunxian 2 reconstruction versions projected onto the morphospace. Variation of the reconstruction of Yunxian falls in a small range when compared with other *Homo* fossils and extant human skulls.

We also performed a PCA analysis on the Procrustes coordinates of the seven reconstruction versions of Yunxian alone (S-Figure 14). On the PC1 axis, YX-1 shows a significant difference from the other versions, distinguished by its relative upward shift of the right zygomatic bone, which also affects the position of its right orbital, and the more prominent area on the occipital bone. On the PC2 axis, the main differences are related to the relative position of both zygomatic bones, the alignment of the parietotemporal region, and the more pronounced left supraorbital ridge, with highest scores from YX-2. The YX-4 version has the highest PC3 score, indicating that YX-4 has a slight contraction in the parietal regions on both sides compared to the other versions. For better visualization, we used the ‘meshDist’ function from the Morpho package to illustrate the shape differences between the maximum and minimum shape coordinates (representing the extreme variations in shape) on the first three principal components (see S-Figure 14). The final version is the averaged preferred version.

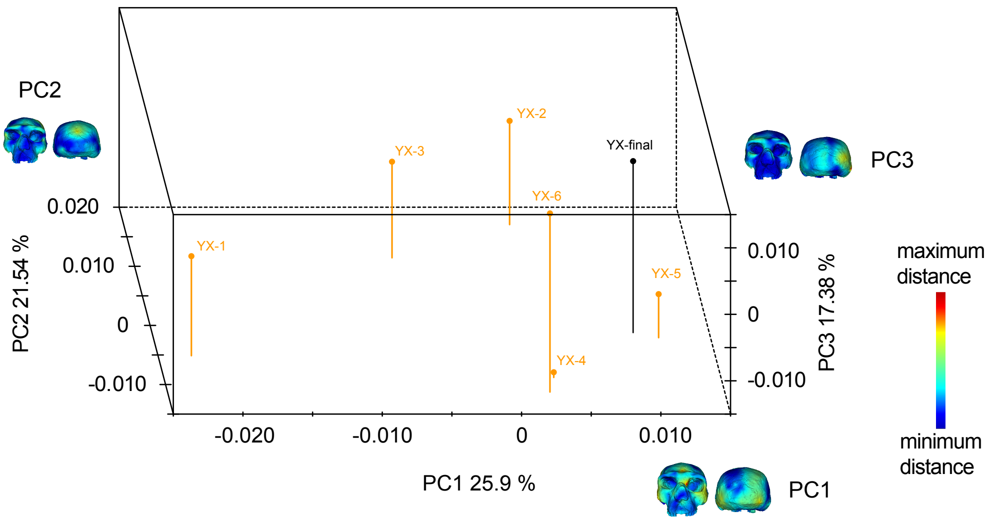


S-Figure 14. PCA on landmark configurations of the seven versions of Yunxian 2 reconstructions. Colored meshes illustrate the shape variations corresponding to the extreme values of each principal component. The Black dot indicates the preferred final version.

We also compared each of the six preliminary versions to the final version used for further analyses, standardized their difference intervals, and visualized these differences using 'meshDist' (S-Figure 15). The primary morphological differences between the six reconstructed versions of Yunxian 2 and the final reconstruction of Yunxian 2 are observed in the position of the left zygomatic bone, the alignment of the sphenoid region, and the right part of the occipital bone. Meanwhile, the reconstructed morphology around the vertex and maxillary region remains relatively stable across all versions.

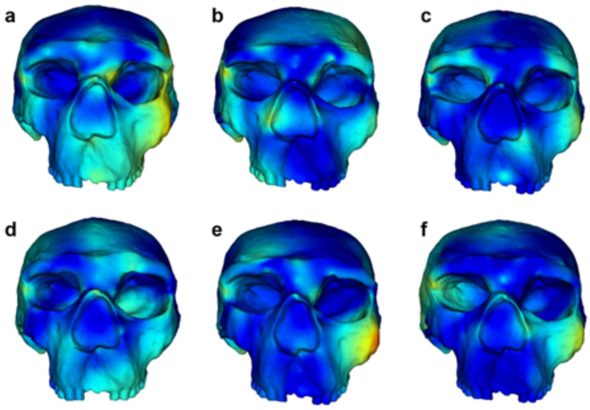


S-Figure 15. Local morphological differences between the first six versions and the final version of the reconstruction of Yunxian 2, implemented by the function 'meshDist' of R. Red areas indicate high variations and blue areas indicate low variations. a to f stands for the six versions of the reconstruction. In anterior view.

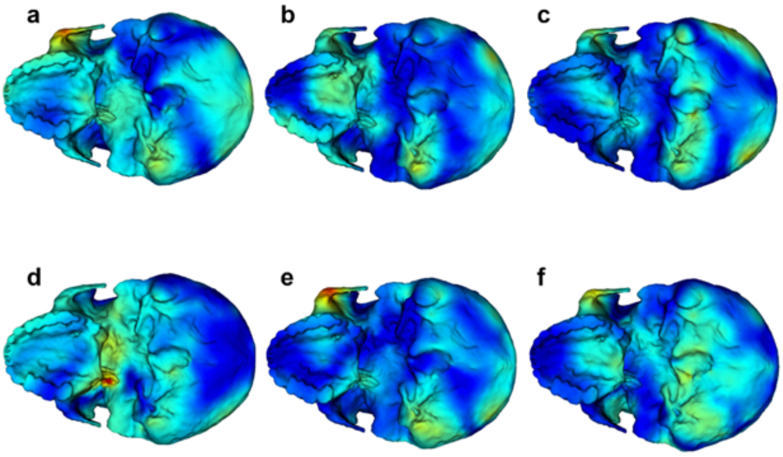


S-Figure 16. Local morphological differences between the first six versions and the final version of the reconstruction of Yunxian 2, implemented by the function 'meshDist' of R. Red areas indicate high variations and blue areas indicate low variations. a to f stands for the six versions of the reconstruction. In ventral view.

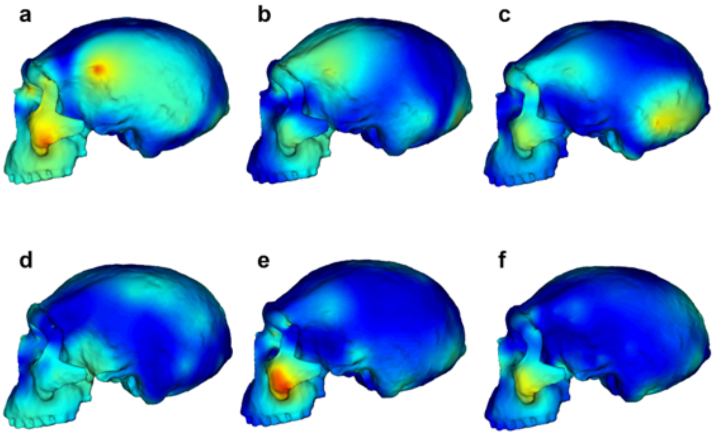


S-Figure 17. Local morphological differences between the first six versions and the final version of the reconstruction of Yunxian 2, implemented by the function 'meshDist' of R. Red areas indicate high variations and blue areas indicate low variations. a to f stands for the six versions of the reconstruction. In left lateral view.

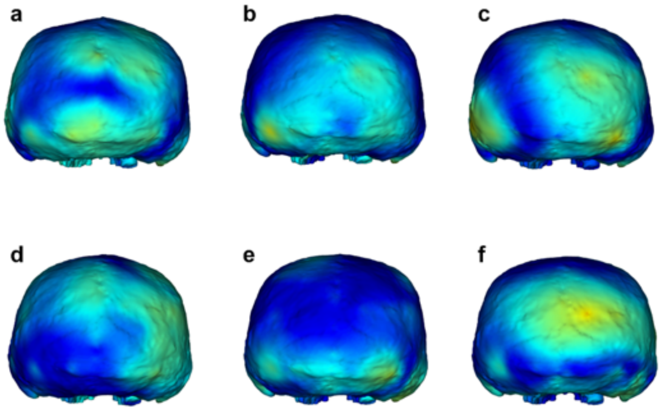


S-Figure 18. Local morphological differences between the first six versions and the final version of the reconstruction of Yunxian 2, implemented by the function 'meshDist' of R. Red areas indicate high variations and blue areas indicate low variations. a to f stands for the six versions of the reconstruction. In posterior view.

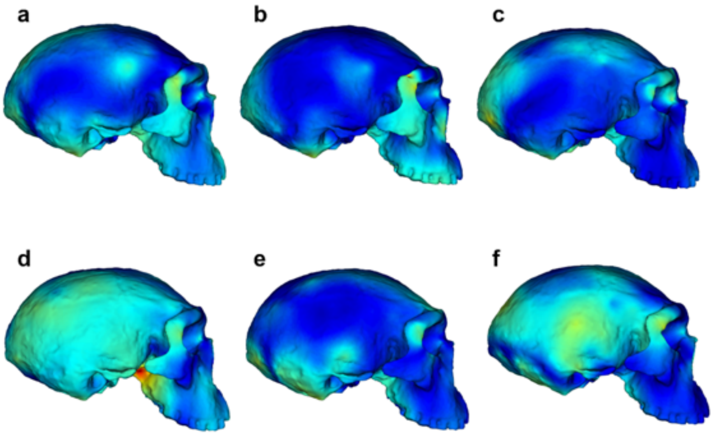

S-Figure 19. Local morphological differences between the first six versions and the final version of the reconstruction of Yunxian 2, implemented by the function 'meshDist' of R. Red areas indicate high variations and blue areas indicate low variations. a to f stands for the six versions of the reconstruction. In right lateral view.

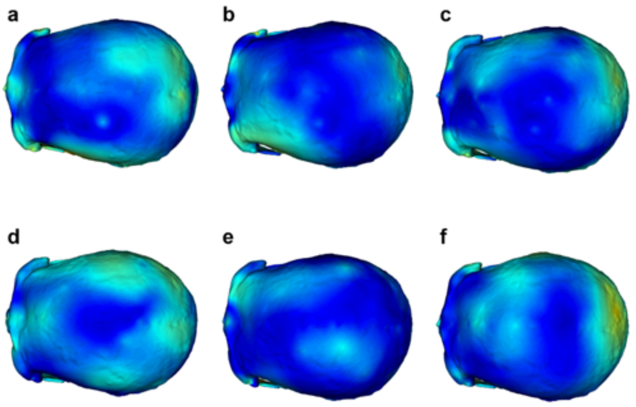


S-Figure 20. Local morphological differences between the first six versions and the final version of the reconstruction of Yunxian 2, implemented by the function 'meshDist' of R. Red areas indicate high variations and blue areas indicate low variations. a to f stands for the six versions of the reconstruction. In superior view.

Our geometric morphometric analysis reveals that the seven reconstruction versions of Yunxian 2 exhibit high morphological consistency compared to other human specimens, demonstrating the reliability of our reconstruction method. Additionally, the morphology of the final preferred version does not show any extreme variations in any of the analyses.

To evaluate the reliability of the reinstatement and reconstruction, we considered the reconstruction result in the context of determining the phylogenetic position of Yunxian 2. We used a Bootstrap resampling method to examine the stability of the phylogeny of *Homo* in relation to the hypothesized error of the reconstruction of Yunxian 2 (see section "Bootstrap analysis of the phylogenetic position of Yunxian 2" below).

**Reconstruction of the endocranial cast**

The matrix filled in the endocranial cavity is very dense. The high resolution CT scans do not result in good contrast between the fossilized bones and the matrix. Also, because of the presence of numerous cracks and high density minerals in the fossilized bones, it is impossible to segment the bones from the matrix. A regular procedure to generate the endocranial cast is therefore also impossible. To estimate the brain size and shape, we selected 46 landmarks (36 symmetrical and 10 in the midsagittal) in the cranial cavity of the CT scans of Yunxian 2 and measured bone thickness at these landmarks. With thickness of the bones, the landmarks were transformed into short "sticks" like those usually used in sculpture. The "sticks" were then transported to the reconstructed model of Yunxian 2. We generated a digital "balloon" within the shell of the reconstructed model of Yunxian 2. By digitally compressing, stretching, and sculpting, the "balloon" was fitted to all the lengths of the landmark "sticks".

**Morphology of the reconstructed Yunxian 2**

The occipital profile is moderately angled in Yunxian 2, less so than in *H. erectus* and *H. heidelbergensis* s.l, but more so than in *H. neanderthalensis* and *H. sapiens*, and without the protruding chignon found in many *H. neanderthalensis*. The upper scale of the occipital bone is about the same length in midline as the lower, and they meet at a minimally developed occipital torus.

The mastoid processes are low but massive, as in *H. erectus*, *H. heidelbergensis* s.l, *H. longi*, Dali, Jinniushan and Narmada. They point downwards and inwards and are much more massive than in Steinheim and most *H. neanderthalensis*. The Amud 1 *H. neanderthalensis* and *H. sapiens* have long, large, conical and downwards pointing mastoid processes. The external auditory meatus is high and anteroposteriorly relatively narrow. The anteroinferior tympanic plate is anteriorly flat and relatively thick, different from *H. erectus*, but similar to *H. heidelbergensis* s.l, *H. neanderthalensis* and *H. sapiens*.

The face is massive. In lateral view the midface is relatively high and projects anteriorly as in *H. erectus* and *H. heidelbergensis* s.l, but less than the degree seen in *H. neanderthalensis* and Bodo. The nasal root beneath the supraorbital torus is relatively flat, rather than deep. The zygomaxillary region is flat and faces anteriorly, but there is some alveolar prognathism below. In facial view (anterior view), the frontal bone is low, with a slight keel approaching bregma, and parasagittal flattening. The supraorbital torus is wide but not massively developed, strongest centrally, with a slight lateral reduction in thickness. The browridge is largely horizontal, with a flat shelf behind it, extending to the temporal lines. The inferior borders of the supraorbital torus are near horizontal above the orbits, which are widest superiorly and highest near the intersection with the zygomaxillary suture.

The upper face is wide, but the orbits are separated by only a moderate interorbital breadth, with a nasal saddle that projects in the midline. The lateral portions of the cheekbones are flat and high. There is a shallow depression corresponding to a canine fossa. There is a slight inferior prominence at the zygomaxillary suture on the left side. The infraorbital foramina are placed quite high and mesial with a small associated furrow extending below. The lower nasal region and maxillae lie well forward of the cheekbones, but without the maxillary inflation found in *H. neanderthalensis* and some *H. heidelbergensis* s.l crania.

The nasal aperture is high and very wide inferiorly, with raised margins along the lateral borders. There is an anterior nasal spine with a gutter sloping outwards and downwards on either side. The nasal volume must have been very large.

In superior view, the cranium is widest in the supramastoid area, and the frontal bone narrows behind the wide supraorbital torus, but post-orbital constriction is moderate, not as marked as in *H. erectus* and *H. heidelbergensis* s.l. fossils. The supraorbital torus only recedes slightly in superior view compared with *H. erectus* and *H. heidelbergensis* s.l. crania.

In posterior view the cranium is also widest in the supramastoid area, below which the massive mastoid processes slope inwards. The temporals and parietals do not converge as strongly as in *H. erectus* fossils, but there is no upper parietal lateral expansion, as found in recent *H. sapiens.* There is slight keeling near bregma, with parasagittal flattening either side. The occipital bone has no central protuberance or suprainiac depression, and the occipital torus is a weak and low ridge of bone that fades laterally. There is no 'en bombe' shape typical of *H. neanderthalensis*.

The inferior view illustrates the large breadth of the cranial base and palate, which is U-shaped and moderately deep, with thick alveolar bone. The central incisor sockets are slightly angled, suggesting some alveolar prognathism. A peg-shaped M3 is present on the left side, but none on the right. The zygomaticoalveolar root emerges from the region of the M1, and recedes only slightly as it progresses laterally.

The supramastoid crest is mound-like and stronger on the right side, and extends forward to overhang the auditory opening. Here it contributes to form a shelf above the recessed auditory meatus, from which the root of the zygomatic process emerges, distal to the mandibular fossa. The mandibular fossae are wide and quite deep, with the left side slightly deeper. The tympanic plates and petrous bones are not robustly built, and the long petrous axis is more coronally oriented than in *H. erectus* fossils. There is a well-marked digastric fossa mesial to the mastoid processes, and an occipitomastoid crest internal to that. The foramen magnum is long and oval and the occipital condyles quite small, while the nuchal plane is extensive, slightly recessed towards the foramen magnum, without a strong external occipital crest, and bordered by a modest occipital torus.

**Comparative morphology of the Yunxian 2 cranium**

As mentioned above, the Yunxian 2 cranium is large in size (S-Table 3), especially compared with most *H. erectus* crania, and it is therefore instructive to describe it against one of the few similarly sized *erectus* fossils - Sangiran 17. It is also about the same size as the Petralona cranium (considered as an example of *H. heidelbergensis* s.l by some researchers), so this also makes for a useful comparison, as Yunxian 2 has, at times, been allocated to both these taxa. Compared with a reconstruction of Sangiran 17 with the face repositioned, in lateral view both crania have a similar low vault height and great length, although the Yunxian 2 midline cranial profile is smoother compared with the more angular contours of Sangiran 17. Yunxian 2 has less total prognathism but a higher and more projecting nasal region, and a smaller, less projecting and thick supraorbital torus. Facially, Sangiran 17 has a deeper nasal root but a smaller nose in all dimensions. It has deeper cheekbones, especially laterally, with a wider infraorbital area and flared zygoma.

In rear view the crania are similar in height and breadth, but Yunxian 2 again has smoother contours, less convergent parietals, less midsagittal keeling and parasagittal flattening, and much less development of an occipital torus. The mastoid processes are larger in Yunxian 2. In basal view both crania have high biauricular and palatal breadths, but Yunxian has larger tooth crowns. The palates are both quite deep. At the rear, Sangiran 17 has a wider foramen magnum, more angled petrous bones, a central occipital crest and a flatter nuchal area, which is recessed as it approaches the much more strongly developed occipital torus. In superior view, Sangiran 17 has a larger supraorbital torus, especially laterally, and greater postorbital constriction. Cranial breadth is greatest above the mastoid area in both crania.

In relation to Petralona, both crania have relatively low and long vaults in lateral view, but Petralona has a higher braincase, with less flattened frontal and more curved parietals in midline. Both crania are prognathic with high and prominent nasal regions, and this area is even more projecting in Yunxian 2. Yunxian 2 has a smaller, less projecting and thick supraorbital torus, while that of Petralona is more like a strongly expressed version of a Neanderthal double-arched shape.

Petralona has slightly deeper cheekbones with a more inflated Neanderthal-like conformation centrally. The nasal bones are more projecting in Yunxian 2, but nasal size and inferred nasal volume appear similar. In rear view the crania are similar in profile, but Petralona is higher, with less keeling and parasagittal flattening than in Yunxian. Petralona has a wider and more strongly marked occipital torus. In basal view both crania have high biauricular and palatal breadths and similar petrous angulations. Petralona has a more squared-off palate anteriorly, while Yunxian has larger premolar and molar crowns, apart from the M3s. At the rear, Petralona has a wider foramen magnum, a central occipital crest and a flatter nuchal area, which is recessed as it approaches the more strongly developed occipital torus. In superior view, Petralona has a stronger supraorbital torus, especially laterally, and greater postorbital constriction. Cranial breadth is greatest above the mastoid area in both crania.

Unfortunately, comparisons with *H. antecessor* are limited by the incompleteness (and in some cases immaturity) of the Spanish fossils, although there are some similarities with Yunxian in zygomaxillary shape.

Overall, the Yunxian 2 cranium shows a distinctive combination of traits, and probably represents a distinct species of *Homo* from other designated Pleistocene human taxa such as *H. erectus*, *H. antecessor*, *H. heidelbergensis s.l*, *H. naledi*, *H. sapiens* and *H. neanderthalensis*.

S-Table 3. Measurements of the Yunxian 2 cranium and comparisons with other Middle-Late Pleistocene *Homo* cranial fossils. Linear measurements in millimeters, angles in degrees, ratios ranged from 0 to 100.

| | Yunxian 2 | Sangiran 17 | Dali | Hualongdong | Harbin | Jinniushan | Maba | Xuchang | Xujiayao | Kabwe | Petralona | Irhoud 1 |
|---|---|---|---|---|---|---|---|---|---|---|---|---|
| Cranial capacity | 1143 | 1020 | 1120 | 1150 | 1400 | 1390 | 1300 | 1800 | 1668 | 1249 | 1162 | 1375 |
| M1. GOL. g-op. Maximum cranial length | 203.67 | 203.6 | 212.2 | ? | 221.3 | 203.6 | 196.5 | 217 | ? | 206.3 | 208.9 | 198.9 |
| M1d. NOL. n-op. Nasio-occipital length. | 200.74 | 197.6 | 202.2 | ? | 212.9 | 195.4 | 188.5 | 212 | ? | 197.6 | 197 | 192.6 |
| M2. Glabella-inion length. | 202.12 | 202.8 | 203.5 | ? | 218.3 | 191.6 | ? | 217 | ? | 208 | 207.7 | 198.9 |
| Glabella-sphenobasion length | 97.27 | 90.2 | 96 | ? | 107.6 | 87.9 | ? | ? | ? | 97.9 | 100.2 | ? |
| Glabella-bregma chord. g-b chord. | 101.35 | 108.3 | 114.4 | 95.5 | 122.2 | 112.4 | 104.1 | 103 | ? | 118.2 | 106.9 | 104.4 |
| g-b/g-op. Glabella-bregma chord index | 49.8 | 53.2 | 53.9 | ? | 55.2 | 55.2 | 53 | 47.45 | ? | 57.3 | 51.2 | 52.5 |
| Length of basal temporal | 94.98 | 106 | 97 | ? | 116.1 | 104 | ? | 112 | 95.5 | 101.8 | 113.6 | ? |
| BTL/g-op. Basal temporal length index. | 46.6 | 52.1 | 45.7 | ? | 52.5 | 51.1 | ? | 51.65 | ? | 49.3 | 54.4 | ? |
| Entire temporal bone length | 104.29 | 100.3 | 97.5 | ? | 104.8 | 97.8 | ? | 103.8 | 91.76 | 97.6 | 104.9 | 102 |
| ETL/g-op. Entire temporal bone length index | 51.2 | 49.3 | 45.9 | ? | 47.4 | 48 | ? | 47.85 | ? | 47.3 | 50.6 | 51.3 |
| M5. BNL. Basion-nasion length. | 111.53 | 108.1 | 108.2 | ? | 117.5 | 97.9 | ? | ? | ? | 109.8 | 105.2 | ? |
| M5(1). Nasion-opisthion length. | 150.83 | 150.3 | 147.4 | ? | 152.2 | 147.7 | ? | ? | ? | 148.5 | 148.1 | ? |
| M6. ba-sphba. Basilar length. | 20.41 | 26.3 | 25.7 | ? | 23.7 | 22.6 | ? | ? | ? | 24.6 | 20.2 | ? |
| M7. FOL. Foramen magnum length (ba-o). | 41.6 | 41.4 | 41.1 | ? | 38 | 52.6 | ? | ? | ? | 41.4 | 45 | ? |
| M8. XCB. Maximum cranial breadth. | 163.67 | 157.2 | 156.4 | 144 | 164.1 | 144.1 | ? | 177 | 158.27 | 146.9 | 163.5 | 151.5 |
| M8c. Squama suture breadth | 151.82 | 146.4 | 139 | ? | 154.6 | 140.9 | ? | 167 | 148.87 | 142.8 | 143.6 | 141.6 |
| Maximum cranial breadth at supramastoid crest. Cranial vault. | 163.67 | 157.2 | 156.7 | ? | 164.1 | 144.1 | ? | 177 | 158.27 | 146.9 | 163.5 | 148.9 |
| Maximum biparietal breadth (Rightmire et al., 2006). Cranial vault. BPB | 151.39 | 145.8 | 155.1 | 138 | 157.4 | 143.8 | 146 | 170.7 | 148.29 | 144.4 | 148.9 | 148.9 |
| Ratio. Maximum biparietal breadth / maximum bimastoid breadth | 92.5 | 92.7 | 99 | ? | 95.9 | 99.8 | ? | 96.4 | 93.7 | 98.3 | 91.1 | 100.4 |
| M9. ft-ft. Least frontal breadth. Frontal. | 97.23 | 93.4 | 105.7 | 104 | 116.1 | 109 | 98.1 | 122.8 | ? | 97.6 | 106.8 | 108.5 |
| Postorbital constriction index. M9/M44 | 79.5 | ? | 90.4 | ? | 88.7 | 88.8 | 91.3 | ? | ? | 78.1 | 84.5 | 94.9 |
| M10. XFB. Maximum frontal breadth. Frontal. | 137.43 | 117.9 | 122.7 | 115.7 | 128.1 | 127.1 | 116.6 | 138.8 | 113.57 | 118.3 | 114.6 | 135.1 |
| M10b. STB. Bistephanic breadth (st-st) | 124.9 | 85.3 | 110 | ? | 121.9 | 115.3 | 112.7 | 138.8 | 110 | 111.2 | 96 | 120.6 |

| | | | | | | | | | | | | |
|---|---|---|---|---|---|---|---|---|---|---|---|---|
| M11. AUB. Biauricular breadth. | 150.93 | 147 | 145.8 | ? | 159.1 | 139.2 | ? | 173.62 | 150.49 | 135.9 | 152.2 | 143.1 |
| M11b. Biradicular breadth | 150.3 | 144.8 | 144.3 | ? | 158.8 | 132.8 | ? | 165.1 | 150.15 | 133.7 | 150.6 | 143 |
| M12. ASB. Biasterion breadth (ast-ast). Temporal. | 137.19 | 129.5 | 121.4 | ? | 134.4 | 126.4 | ? | 136.7 | 133.35 | 120.4 | 114.6 | 123.2 |
| M14. WCB. Minimum cranial breadth. | 65.71 | 75.9 | 80.6 | ? | 76.7 | 80.9 | ? | ? | ? | 72 | 83 | 87.6 |
| M16. Foramen magnum breadth. | 29.74 | 32.1 | 31.5 | ? | 30.6 | 35.4 | ? | ? | ? | 34.1 | 36.4 | ? |
| M17. BBH. Basion-Bregma height. | 109.9 | 108 | 116.4 | ? | 132.6 | 110.7 | ? | ? | ? | 128.8 | 121.9 | ? |
| M13a. MDB. Mastoid width. Temporal. | 18.51 | 15.8 | 18.4 | ? | 21.3 | 17.65 | ? | 13.9 | 13.65 | 16.1 | 17.5 | 13.3 |
| Maximum mastoid width | 25.06 | 22.5 | 24.65 | ? | 25.4 | 20.9 | ? | 20.95 | 16.58 | 20.7 | 23.3 | 20.2 |
| M19a. MDH. Mastoid height. Temporal. | 31.53 | 24.45 | 26.55 | ? | 30.65 | 20.5 | ? | 19.15 | 29.91 | 30.9 | ? | 28.3 |
| Minimum distance between the mastoid and supramastoid crests | 12.04 | 12.1 | 15.2 | ? | 14.25 | 14.8 | ? | 14.1 | 10.33 | 8.5 | 13.4 | 11.8 |
| Maximum bimastoid breadth | 161.24 | 146.9 | 140.6 | ? | 147.5 | 128 | ? | 166.1 | 149.53 | 143.3 | 151.5 | 148.3 |
| M20. Porion-bregmatic projective height. | 101.88 | 102.2 | 102.9 | ? | 113.9 | 99.7 | ? | 114.6 | 104.29 | 107.1 | 107.7 | 112.4 |
| AVH. Auriculare-vertex projective height | 95.46 | 97 | 101.6 | 100 | 109.5 | 97.1 | ? | 103.9 | 102.28 | 103.3 | 107.1 | 110.6 |
| Biporion breadth. Cranial vault. | 136.16 | 140.2 | 135.3 | ? | 147.9 | 128.7 | ? | 158.5 | 132.87 | 125.5 | 141.1 | 133.5 |
| Porion-basion projective height | 9.36 | 8.4 | 13.6 | ? | 18.8 | 12.9 | ? | ? | ? | 22.1 | 14.7 | ? |
| M29. FRC. n-b. Frontal sagittal chord. | 104.67 | 112 | 114.9 | 98.2 | 125.1 | 113.5 | 107.9 | 104.9 | ? | 120 | 108.3 | 107.7 |
| FRF. Nasion-subtense fraction. Frontal. | 53.95 | 50.7 | 47.7 | ? | 58.6 | 57.3 | 53.6 | 51.8 | ? | 59.7 | 56.6 | 39.7 |
| FRF/M29. Nasion-subtense fraction relative to the frontal sagittal chord. | 51.5 | 45.3 | 41.5 | ? | 46.8 | 50.5 | 49.7 | 49.4 | ? | 49.8 | 52.3 | 36.9 |
| Metopion subtense. Frontal. | 13.43 | 16.8 | 25.4 | 19.5 | 25.3 | 21.4 | 22.6 | 20.8 | ? | 21.6 | 19.9 | 22.1 |
| Metopion subtense/M29. Metopion subtense relative to the nation-bregma chord. | 12.8 | 15 | 22.1 | 19.9 | 20.2 | 18.9 | 20.9 | 19.8 | ? | 18 | 18.4 | 20.5 |
| Lower frontal inclination angle (m-g-i) | 53.21 | 57.7 | 68.3 | ? | 64 | 59.8 | 69 | 64.5 | ? | 60.5 | 57.5 | 73.1 |
| M30. PAC. Parietal sagittal chord. | 106.98 | 87.8 | 109.3 | 95.5 | 106.1 | 104.7 | 112.1 | 109.2 | 103.59 | 110.8 | 104.2 | 122.7 |
| M30(2). Bregma-sphenion chord (b-sphn). | 100.3 | 92.9 | 91.4 | ? | 93.9 | 95.7 | 89 | 100.5 | ? | 89.5 | 90.8 | 96.2 |
| M30(3). Lambda-asterion chord (l-ast). | 133.9 | 96.2 | 91.8 | ? | 105.8 | 79.9 | ? | 99.3 | 143.6 | 90 | 91.4 | 91.6 |
| ASI. Asterion-inion chord | 86.49 | 79.1 | 76 | ? | 81.7 | 72.6 | ? | 80.4 | 76.36 | 75.2 | 67 | 76.8 |
| M30c. Bregma-asterion chord (b-ast). Parietal. | 134.1 | 133.3 | 131.9 | ? | 146 | 121.7 | ? | 146.4 | 143.89 | 139.3 | 141 | 144.3 |
| Parietal chord index. M30/M1 | 52.5 | 43.1 | 51.5 | ? | 47.9 | 51.4 | 57.05 | 50.35 | ? | 53.7 | 49.9 | 61.7 |
| M31. OCC. l-o. Occipital sagittal chord. | 84.77 | 88 | 90.3 | ? | 103.1 | 72.7 | ? | ? | 57.195 | 91.1 | 91.6 | ? |

| | | | | | | | | | | | | |
|---|---|---|---|---|---|---|---|---|---|---|---|---|
| M31(1). LIC. Lambda-inion chord (l-i) | 54.78 | 75.4 | 69.2 | ? | 79.8 | 53.2 | 52 | 84.2 | 96.92 | 55.7 | 64.7 | 47.3 |
| Lambda-opisthocranion chord l-op | 50.29 | 72 | 49.5 | ? | 56.7 | 39.6 | 52 | 82.3 | 79.93 | 43.4 | 61.9 | 47.3 |
| M31(2). Inion-opisthion chord (i-o) | 57.44 | 53.1 | 39 | ? | 62.2 | 39.2 | ? | ? | 52.43 | 63.6 | 59.6 | ? |
| Ratio. M31(1)/M31(2). Length of occipital (lambda-inion) plane compared to nuchal (inion- opisthion) plane. X100. | 95.4 | 142 | 177.4 | ? | 128.3 | 135.7 | ? | ? | 184.9 | 87.6 | 108.6 | ? |
| Opisthocranion-opisthion chord | 62.32 | 55.5 | 64.5 | ? | 77.9 | 58.4 | ? | ? | 70.52 | 67.7 | 62.6 | ? |
| Sphenobasion-opisthion length | 61.43 | 67.4 | 63.9 | ? | 58.2 | 70.6 | ? | ? | ? | 63.6 | 63.6 | ? |
| M32(1). Frontal inclination angle (b-n-i). Frontal. | 45.03 | 48.3 | 51.8 | ? | 50.9 | 48.6 | 51.5 | 51.4 | ? | 49.4 | 50.9 | 52.9 |
| M32(2). Bregma angle (b-g-i). Frontal. | 42.61 | 45 | 47.2 | ? | 46.9 | 45 | 48.5 | 47.2 | ? | 45.9 | 45 | 49.7 |
| M32(5). FRA. Frontal angle (b-m-n, degree) | 150.88 | 146.3 | 129.7 | 134.9 | 135.9 | 138.7 | 134.7 | 137.3 | ? | 141.1 | 139.7 | 132.7 |
| M33d. OCA. Occipital angle (degree). | 97.72 | 84.6 | 94.2 | ? | 93.4 | 95.6 | ? | ? | 49.19 | 102.2 | 95.7 | ? |
| M33e. PAA. Parietal angle (degree). | 150.53 | 162.4 | 145.8 | 141.3 | 146.5 | 142 | 139.2 | 152.7 | 145.14 | 145.8 | 141.4 | 150.3 |
| M33(4). Lambda-inion-opisthion angle (l-i-o) | 97.85 | 84.6 | 98.4 | ? | 93.3 | 102.8 | ? | ? | 95.26 | 100.5 | 95.8 | ? |
| Temporal squama length (Martínez and Arsuaga, 1997). | 70.2 | 83 | 70.9 | ? | 71.4 | 68.5 | ? | 70.8 | 64.65 | 69.3 | 62.7 | 79.3 |
| Temporal squama height (Martínez and Arsuaga, 1997). | 48.15 | 43.4 | 46.5 | 45 | 52.9 | 41.2 | ? | 39.8 | 47.52 | 51.7 | 47.2 | 38.3 |
| Temporal squama angle (Martínez and Arsuaga, 1997). | 64.87 | 36.5 | 46.8 | ? | 52.8 | 50.2 | ? | 38.7 | 29.515 | 52 | 36.2 | 32 |
| Temporal muscle attachment height | 64.48 | 89.3 | 89.9 | ? | 83.3 | 87.6 | ? | 76.1 | 72.33 | 91.4 | 88 | 75.8 |
| Temporal muscle attachment length | 131.4 | 151.7 | 131.4 | ? | 143.4 | 141.9 | 140 | 140 | ? | 143.7 | 122.3 | 135.2 |
| Temporal muscle attachment length index | 64.5 | 74.5 | 61.9 | ? | 64.8 | 69.7 | 71.25 | 64.55 | ? | 69.7 | 58.5 | 68 |
| Transverse tympanic width | 27.53 | 31.6 | 23.2 | ? | 22.5 | 20.7 | ? | ? | 22.14 | 26 | 25 | ? |
| Tympanic axis angle | 94.92 | 87.6 | 101.2 | ? | 94.8 | 100.4 | ? | ? | 90.44 | 105.8 | 92.1 | ? |
| Tympanic axis length | 27.77 | 25.8 | 25.6 | ? | 23.1 | 21.4 | ? | ? | 22.19 | 26.5 | 26.2 | ? |
| Petrous axis angle | 114.58 | 127.6 | 136.9 | ? | 128.4 | 131.7 | ? | ? | 56.4 | 122 | ? | ? |
| Petrous axis length | 15.26 | 12.8 | 12.9 | ? | 14.5 | 13 | ? | ? | 10.66 | 11.6 | ? | ? |
| Tympanic angle | 18.28 | 39.9 | 34.9 | ? | 38.6 | 30.4 | ? | ? | 22.82 | 16.8 | ? | ? |
| Postglenoid-ectoglenoid length | 8.255 | 20.8 | 18.3 | ? | 17.4 | 14.5 | ? | 19.4 | 15.76 | 18.1 | 12.7 | 24.2 |
| Postglenoid-entoglenoid length | 25.13 | 25.5 | 21.3 | ? | 24.8 | 15.4 | ? | ? | 23.62 | 23.8 | 23.1 | 17.8 |
| Ectoglenoid-entoglenoid length | 31.2 | 29.1 | 29.5 | ? | 29.9 | 23.4 | ? | ? | 27.97 | 30.2 | 27.8 | 30.2 |
| Chord length of the parietomastoid suture. Incisura parietalis - asterion. | 21.39 | 17.2 | 26.7 | ? | 33.4 | 29.3 | ? | 33 | 27.04 | 28.3 | 42.2 | 22.7 |

| | | | | | | | | | | | | |
|---|---|---|---|---|---|---|---|---|---|---|---|---|
| Occipital height (l-sphba) | 119.14 | 111.7 | 123.8 | ? | 130.7 | 117 | ? | ? | ? | 125.3 | 120.5 | ? |
| Occipital subtense | 36.6 | 46 | 41.7 | ? | 47.7 | 33.8 | ? | ? | 48.02 | 37.1 | 41.8 | ? |
| Occipital plane index. l-op/bi-ast | 36.7 | 55.6 | 40.8 | ? | 42.2 | 31.3 | ? | 60.2 | 59.9 | 36 | 54 | 38.4 |
| Inion-endinion | ? | ? | 26.4 | ? | 25.1 | ? | ? | ? | 30.27 | 35.3 | ? | 37.9 |
| M40. BPL. Basion-prosthion length. Cranial vault. | 126.05 | 130.6 | 107.8 | ? | 125.1 | 101.6 | ? | ? | ? | 117.1 | 117.1 | ? |
| M43. FMT. Bi-frontomalare temporale breadth (fmt-fmt). Frontal. | 132.44 | 117 | 124.4 | ? | 140.2 | 129 | 117.2 | 140 | ? | 134.4 | 133.2 | 127.2 |
| M43a. FMB. Bifrontal breadth. Frontal. | 121.4 | 110.4 | 116.9 | ? | 130.9 | 122.7 | 107.4 | 120 | ? | 124.9 | 126.4 | 113 |
| M43b. NAS. Nasio-frontal subtense. | 24.99 | 24.1 | 20.4 | ? | 27.1 | 21.4 | 18.8 | 13.1 | ? | 26.4 | 21.7 | 19 |
| M43b/M43a. Nasion-frontal subtense relative to the bifrontal breadth. | 20.6 | 21.8 | 17.5 | ? | 20.7 | 17.4 | 17.5 | 10.9 | ? | 21.1 | 17.2 | 16.8 |
| Frontal. Supraorbital torus breadth. XSOT | 132.59 | 121.7 | 129 | 122 | 145.7 | 135.5 | 119 | 140.7 | ? | 139.4 | 135.2 | 128.3 |
| M44. EKB. Biorbital breadth (ek-ek) | 122.31 | 110.4 | 116.9 | 109 | 130.9 | 122.7 | 107.4 | ? | ? | 124.9 | 126.4 | 114.3 |
| Supraorbital torus thickness central. Frontal. | 13 | 16.4 | 18 | 16.5 | 15 | 10.8 | 12.4 | 13.8 | ? | 20 | 16.5 | 13.1 |
| Supraorbital torus thickness lateral. Frontal. | 12 | 14.7 | 13 | 11.5 | 15.5 | 13.4 | 9.5 | 12.7 | ? | 17.2 | 11.5 | 17.2 |
| Supraorbital torus thickness medial. Frontal. | 19.19 | 18 | 19.2 | 15.25 | 21.4 | 15.8 | 16.8 | ? | ? | 20 | 20.1 | 18.8 |
| M45. ZYB. Bizygomatic breadth (zy-zy) | 146.08 | 154 | 143.8 | ? | 162.4 | 148.5 | ? | ? | 159.5 | 143.4 | 158.1 | ? |
| M45(1). JUB. Bijugal breadth. | 137.16 | 131.3 | 126.5 | ? | 138.6 | 130.9 | 114.2 | ? | ? | 134.1 | 138.2 | 127.9 |
| M46b. ZMB. Bimaxillary breadth. | 118.9 | 118.2 | 111.4 | 111.5 | 113.8 | 116.9 | ? | ? | ? | 110.6 | 123.9 | 108.1 |
| Facial proportion. M46b/M43 | 89.8 | 101 | 89.5 | ? | 81.2 | 90.6 | ? | ? | ? | 82.3 | 93 | 85 |
| Temporal gutter angle | 62.97 | 40.3 | 59.4 | ? | 54.8 | 58.5 | 48.2 | ? | ? | 51 | 23.5 | 40.4 |
| M48. NPH. Upper facial height. Nasion-prosthion height (n-pr) | 89.6 | 71 | 64.1 | 80.6 | 76.4 | 72.5 | ? | ? | ? | 90.9 | 90.9 | 83.5 |
| M48(1). Nasospinale-prosthion distance (ns-pr) | 31.38 | 14.8 | 16.8 | 16.8 | 16.2 | 19.7 | ? | ? | ? | 26.9 | 22.4 | 23 |
| M74. Upper facial angle. Nasion-prosthion relative to the FH | 70.71 | 66.4 | 82.1 | 91 | 78 | 82.9 | ? | ? | ? | 82.2 | 81.2 | 75.9 |
| M74(1). Clivus-alveolar plane angle | 62.34 | 66.3 | 55.1 | ? | 57.9 | 74.1 | ? | ? | ? | 65 | 67.3 | 64.8 |
| Nasiospinale-alveolare length | 33.5 | ? | ? | ? | 19.5 | 23 | ? | ? | 16.22 | 31.5 | 28.8 | ? |
| Nasiospinale-alveolare angle | 71.19 | ? | ? | 93 | 81.5 | 86.7 | ? | ? | ? | 84.9 | 92.5 | ? |
| M48d. WMH. Cheek height. | 31.92 | 36.4 | 22.6 | 25 | 28.3 | 27.6 | ? | ? | ? | 26.5 | 32.6 | 23.45 |
| M49a. DKB. Interorbital breadth (d-d) | 27.93 | 29.2 | 20.6 | ? | 24.7 | 26.2 | 19.9 | ? | ? | 25.3 | 31.7 | 27.3 |
| M50. IOW. Anterior interorbital breadth (mf-mf) | 23.36 | 23.7 | 23.5 | ? | 31.4 | 33.6 | 22.4 | ? | ? | 28.1 | 31 | 26.6 |

| | | | | | | | | | | | | |
|---|---|---|---|---|---|---|---|---|---|---|---|---|
| M51. Orbital breadth (mf-ek) | 49.88 | 47.6 | 50.5 | ? | 52.4 | 46.4 | 47 | ? | ? | 52.7 | 49 | 45.1 |
| M51a. OBB. Orbital breadth. | 46.18 | 45.4 | 51 | 43 | 57.3 | 50.7 | 45.5 | ? | ? | 49.5 | 48.5 | 44.9 |
| M52. OBH. Orbital height. | 33.91 | 36.4 | 34.2 | 40.6 | 41.6 | 36 | 38 | ? | ? | 39.2 | 33.9 | 33.3 |
| M54. NBL. Nasal breadth. | 38.88 | 33.2 | 34.9 | 30 | 36.2 | 31.8 | ? | ? | 27.46 | 31.5 | 38.5 | 36 |
| M57(2). Upper nasal breadth of nasal bones | 19.08 | 13.6 | 8.5 | ? | 17.9 | 13 | 13.4 | ? | ? | 17.7 | 15 | 15.9 |
| M55. NH. Nasal height. n-ns. | 58.96 | 55.9 | 48.4 | 63.5 | 61 | 52.1 | ? | ? | ? | 63.9 | 68 | 56.5 |
| M61. MAB. Maxilloalveolar breadth. | 81.34 | 78.1 | 73.9 | 74 | 79.2 | 65.5 | ? | ? | 69.47 | 84.8 | 86.1 | 73.5 |
| Maxilloalveolar length | 81.42 | 62.4 | 69.1 | ? | 74 | 65.1 | ? | ? | 66 | 67.2 | 68.2 | 63.1 |
| Maxillary palate length | 49.96 | 36.7 | 49.8 | ? | 45.8 | 39.5 | ? | ? | 39.81 | 42.4 | 48.3 | 45 |
| M63. Internal palatal breadth | 45.27 | 43 | 40.3 | 46 | 39 | 34.8 | ? | ? | 38 | 47.4 | 48.4 | 40.6 |
| External alveolar breadth at canine level | 54.87 | 51.5 | 51.3 | ? | 68.8 | 46.4 | ? | ? | 48.92 | 52.4 | 51.7 | 50.8 |
| External alveolar breadth at P3 level | 66.44 | 60.9 | 62 | ? | 73.1 | 54.7 | ? | ? | 58.67 | 62.1 | 65.7 | 56.6 |
| External alveolar breadth at P4 level | 72.84 | 66.1 | 66.2 | ? | 74.6 | 60.5 | ? | ? | 63.17 | 67.7 | 73.2 | 65.1 |
| External alveolar breadth at M1/M2 level | 80.76 | 72.3 | 67.2 | ? | 74.5 | 64.8 | ? | ? | 67.1 | 78.7 | 85.2 | 73.5 |
| M61 relative to the external palatal breadth at the canine level. | 148.2 | 151.7 | 144.1 | ? | 115.1 | 141.2 | ? | ? | 142 | 161.8 | 166.5 | 144.7 |
| M61 relative to the external palatal breadth at the P3 level. | 122.4 | 128.2 | 119.2 | ? | 108.3 | 119.7 | ? | ? | 118.4 | 129.1 | 131.1 | 129.9 |
| M76a. SSA. Zygomaxillary angle (degree). | 114.24 | 125.6 | 122.6 | ? | 112.3 | 119.5 | ? | ? | ? | 107.3 | 114.9 | 116.8 |
| Subspinale subtense | 35.73 | 30.3 | 30.5 | ? | 38.2 | 33.9 | ? | ? | ? | 40.8 | 40.2 | 33.5 |
| M77a. NFA. Nasio-frontal angle (fm:a-n-fm:a) | 133.93 | 132.8 | 141.8 | 154.2 | 135.2 | 142.7 | 143 | ? | ? | 136 | 142 | 145.3 |
| Width of nasal bridge. Rightmire 1998 | 25.78 | 28.7 | 24.7 | ? | 29.9 | 31 | 20.1 | ? | ? | 27.4 | 32.5 | 27.8 |
| Nasal bridge height. Rightmire 1998. | 12.06 | 20.5 | 12.1 | ? | 16.2 | 11.6 | 12.4 | ? | ? | 16.8 | 18.1 | 20.5 |
| Nasal bridge index. Rightmire 1998. | 46.8 | 71.4 | 49 | ? | 54.2 | 37.4 | 61.7 | ? | ? | 61.3 | 55.7 | 73.7 |
| Nasal bridge angle. Rightmire 1998. | 94.7 | 73.4 | 91.4 | ? | 85.8 | 103.4 | 75.7 | ? | ? | 77.3 | 83.6 | 67.4 |
| M41c. XML. Maximum malar length | 61.41 | 70 | 54 | ? | 52.7 | 52 | ? | ? | ? | 56.6 | 59.1 | ? |
| Maximum malar height (Rightmire et al., 2006) | 48.42 | 62.3 | 43.1 | ? | 52.1 | 41.6 | ? | ? | ? | 54.1 | 55 | 48.7 |
| Zygomaxillare anterior-zygoorbitale (zm:a-zo) | 40.82 | 47.9 | 32.9 | ? | 33.9 | 35.4 | ? | ? | ? | 33.4 | 44.9 | 32.9 |
| M41d. MLS. Malar subtense | 17.29 | 22.6 | 10.8 | ? | 11.5 | 10.4 | ? | ? | ? | 12.3 | 14.8 | ? |
| IZM height. Inferior zygomatic margin height. | 22.29 | 14.2 | 16.9 | ? | 18.9 | 21.2 | ? | ? | ? | 26.5 | 30.2 | 22.9 |

| | | | | | | | | | | | | |
|---|---|---|---|---|---|---|---|---|---|---|---|---|
| Malar angle | 42.88 | 120.3 | 50.3 | ? | 67.9 | 58.8 | ? | ? | ? | 73.8 | 67.8 | 66.7 |
| Infraorbital plate angle in parasagittal plane. | 81.58 | 101.25 | 68.48 | ? | 74.12 | 74.92 | ? | ? | ? | 78.94 | 85.58 | 74.2 |
| Orbital incline angle in parasagittal plane. | 87.23 | 93.66 | 105.78 | ? | 105.95 | 99.3 | 93 | ? | ? | 106.9 | 98.54 | 103.31 |
| bimandibular fossa breath | 102.84 | 112.1 | 104.7 | ? | 116.2 | 97.7 | ? | ? | 123.3 | 107.8 | 109.5 | 113.6 |
| Mandibular fossa depth | 13.17 | 10.5 | 6.5 | ? | 11.8 | 9.3 | ? | 8 | 12.13 | 10.1 | 13.8 | ? |
| Midfacial prognathism | 30.97 | 26.3 | 5.6 | ? | 15.8 | 9.1 | ? | ? | ? | 12.3 | 13.4 | 16.7 |
| Dental. Upper I1. Mesiodistal length. | ? | ? | ? | ? | ? | 10.1 | ? | ? | 9.77 | 8.2 | ? | ? |
| Dental. Upper I1. Buccolingual width. | ? | ? | ? | ? | ? | 8.5 | ? | ? | 8.19 | 9.7 | ? | ? |
| Dental. Upper I1. Width/length. | ? | ? | ? | ? | ? | 84.2 | ? | ? | 83.8 | 118.3 | ? | ? |
| Dental. Upper I1. Area relative to M1. LxW of I1 / LxW of M1 | ? | ? | ? | ? | ? | 59.7 | ? | ? | 41.9 | 46 | ? | ? |
| Dental. Upper I1. Labial convexity. ASUDAS grades. UI1LC | ? | ? | ? | ? | ? | 2 | ? | ? | 4 | ? | ? | ? |
| Dental. Upper I1. Lingual shoveling. ASUDAS grades. UI1SS | ? | ? | ? | ? | ? | 2 | ? | ? | 2 | ? | ? | ? |
| Dental. Upper I1. Labial shoveling. ASUDAS grades. UI1DS | ? | ? | ? | ? | ? | 0 | ? | ? | 0 | ? | ? | ? |
| Dental. Upper I1. Tuberculum Dentale. ASUDAS grades. UI1TD | ? | ? | ? | ? | ? | 0 | ? | ? | 2 | ? | ? | ? |
| Dental. Upper I2. Mesiodistal length. | 8.01 | ? | ? | 5.8 | ? | 7.9 | ? | ? | ? | 6.3 | ? | ? |
| Dental. Upper I2. Buccolingual width. | 9.01 | ? | ? | 6.9 | ? | 7.8 | ? | ? | ? | 8.2 | ? | ? |
| Dental. Upper I2. Width/length. | 112.5 | ? | ? | 119 | ? | 98.7 | ? | ? | ? | 130.2 | ? | ? |
| Dental. Upper I2. Area relative to I1. LxW of I2 / LxW of I1 | ? | ? | ? | ? | ? | 42.9 | ? | ? | ? | 64.9 | ? | ? |
| Dental. Upper I2. Labial convexity. ASUDAS grades as UI1LC. | 5.5 | ? | ? | 4 | ? | 3 | ? | ? | ? | 4 | ? | ? |
| Dental. Upper I2. Lingual shoveling. ASUDAS grades. UI2SS | 0.5 | ? | ? | 3 | ? | 2.5 | ? | ? | ? | ? | ? | ? |
| Dental. Upper I2. Labial shoveling. ASUDAS grades. UI2DS | 0 | ? | ? | 0 | ? | 0 | ? | ? | ? | ? | ? | ? |
| Dental. Upper I2. Tuberculum Dentale. ASUDAS grades. UI2TD | 0 | ? | ? | 0 | ? | 0 | ? | ? | ? | ? | ? | ? |
| Dental. Upper canine. Mesiodistal length. | 9.02 | 9.3 | ? | ? | ? | 8.8 | ? | ? | 10.89 | 10.3 | 10 | ? |
| Dental. Upper canine. Buccolingual width. | 11.29 | 9 | ? | ? | ? | 9.5 | ? | ? | 10.27 | 10.9 | 9.7 | ? |
| Dental. Upper canine. Width/length. | 125.2 | 96.8 | ? | ? | ? | 108 | ? | ? | 94.3 | 105.8 | 97 | ? |
| Dental. Upper canine. Area relative to M1. LxW of C1 / LxW of M1 | 47.65 | 62.4 | ? | ? | ? | 58.2 | ? | ? | 58.5 | 64.9 | 55 | ? |
| Dental. Upper canine. Tuberculum Dentale. ASUDAS grades. UCTD | 0 | 0 | ? | ? | ? | 1 | ? | ? | 0.5 | ? | 0 | ? |
| Dental. Upper canine. Mesial marginal ridge. ASUDAS grades. UCMR | 0 | 1 | ? | ? | ? | 3 | ? | ? | 2.5 | ? | 1 | ? |

| | | | | | | | | | | | | |
|---|---|---|---|---|---|---|---|---|---|---|---|---|
| Dental. Upper canine. Distal marginal ridge. ASUDAS grades. As UCMR | 1.5 | 1 | ? | ? | ? | 2 | ? | ? | 2.5 | ? | 1 | ? |
| Dental. Upper canine. Distal accessory ridges. ASUDAS grades. UCDR | ? | 0 | ? | ? | ? | 0 | ? | ? | 1 | ? | 0 | ? |
| Dental. Upper P3. Mesiodistal length. | 9.215 | 6.9 | ? | 7.7 | ? | 8.2 | ? | ? | 9.35 | 7.3 | 8.6 | ? |
| Dental. Upper P3. Buccolingual width. | 15.125 | 9.7 | ? | 9.9 | ? | 10.6 | ? | ? | 11.14 | 10.8 | 12.7 | ? |
| Dental. Upper P3. Width/length. | 163.6 | 140.6 | ? | 129.9 | ? | 129.3 | ? | ? | 119.1 | 147.9 | 147.7 | ? |
| Dental. Upper P3. Area relative to M1. LxW of P3 / LxW of M1 | 66.3 | 49.9 | ? | 53.1 | ? | 60.5 | ? | ? | 54.5 | 45.6 | 61.9 | ? |
| Dental. Upper P3. Mesial accessory ridges. ASUDAS grades. UP3M MXPAR | 0 | 0 | ? | 2.5 | ? | 0 | ? | ? | 4 | ? | 0 | ? |
| Dental. Upper P3. Distal accessory ridges. ASUDAS grades. UP3D MXPAR | 0 | 0 | ? | 2.5 | ? | 1.5 | ? | ? | 2.5 | ? | 0 | ? |
| Dental. Upper P4. Mesiodistal length. | 8.495 | ? | ? | ? | ? | 7.9 | ? | ? | 8.93 | ? | 7.6 | ? |
| Dental. Upper P4. Buccolingual width. | 14.68 | ? | ? | ? | ? | 10.6 | ? | ? | 11.01 | ? | 12.7 | ? |
| Dental. Upper P4. Width/length. | 172.25 | ? | ? | ? | ? | 134.2 | ? | ? | 123.3 | ? | 167.1 | ? |
| Dental. Upper P4. Area relative to M1. LxW of P4 / LxW of M1 | 59.35 | ? | ? | ? | ? | 58.3 | ? | ? | 51.4 | ? | 54.7 | ? |
| Dental. Upper P4. Mesial accessory ridges. ASUDAS grades. UP4M MXPAR | 0 | ? | ? | ? | ? | 0 | ? | ? | 1 | ? | 0 | ? |
| Dental. Upper P4. Distal accessory ridges. ASUDAS grades. UP4D MXPAR | 0 | ? | ? | ? | ? | 1.5 | ? | ? | 1 | ? | 2 | ? |
| Dental. Upper M1. Mesiodistal length. | 13.37 | 10.9 | ? | 11.65 | ? | 11.5 | ? | ? | 13.63 | 12.9 | 12.6 | ? |
| Dental. Upper M1. Buccolingual width. | 16.085 | 12.3 | ? | 12.45 | ? | 12.5 | ? | ? | 14.02 | 13.4 | 14 | ? |
| Dental. Upper M1. Width/length. | 120.7 | 112.8 | ? | 106.9 | ? | 108.7 | ? | ? | 102.9 | 103.9 | 111.1 | ? |
| Dental. Upper M1. Metacone size. ASUDAS grades. UM1MC | 4.5 | 3 | ? | 3 | ? | 1.5 | ? | ? | 5 | 3 | 3 | ? |
| Dental. Upper M1. Hypocone size. ASUDAS grades. UM1HC | 5.5 | 4 | ? | 5 | ? | 4.5 | ? | ? | 4 | 3.5 | 4 | ? |
| Dental. Upper M1. Cusp 5 size. ASUDAS grades. UM1C5 | 1.5 | 0.5 | ? | 3.5 | ? | 1.5 | ? | ? | 1 | 1.5 | 2 | ? |
| Dental. Upper M1. Carabelli"s cusp. ASUDAS grades. UM1CB | 0 | 0 | ? | 0.5 | ? | 1 | ? | ? | 2.5 | 0 | 0 | ? |
| Dental. Upper M1. Mesostyle size. ASUDAS grades. UM1PS | 0 | 0 | ? | 0 | ? | 1 | ? | ? | 0 | 0 | 1.5 | ? |
| Dental. Upper M1. Enamel Extension. ASUDAS grades. UMEE | 0 | 0.5 | ? | 0 | ? | 0 | ? | ? | 0 | 0 | 0 | ? |
| Dental. Upper M2. Mesiodistal length. | 13.15 | 11.6 | ? | 11.5 | 13.6 | 11.1 | ? | ? | 13.23 | 12.4 | 11.3 | ? |
| Dental. Upper M2. Buccolingual width. | 16.86 | 12.3 | ? | 12.95 | 16.6 | 12.3 | ? | ? | 14.43 | 13.4 | 14.2 | ? |
| Dental. Upper M2. Width/length. | 128.05 | 106 | ? | 112.65 | 122.1 | 110.8 | ? | ? | 109.1 | 108.1 | 125.7 | ? |
| Dental. Upper M2. Area relative to M1. LxW of M2 / LxW of M1 | 104.8 | 106.4 | ? | 102.7 | ? | 95 | ? | ? | 99.9 | 96.1 | 91 | ? |
| Dental. Upper M2. Metacone size. ASUDAS grades. UM2MC | 3.5 | 2 | ? | 3 | 1 | 2 | ? | ? | 3.5 | 3 | 3 | ? |

| | | | | | | | | | | | | |
|---|---|---|---|---|---|---|---|---|---|---|---|---|
| Dental. Upper M2. Hypocone size. ASUDAS grades. UM2HC | 4 | 3 | ? | 3 | 3 | 3 | ? | ? | 4 | 3 | 3 | ? |
| Dental. Upper M2. Cusp 5 size. ASUDAS grades. UM2C5 | 3.5 | 0.5 | ? | 2.5 | 0.5 | 4 | ? | ? | 4.5 | 3.5 | 3 | ? |
| Dental. Upper M2. Carabelli"s cusp. ASUDAS grades. UM2CB | 0 | 0 | ? | 1 | 0 | 0 | ? | ? | 2 | 0 | 0 | ? |
| Dental. Upper M2. Mesostyle size. ASUDAS grades. UM2PS | 0 | 0 | ? | 0 | 0 | 0 | ? | ? | 0 | 0 | 0 | ? |
| Dental. Upper M2. Enamel Extension. ASUDAS grades. UMEE | 0 | 0.5 | ? | 0 | 1 | 1 | ? | ? | 0 | 0 | 0 | ? |
| Dental. Upper M3. Mesiodistal length. | 6.38 | 9.2 | ? | 7.6 | ? | 8.5 | ? | ? | 10.71 | 9.4 | 10.7 | ? |
| Dental. Upper M3. Buccolingual width. | 6.17 | 12.8 | ? | 9.2 | ? | 9.9 | ? | ? | 13.92 | 12.3 | 13.4 | ? |
| Dental. Upper M3. Width/length. | 96.7 | 139.1 | ? | 121.1 | ? | 116.5 | ? | ? | 13 | 130.9 | 125.2 | ? |
| Dental. Upper M3. Area relative to M1. LxW of M3 / LxW of M1 | 18.45 | 87.8 | ? | 48.2 | ? | 58.5 | ? | ? | 78 | 66.9 | 81.3 | ? |
| Dental. Upper M3. Metacone size. ASUDAS grades. As UM2MC | 0 | 1 | ? | ? | ? | 0.5 | ? | ? | 1.5 | 1 | 2 | ? |
| Dental. Upper M3. Hypocone size. ASUDAS grades. As UM2HC | 0 | 2 | ? | ? | ? | 1.5 | ? | ? | 2.5 | 1 | 2 | ? |
| Dental. Upper M3. Cusp 5 size. ASUDAS grades. As UM2C5 | 0 | 0 | ? | ? | ? | 2 | ? | ? | 3.5 | 0 | 0 | ? |
| Dental. Upper M3. Carabelli"s cusp. ASUDAS grades. As UM2CB | 0 | 0 | ? | ? | ? | 0 | ? | ? | 0 | 0 | 0 | ? |
| Dental. Upper M3. Mesostyle size. ASUDAS grades. As UM2PS | 0 | 0 | ? | ? | ? | 0 | ? | ? | 0 | 0 | 0.5 | ? |
| Dental. Upper M3. Enamel Extension. ASUDAS grades. UMEE | 0 | 0.5 | ? | ? | ? | 0 | ? | ? | 0 | 1 | 0 | ? |

**Taxonomic scope for comparison and phylogenetic analyses**

We scored and measured morphological characters from 104 cranial and mandibular specimens of the *Homo* genus (S-Table 4). Specimens from the same locality, with similar date range and with similar morphology and generally accepted as the same species/population were grouped into one operational taxonomic unit (OTU). Such an OTU can be considered a paleodeme and has more informative characters for phylogenetic analysis. In modern phylogenetics, a phylogeny is a model that represents the evolutionary relationships among OTUs. The OTUs can represent many types of comparable taxa, such as a family of organisms, individuals, populations, or even virus strains (*46, 47*). An OTU is not necessarily a species. Our phylogenetic analysis is at the population level. When there are multiple specimens from the same locality with the same age range, we have combined these specimens into a paleodeme. Many *Homo* fossils are represented by a single individual, such as Petralona, Ceprano, Dali, Jinniushan, Harbin et al. We considered these *Homo* fossils to be representative of the populations from those localities. Since fossilization and fossil finding are all rare events, it is reasonable as a first assumption that the discovered fossils most likely represent the typical morphological condition of the population that once lived in these localities. Finding a variant with a rare morphology is statistically an even rarer event. In our phylogenetic analysis, we did not group fossils from different localities and different ages into previously known *Homo* species, such as *H. erectus, H. heidelbergensis, H. neanderthalensis, H. sapiens*, or *H. longi,* although this is a common palaeoanthropological practice, and usually accompanied by much debate. As we have emphasized, our phylogenetic analysis is at the population (palaeodeme) level, and we wish to see how the individual fossils group, without prior assumptions. The *a priori* combination of fossils into species without a reliable phylogeny can obscure evolutionary processes.

After combination, 61 OTUs were used as terminal taxa for the phylogenetic and biogeographic analyses. The OTUs cover all the major clades or forms of the *Homo* genus (S-Table 4). For each terminal taxon/specimen, we use the most recently published dating results. For the combined terminal taxon, dates for all the specimens were used as an age range.

S-Table 4. Specimens used for comparison and phylogenetic analysis

| OTUs* | Specimens and common taxonomic allocation | Specimen source** | Estimated or determined age (kyr) | Estimated or measured cranial capacity (ml) | Age Reference | Cranial Capacity Reference |
|---|---|---|---|---|---|---|
| Antecessor | *H. antecessor* ATD6-15 ATD6-69 ATD6-96 | NHM replicas; IVPP AN2245 | 949000-772000 | 1000 | Ref. (*48*) | Ref. (*49*) |
| | *H. antecessor* Dental | | 949000-772000 | | Ref. (*48*) | |
| Narmada | *Homo* sp. Narmada | NHM replica; IVPP AN1631 | 780000-236000 | 1155-1421 | Ref. (*50*) | Ref. (*51*) |
| Irhoud | *H. sapiens* Irhoud1 | NHM replica | 349000-281000 | 1369-1381 | Ref. (*52*) | Ref. (*53*) |
| | *H. sapiens* Irhoud2 | NHM replica | 349000-281000 | 1467-1473 | Ref. (*52*) | Ref. (*53*) |
| Florisbad | *H. sapiens* Florisbad | NHM replica | 294000-224000 | 1280 | Ref. (*54*) | Ref. (*55, 56*) |

| | | | | | | |
|---|---|---|---|---|---|---|
| Omo 2 | *H. sapiens* Omo 2 | NHM replicas | 195000-255000 | 1487-1495 | Ref. (*53*) | Ref. (*53*) |
| LH 18 | *H. sapiens* LH18 | NHM replicas and CT data | 300000-150000 | 1232-1242 | | Ref. (*53*) |
| Skhul | *H. sapiens* Skhul 5 | NHM replicas | 130000-90000 | 1362-1364 | | Ref. (*53*) |
| | *H. sapiens* Skhul 9 | NHM replicas | 130000-90000 | 1400-1587.33 | | Ref. (*56, 57*) |
| Qafzeh | *H. sapiens* Qafzeh 9 | NHM replicas | 120000-90000 | 1492-1502 | | Ref. (*53*) |
| Mladec | *H. sapiens* Mladec 1 | IVPP AN1961, AN1628 | 40000 | 1606 | | Ref. (*57*) |
| | *H. sapiens* Mladec 2 | NHM replica | 40000 | 1390 | | Ref. (*57*) |
| | *H. sapiens* Mladec 5 | NHM replica | 40000 | 1500-1650 | | Ref. (*56, 57*) |
| | *H. sapiens* Mladec 6 | NHM replica | 40000 | | Ref. (*53*) | |
| Cro Magnon | *H. sapiens* Cro-Magnon 1 | NHM replicas | 31000 | 1573-1575 | Ref. (*53*) | Ref. (*53*) |
| | *H. sapiens* Cro-Magnon 2 | | 31000 | | Ref. (*53*) | |
| | *H. sapiens* Cro-Magnon 3 | NHM replica | 31000 | 1781-1845 | Ref. (*53*) | Ref. (*53*) |
| Oase | *H. sapiens* Oase1 | IVPP AN2250-2, replica | 41470-39410 | | Ref. (*58*) | |
| | *H. sapiens* Oase2 | IVPP AN2250-1, replica | 41470-39410 | 1400-1500 | Ref. (*58*) | This research |
| ZKD UC | *H. sapiens* Upper Cave 101 | IVPP AN59, AN62, replica | 39000-36300 | 1500 | Ref. (*59*) | Ref. (*57*) |
| | *H. sapiens* Upper Cave 103 | IVPP AN61, replica | 39000-36300 | 1290-1300 | Ref. (*59*) | Ref. (*57*) |
| Liujiang | *H. sapiens* Liujiang | IVPP PA89, original | 33000-23000 | 1567 | Ref. (*59*) | Ref. (*60*) |
| SH | *H. neanderthalensis* SH4 | NHM replica; IVPP AN2183-1, replica | 430000 | 1360 | Ref. (*18*) | Ref. (*61*) |
| | *H. neanderthalensis* SH5 | IVPP AN2183-2, replica | 430000 | 1092 | Ref. (*18*) | Ref. (*61*) |
| Tabun 1 | *H. neanderthalensis* Tabun C1 | NHM original | 170000 | 1270.5-1271 | | Ref. (*56, 57*) |
| Tabun 2 | *H. sapiens* Tabun C2 | NHM replica | 300000-170000 | | | |
| Spy | *H. neanderthalensis* spy 1 | NHM replicas | 40000 | 1278-1296 | Ref. (*53*) | Ref. (*53*) |
| | *H. neanderthalensis* Spy 2 | | 40000 | 1524-1538 | Ref. (*53*) | Ref. (*53*) |

| | | | | | | |
|---|---|---|---|---|---|---|
| Forbes' Quarry (Gibraltar) | *H. neanderthalensis* Gibraltar 1 (=Forbes' Quarry 1) | NHM original | 75000 | 1209-1217 | Ref. (*53*) | Ref. (*53*) |
| Amud | *H. neanderthalensis* Amud | NHM replicas | 53000 | 1731-1763 | Ref. (*53*) | Ref. (*53*) |
| La Chapelle | *H. neanderthalensis* La Chapelle aux Saints | IVPP AN2301-1, AN2301-2 replica and CT data | 52000 | 1487-1493 | Ref. (*53*) | Ref. (*53*) |
| La Ferrassie | *H. neanderthalensis* La Ferrassie1 | NHM replica | 70000 | 1638-1648 | Ref. (*53*) | Ref. (*53*) |
| Shanidar | *H. neanderthalensis* Shanidar1 | NHM replica | 75000-45000 | 1650 | | Ref. (*56, 62*) |
| | *H. neanderthalensis* Shanidar5 | NHM replica | 75000-45000 | 1550 | | Ref. (*56, 62*) |
| Césaire | *H. neanderthalensis* St Césaire | NHM replicas | 41950-40660 | | Ref. (*63*) | |
| Saccopastore | *H. neanderthalensis* Saccopastore 1 | NHM replica | 250000 | 1234.3 | Ref. (*56, 62*) | Ref. (*56, 62*) |
| | *H. neanderthalensis* Saccopastore 2 | NHM replica | 250000 | 1295 | Ref. (*56, 62*) | Ref. (*56, 62*) |
| Neanderthal type | Neanderthal 1 | NHM replica | 42000 | 1337.8 | Ref. (*56, 62*) | Ref. (*56, 62*) |
| Xiahe | *Homo* sp. Xiahe | Original | 155000-164500 | | Ref. (*64*) | |
| Denisova | *Homo* sp. Denisovans | Denisova 3, 4, 8 | 51600-122700 | | Ref. (*65*) | |
| Xujiayao | *Homo* sp. Xujiayao | IVPP PA 1490, 1495, 1498, 1496, 1480, 1481, 1497, 1500, original | 160000-200000 | 1555-1787 | Ref. (*66*) | Ref. (*66*) |
| Yunxian | *Homo* sp. Yunxian | EV 9001, EV 9002, original | 936000-1100000 | 1143 | Ref. (*5*) | This research. |
| Dali | *Homo* sp. Dali | IVPP AN1369, original | 267700-258300 | 1120 | Ref. (*67*) | Ref. (*68*) |
| Hualongdong | *Homo* sp. Hualongdong | IVPP Hualongdong, original | 331000-275000 | 1150 | Ref. (*69*) | Ref. (*69*) |
| Harbin | *Homo* sp. Harbin | HBSM2018-000018(A), original | 309000-146000 | 1400 | This research | This research |
| Jinniushan | *Homo* sp. Jinniushan | IVPP AN2118, original | 310000-200000 | 1390 | Ref. (*60*) | Ref. (*60*) |

| | | | | | | |
|---|---|---|---|---|---|---|
| Maba | *Homo* sp. Maba | IVPP AN629, original | 278000-230000 | 1300 | Ref. (*70*) | Ref. (*71*) |
| Xuchang | *Homo* sp. Xuchang | IVPP replica, original | 125000-105000 | 1800 | Ref. (*72*) | Ref. (*72*) |
| Mauer | *H. heidelbergensis* Mauer 1 | NHM replica | 649000-569000 | | Ref. (*73*) | |
| Arago | *Homo* sp. Arago 21, 47 | NHM replicas | 469000-407000 | 1138.667-1166 | Ref. (*74*) | Ref. (*56, 57*) |
| | *H. Homo* sp. Arago 13 | NHM replicas | 469000-407000 | | Ref. (*74*) | |
| | *Homo* sp. Arago 2 | NHM replicas | 469000-407000 | | Ref. (*74*) | |
| Kabwe | *H. rhodesiensis* Broken Hill | NHM E 686, original | 324000-274000 | 1249 | Ref. (*75*) | Ref. (*53*) |
| Petralona | *Homo* sp. Petralona1 | NHM replica | 400000-150000 | 1160-1164 | Ref. (*53*) | Ref. (*53*) |
| Ceprano | *Homo* sp. Ceprano | Original, CT data | 430000 | 1185 | Ref. (*56, 62*) | Ref. (*56, 62*) |
| Saldanha | *Homo* sp. Saldanha (=Elandsfontein) | NHM replicas | 700000-350000 | 1216.667 | Ref. (*56, 62*) | Ref. (*56, 62*) |
| Bodo | Bodo | NHM replica | 600000 | 1200-1325 | Ref. (*56, 62*) | Ref. (*76*) |
| Tighenif (formerly Ternifine) | *Homo* sp. Ternifine | NHM replica | 750000 | | Ref. (*55*) | Ref. (*55, 56, 62*) |
| | *Homo* sp. Ternifine 2 | NHM replica | 750000 | | Ref. (*55*) | |
| | *Homo* sp. Ternifine 3 | NHM replica | 750000 | | Ref. (*55*) | |
| | *Homo* sp. Ternifine 4 | NHM replica | 750000 | 1300 | Ref. (*55*) | |
| Peking | *H. erectus* Peking 10 | IVPP AN1, replica | 580000-280000 | 1225 | Ref. (*60, 77*) | Ref. (*60*) |
| | *H. erectus* Peking 12 | IVPP AN3, replica | 580000-280000 | 1030 | Ref. (*60, 77*) | Ref. (*60*) |
| | *H. erectus* Peking 13 | IVPP AN55, replica | 580000-280000 | | Ref. (*60, 77*) | |
| | *H. erectus* Peking 52 | IVPP AN22, replica | 580000-280000 | | Ref. (*60, 77*) | |
| | *H. erectus* Peking RC1996 | IVPP AN742-1, AN742-2, replica | 580000-280000 | 1030 | Ref. (*60, 77*) | Ref. (*60*) |
| Nanjing 1 | *H. erectus* Nanjing1 | IVPP AN1353, original | 620000-580000 | 876 | Ref. (*60*) | Ref. (*60*) |

| | | | | | | |
|---|---|---|---|---|---|---|
| Hexian | *H. erectus* Hexian | IVPP AN1368, original | 437000-387000 | 1025 | Ref. (*78*) | Ref. (*60*) |
| Sambungmacan | *H. erectus* Sambungmacan1 | NHM replica and CT data | 200000 | | Ref. (*53*) | |
| | *H. erectus* Sambungmacan3 | NHM replica and CT data | 200000 | 898-906 | Ref. (*53*) | Ref. (*53*) |
| Sangiran | *H. erectus* Sangiran2 | NHM replica | 1500000-1300000 | 789-797 | Ref. (*53*) | Ref. (*53*) |
| | *H. erectus* Sangiran17 | NHM replica | 1500000-1300000 | 1020 | Ref. (*53*) | Ref. (*56, 62*) |
| Ngandong | *H. erectus* Ngandong 7 | IVPP AN166 | 117000-108000 | 1013 | Ref. (*79*) | Ref. (*57*) |
| | *H. erectus* Ngandong 9 | IVPP AN345 | 117000-108000 | | Ref. (*79*) | |
| | *H. erectus* Ngandong 12 | IVPP AN170 | 117000-108000 | 1127 | Ref. (*79*) | Ref. (*57*) |
| Dmanisi | *H. erectus* Dmanisi 211 2282 | NHM replicas | 1770000 | 650 | Ref. (*80*) | Ref. (*80*) |
| | *H. erectus* Dmanisi 2280 | IVPP AN2181 replica | 1770000 | 730 | Ref. (*80*) | Ref. (*80*) |
| | *H. erectus* Dmanisi 2700 2735 | NHM replicas | 1770000 | 601 | Ref. (*80*) | Ref. (*80*) |
| | *H. erectus* Dmanisi 4500 2600 | NHM replicas | 1770000 | 546 | Ref. (*80*) | Ref. (*80*) |
| StW 53 | *Homo* sp. STW53 | NHM replica, StW 53, original, CT data | 1900000 | 570 | Ref. (*56, 62*) | Ref. (*56, 62*) |
| OH 9 | *H. erectus* OH9 | NHM replica; IVPP AN748, original, CT data | 1470000 | 1009-1017 | Ref. (*53*) | Ref. (*53*) |
| Turkana | *H. erectus* Turkana | KNM-WT15000, original, CT data | 1535000 | 846-854 | Ref. (*53*) | Ref. (*53*) |
| | *H. erectus* ER 3733 | KNM-ER 3733, original, CT data | 1780000 | 876-880 | Ref. (*53*) | Ref. (*53*) |
| | *H. erectus* ER 3883 | KNM-ER 3883, original, CT data | 1570000 | 837-839 | Ref. (*53*) | Ref. (*53*) |

| | | | | | | |
|---|---|---|---|---|---|---|
| Habilis | *H. habilis* OH24 | NHM replica; IVPP AN2179, original | 1800000 | 597 | Ref. (*56, 62*) | Ref. (*56, 62*) |
| | *H. habilis* OH7 | NHM replica; IVPP AN2292, original, CT data | 1780000 | | Ref. (*56, 62*) | Ref. (*56, 62*) |
| | *H. habilis* ER1805 | KNM-ER 1805, original, CT data | 1850000 | 616 | Ref. (*56, 62*) | Ref. (*56, 62*) |
| Naledi | *H. naledi* Lesedi | UW 102b-511, 102a-011, LES1, 102a-089, 102c-240, 102a-023, original, CT data | 236000-335000 | 610 | Ref. (*81, 82*) | Ref. (*82*) |
| | *H. naledi* Naledi | UW DH1-3, 101-1473, 101-1277, 101-1261, 101-377, DH3, original, CT data | 236000-335000 | 465 | Ref. (*81, 82*) | Ref. (*1*) |
| Flores | *H. floresiensis* | LB1, LB6/1, replica | 50000-190000 | 426 | Ref. (*83*) | Ref. (*84*) |

* OTUs: operational taxonomy units. ** KNM: National Museum of Kenya; StW: Sterkfontein, University of the Witwatersrand; NHM: Natural History Museum, London; IVPP: Institute of Vertebrate Paleontology and Paleoanthropology; HBSM: Hebei Science Museum; UW: University of the Witwatersrand; LES: Lesedi Chamber (U.W.102) in the Rising Star system; DH: Dinaledi Hominin from the Dinaledi Chamber of the Rising Star cave system; LB: Liang Bua cave on the island of Flores, Indonesia.

**Morphological data collection**

Character state scoring and linear and angular measurements were taken at the specimen level. Definitions of discrete characters and metric measurements were updated based on our recent phylogenetic work (*85*). Most of the 244 discrete characters are widely used and discussed in palaeoanthropological researches (e.g. in the resources of Ref. (*61, 80, 86-91*)). We revised and re-defined the characters, and presented illustrations for those not in outright or unmistakable definition. The continuous characters include 255 linear measurements and 22 angles. The 128 ratios included in our previous analyses, which are derived from the linear measurements (*85*), were excluded from our current analyses, following the request of one reviewer of the manuscript. The linear and angular measurements were taken following the standards defined by Martin (*92*) and Howells (*93*). X-ray CT images and the surface models were exported to the VG Studio Max 3.2, and all the measurements were taken from the digital 3D models in VG Studio Max 3.2. Gross morphology of hominid dentitions shows extensive variation. Without considering these variations, discrete characters cannot fully reflect the dental traits. As we have suggested that the wide range of variation should be scored on a ranked scale (*85*), here, the morphological traits of the permanent upper and lower dentitions of *Homo* fossils were scored by using the standard of the Arizona State University Dental Anthropology System (*94*). This ranking system is widely accepted as a standard (*95, 96*), and it has been used to infer the phylogenetic relationships of hominids (*97*).

Discrete character definitions, linear and angle measurement standards, original scoring and measurements are stored in MorphoBank, which is a publicly available web application and database widely used for large-scale, online morphological character standardization and data collection (*98*). To standardize the characters, scoring and measurements, we used the labeling tools of MorphoBank (*99*) to illustrate the anatomical features and homology. MorphoBank Current project is an update of the MorphoBank Project 3385. By following the methods of scoring in Ref. (*99*), we loaded and labelled media to document the exemplars of the characters and measurements, and recorded how the states of discrete characters looked in each specimen. Two hundred and forty-four discrete characters and 405 continuous characters were defined and scored or measured for the 104 *Homo* specimens. In total, 1626 media, 10682 labels and 24529 cell scorings were input in MorphoBank Project 4045, which is an update of MorphoBank Project 3385.

**Characters for phylogenetic analysis**

The 244 discrete characters were all equally weighted. Forty-six multi-state characters were set as "ordered". When the scored specimens were merged into a terminal taxon, their character states were also merged. The merged cells with multiple states were set to polymorphism.

Continuous characters can capture quantitative variation that discrete characters do not fully represent by providing a more comprehensive view of phenotypic or ecological evolution, and thus can provide additional insight into evolutionary processes (*100-102*). The use of continuous characters in phylogenetic analyses can improve morphological phylogenetics (*103-105*). Linear measures are usually size related due to allometric and/or isometric effects. While not all studies explicitly state that continuous characters must be transformed, there is a consensus in the literature that transformations are often necessary to meet the assumptions of phylogenetic comparison methods and to ensure biologically meaningful results (*100, 101, 106*). Often used data transformation methods include allometric scaling, residual analysis, geometric means, use ratios or indices. Here we converted all linear measurements of the skull and upper dentition of a scored specimen into ratios by dividing the measurements by the 1/3 power of the cranial capacity or maximum frontal width of that specimen. Cranial capacity or maximum frontal width was chosen as the denominator because it could be measured in most specimens. The linear measurements of the mandibles and lower dentitions of a scored specimen were divided by the bi-ramus breadth at the alveolar margin of this specimen.

Cranial capacity was chosen as a reasonable proxy for body mass. It is a general phenomenon that linear measurements are allometrically related to body mass. However, estimating the body mass of an extinct animal always involves much uncertainty and debate. Estimating cranial capacity, on the other hand, is more direct and less debatable. The close relationship between body mass and cranial capacity, which in turn is closely related to brain size, has been demonstrated in many classical analyses (e.g. Ref. (*107-109*)). The result of a linear measurement divided by the cube root of the cranial capacity of the same individual can therefore be interpreted as the relative size of this measurement in relation to the body mass of this examined individual. Similarly, the maximum frontal width, or the bi-ramus breadth of a mandible, can also be regarded as a reasonable proxy for the body mass.

After the transformation of the linear measures into ratios, all the linear measurement ratios and angle variables were normalized to further reduce the potential correlations among all metric characters. Given a variable, a value of this variable minus the minimum of the variable, then the result was divided by the difference between the maximum and minimum of this variable among all the scored specimens. After transformation and normalization, all the continuous characters have a range between 0 and 1.

In total 277 normalized continuous characters and 244 discrete characters were used for phylogenetic analysis. The names of the characters were listed in Appendix 1 and Appendix 2.

**Test of potential correlations among continuous characters**

Because the distribution of each character cannot be guaranteed to be normal, we used the non-parameter Kendall correlation analysis to test the potential correlation between continuous characters. Due to the large number of missing values, we calculated the pairwise Kendall Tau. Among 277 continuous characters, 10 characters have too few pairwise values with other characters for calculating Kendall Tau. These characters were removed from correlation analysis.

The results of correlation analysis of the continuous characters shows that the majority of pairwise correlations between characters are weak. Only 0.3% are strong (Kendall Tau > 0.8), and 9.8% can be considered moderately strong (0.8 > Kendall Tau > 0.5). We do not see a particularly strong correlation between linear measurements and ratios. We did not remove or reduce the weight for one of the strongly correlated pairs during the phylogenetic analysis, because we believe that the related characters reflect the relative importance of morphology. This is equivalent to increasing the weight for important characters.

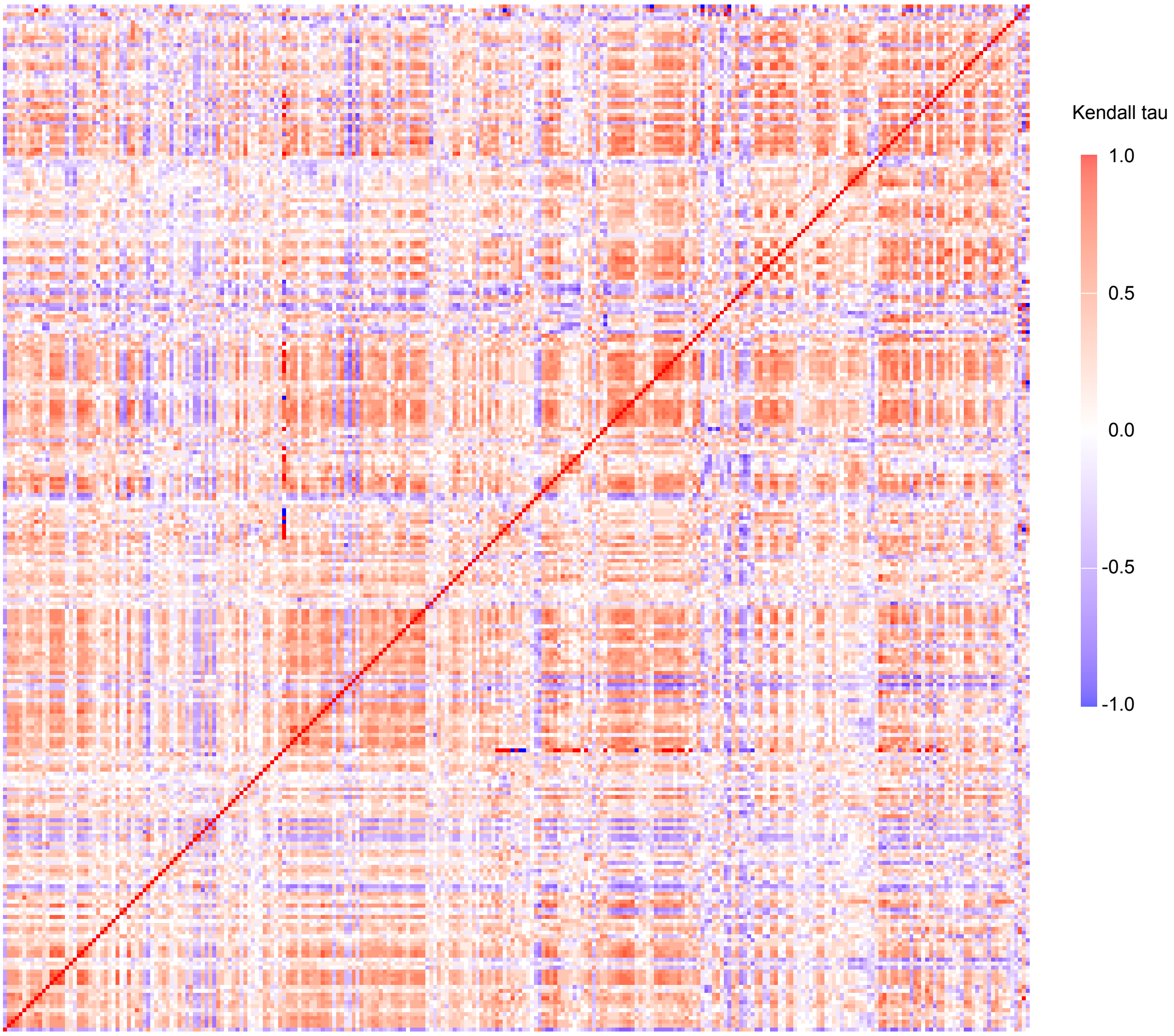

S-Figure 21. Non-parameter Kendall correlation analysis among 267 continuous characters. Each color dot represents a pairwise Kendall Tau.

**Parsimony analysis**

Parsimony analysis of the dataset (discrete and continuous, Appendix 3) was undertaken by using TNT, Tree analysis using New Technology, a parsimony analysis program subsidized by the Willi Hennig Society (*31*). We used the parallel version of TNT on one hundred CPU cores. We ran multiple replications, using sectorial searches, drifting, ratchet and fusing combined (Appendix 4). Random sectorial search, constraint sectorial search and exclusive sectorial search were set with default settings. Ten cycles of tree drifting, 10 cycles of ratchet and 10 cycles of tree fusing were performed in the search. The search level was set as 10 for 61 or 60 (without Denisovan) taxa. Optimal scores were hit 10 times independently, each hit with 1000 initial replications. In total 1 million replications were performed (10000 replications on each core). Some characters are set as ordered (Appendix 4, 5). All characters have equal weight. No constraint was used for the parsimony analysis. About 20 hours were required to finish the non-constraint parsimony search on our computing cluster. More than 3488 billion rearrangements were examined. Fifty-two trees with a best score of 2512.31 were retained.

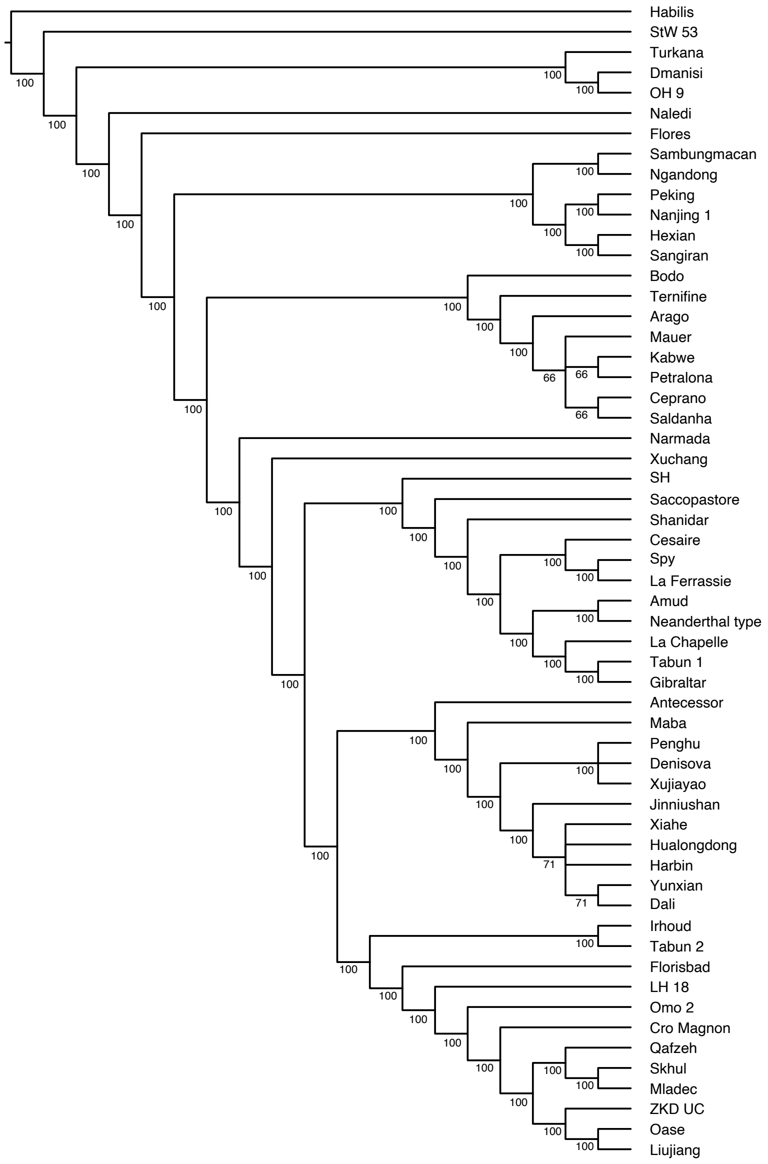


S-Figure 22. The preferred majority rule consensus tree of the 42 most parsimonious trees. Linear measurements of cranium and upper dentition were transformed into ratios by dividing the measurements by the 1/3 power of the cranial capacity. Numbers near internal nodes indicate majority consensus percentage.

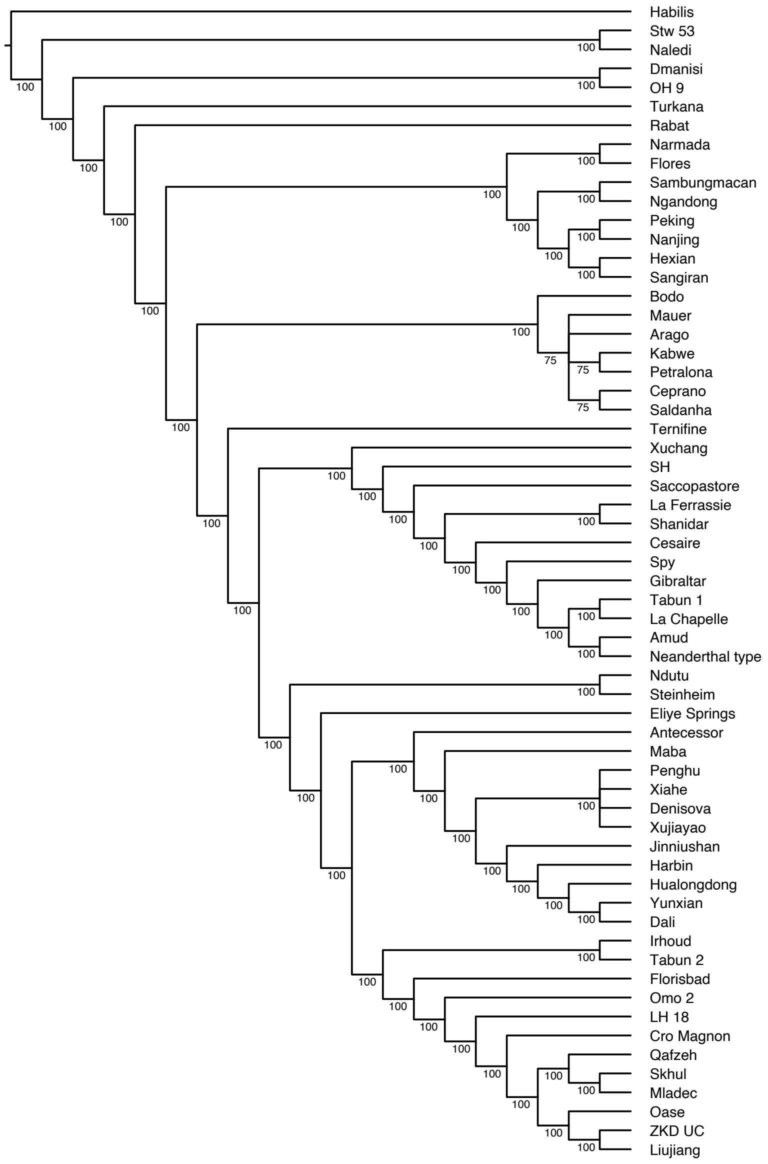


S-Figure 23. Majority rule consensus tree of the 24 most parsimonious trees including Eliye Springs, Ndutu, Steinheim and Rabat. Linear measurements of cranium and upper dentition were transformed into ratios by dividing the measurements by the 1/3 power of the cranial capacity. Numbers near internal nodes indicate majority consensus percentage. The main topological feature of this tree is identical to the tree excluding the 4 fossils (as in S-figure 22).

Fossils such as Eliye Springs, Ndutu, Steinheim and Rabat are either incomplete, distorted, surface eroded, or in a pathological state. Some morphological landmarks are difficult to identify. Previous phylogenetic analyses have shown that different tree search strategies can lead to different positions of these fossils (*85*). In our current analyses, we removed the four OTUs. When they were included in the parsimony analysis, the result does not change the major tree topology (S-Figure 23), particularly the relationships among Neanderthal, *sapiens* and *longi* monophyletic clades are identical to the preferred most parsimonious tree excluding the 4 OTUs.

When the linear measurements of cranium and upper dentition were transformed into ratios by dividing them by the maximum frontal width, the parsimony analysis revealed almost identical tree topology as the one based ratios transformed by dividing by the 1/3 power of the cranial capacity (S-Figure 24). The latter analysis strategy is preferred because the transformation covers more available data and including fewer missing data.

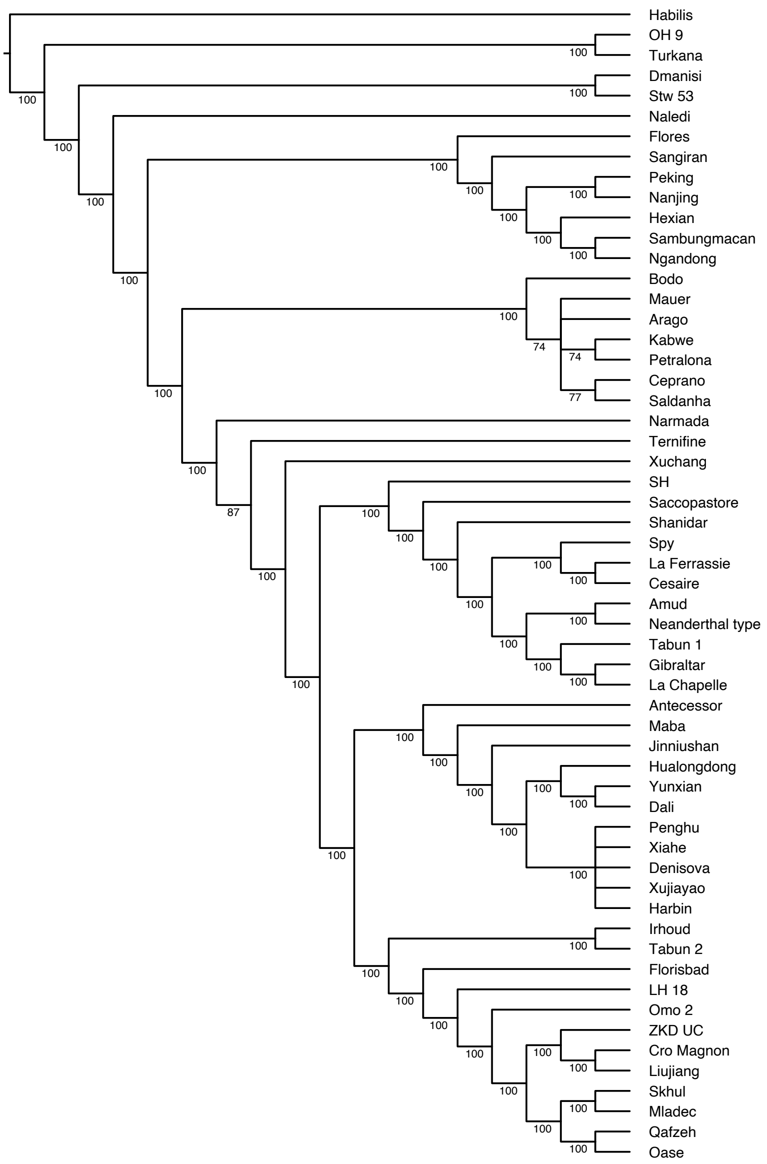


S-Figure 24. Majority rule consensus tree of the 31 most parsimonious trees. Linear measurements of cranium and upper dentition were transformed into ratios by dividing the measurements by the maximum frontal width. Numbers near internal nodes indicate majority consensus percentage. The main topological feature of this tree is identical to the tree based on the linear measurements of cranium and upper dentitions were transformed into ratios by dividing the measurements by the 1/3 power of the cranial capacity (as in S-figure 22).

**Test of influence of potential correlations among characters on the phylogeny**

The characters used for phylogenetic analysis are assumed to be independent and identically distributed. However, this assumption is generally accepted but rarely tested, especially for nucleotide characters (*110-112*). Following the most common practice in morphological data analyses, in which most characters can be assumed to be independent, both discrete and continuous characters studied here were treated as independent data points, and thus no correlations among characters were considered. However, some characters are likely to be correlated due to their anatomical structure or synergy in function. When we built the data matrix, we consciously avoided redundant and potentially correlated discrete characters. Ratio transformation and normalisation of the continuous characters can significantly reduce the potential correlations. As parsimony has no explicit model assumptions, the consequence of ignoring character correlation is hard to predict. One obvious consequence would be overestimating the number of changes (parsimony length) in the tree and probably would probably aggravate long-branch attraction. In Bayesian tip-dating analysis, the overestimation of character changes is reflected in the branch lengths each of which is a product of divergence time and evolutionary rate. With sufficient fossils and relatively accurate ages, the divergence time estimates would be less affected while resulting in accelerated evolutionary rates. The ignorance of character correlation would also include erroneous or overconfident topological inference (*113*), although simulation studies showed that the estimate is relatively robust (*104*). Some studies also show that when the correlation is low, treating the characters as independent still can produce reliable estimates of topology and times (*104, 114*). Further efforts are still needed in model developments for morphological characters.

How to evaluate the confidence that one should be in a given phylogenetic tree, and how to measure the support for phylogenetic trees are interesting topic in the process of reconstruction of phylogenetic trees (*111, 115*).

We used three resampling methods and Bremer support analysis to evaluate the stability of the phylogenetic tree, particularly to test how much the topology of the phylogeny changes as the proportion of character duplication is increased. The first resampling test is the symmetric resampling (*116*). In symmetric resampling, a certain proportion of characters (probability 2p) are changed: either duplicated or deleted with equal probability. The probability p is set to 0.05, 0.1, 0.2, 0.3, 0.4, 0.5, corresponding to 5% to 50% of the characters being changed. The resampling results were mapped on the majority consensus of the most parsimonious trees inferred from the original data matrix. We found that the increase in probability p, which represents the proportion of character duplication, caused an increase in the uncertainty of the phylogeny topology, especially among the basal clades. However, the supports are mainly positive (S-Figure 25) and the major clades identified in the most parsimonious tree remain stable. For the probability p = 0.05 and 0.1, the resampling results positively support almost all clades identified in the most parsimonious tree. The distribution of supports also shows a similar pattern (S-Figure 26).

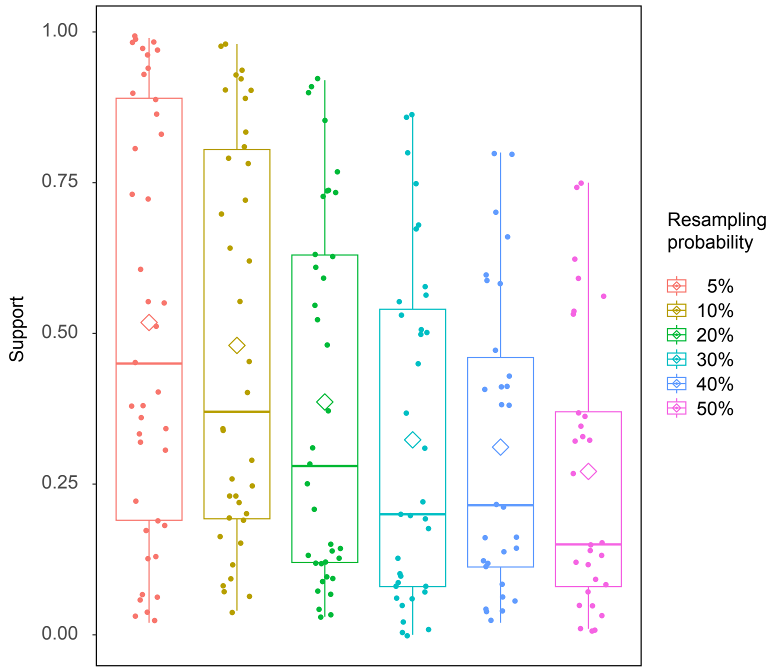


S-Figure 25. Symmetric resampling analyses with different resampling probability. Resampling trees were mapped on to the majority consensus of the most parsimonious trees inferred from the original data matrix. Supports of different resampling strength are mainly positive.

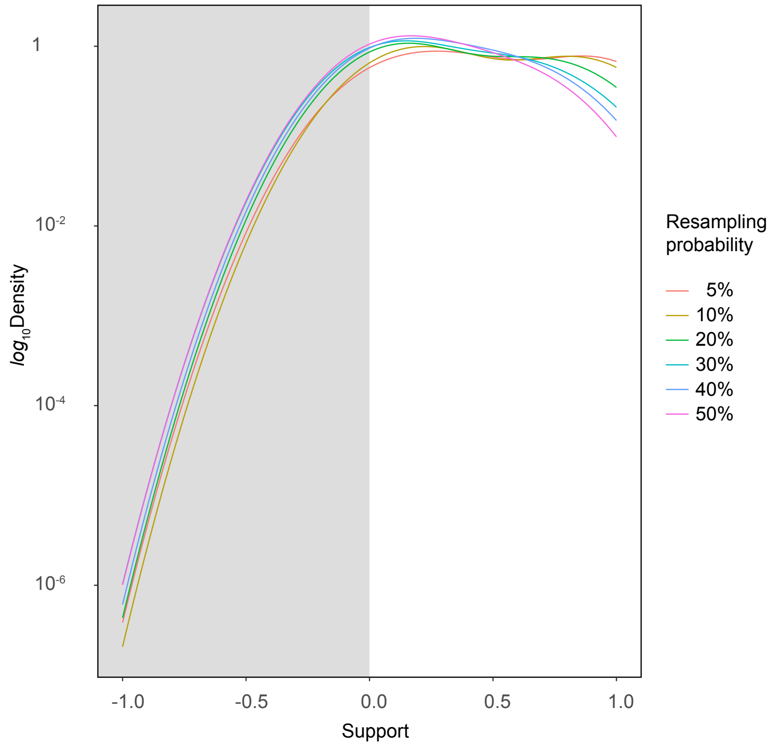

S-Figure 26. Symmetric resampling analyses with different resampling probability. Resampling trees were mapped on to the majority consensus of the most parsimonious trees inferred from the original data matrix. The distribution of supports of different resampling strength shows a similar pattern.

The second resampling method is random character duplication. It is similar to the symmetric resampling, but only duplicates characters and does not delete any characters. The probability p is also set to 0.05, 0.1, 0.2, 0.3, 0.4, 0.5. The resampling results show that the topology of the phylogenetic tree is almost identical to the most parsimonious tree for 5%, 10%, 20% and 30% character duplication (S-Figure 27). For 40% and 50% character duplication, the resolution of the tree decreases, but the general patterns are consistent with the most parsimonious tree (S-Figure 27).

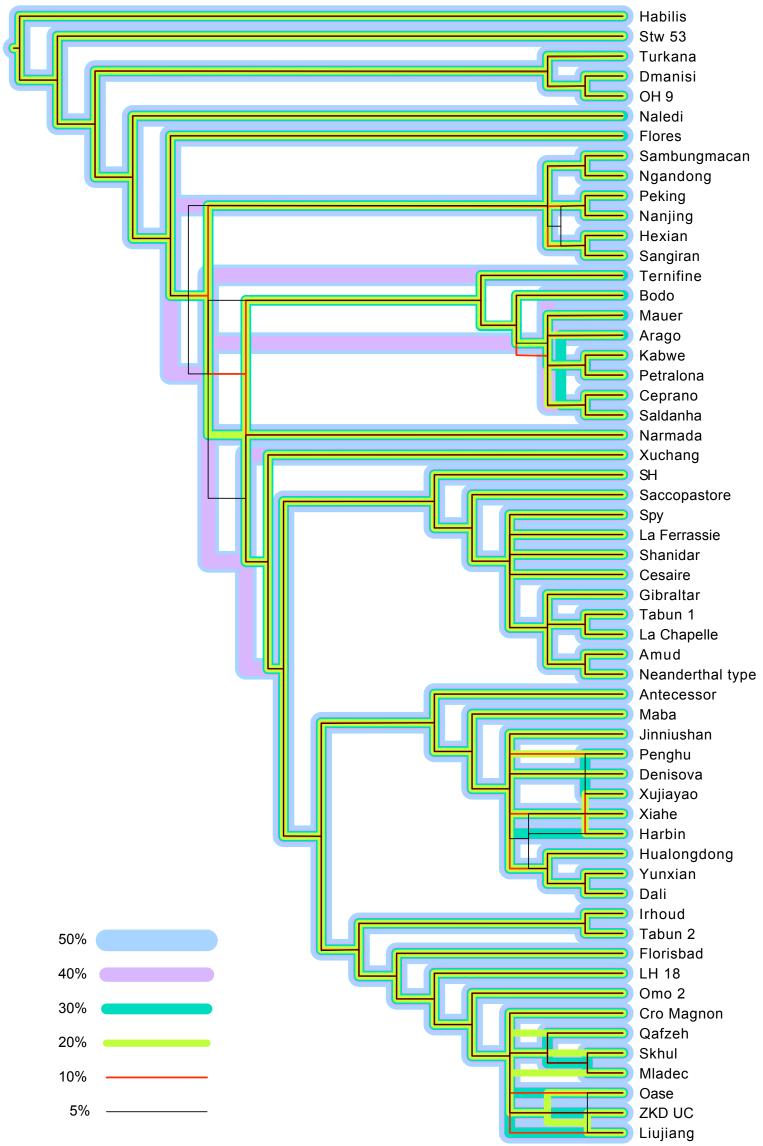

S-Figure 27. Random duplication resampling analysis shows that major clades can be identified even with very high resampling strength. The topology of the phylogenetic tree is almost identical to the most parsimonious tree for 5% ~ 30% character duplication.

The third resampling method is bootstrap resampling (*100*), which is integrated into TNT (*117*). The resampling strength of bootstrap resampling is equivalent to that of symmetric resampling when the probability p of symmetric resampling is set to 0.5, which corresponds to 50% of characters changing. We used the transfer bootstrap expectation (TBE, gradual transfer distance (*118*)) to measure the support of branches (S-figure 28). We used a parallel version of TNT and ran the bootstrap resampling on 100 cores. The bootstrap resampling yielded 1 million trees. The TBE based on these 1 million trees suggests that most nodes of the phylogeny are well supported with a TBE greater than 70%. If the Felsenstein bootstrap percentage or TBE of a node is greater than 70%, the node is considered to be strongly supported (*118, 119*). With such a criterion, most nodes are strongly supported.

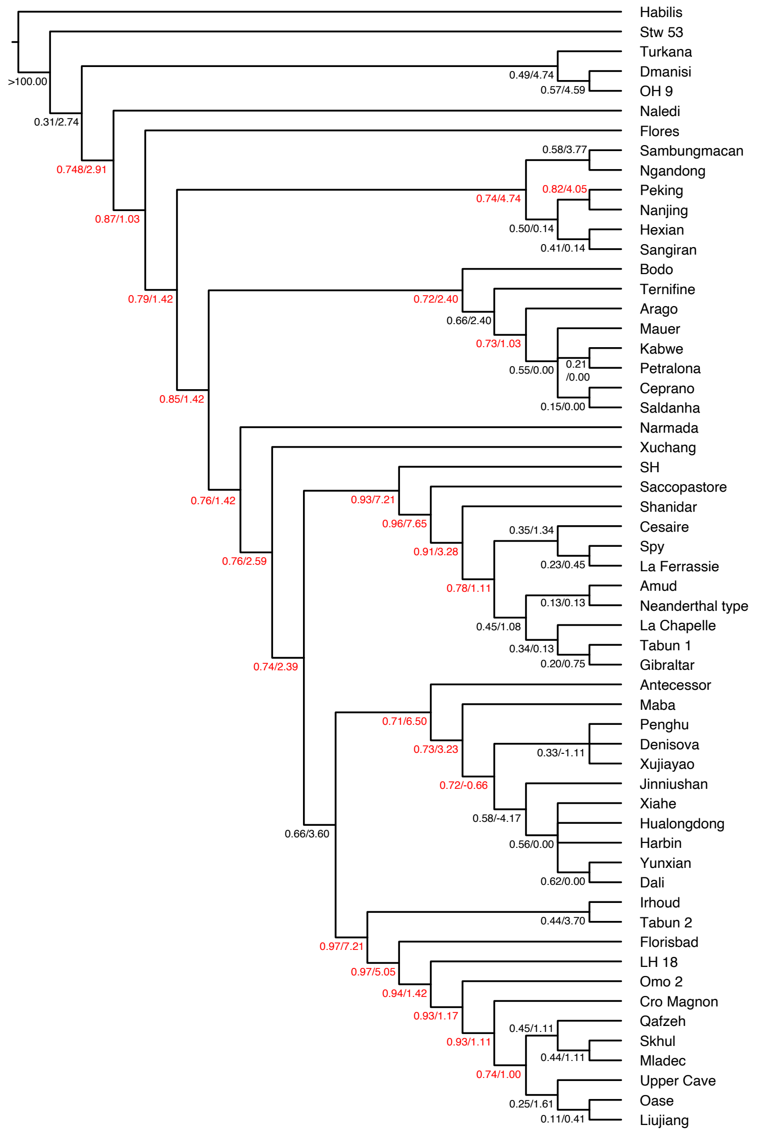

S-Figure 28. The preferred majority rule consensus tree of the 42 most parsimonious trees (as in S-Figure 22). Numbers before the slashes are the TBE (*118*) of 1 million trees of the bootstrap analysis (*100*). Numbers after slashes are the Bremer supports (*120*). Number in red color indicates that the bootstrap support TBE is greater than 70% and the node is strongly supported (*118, 119*).

Resampling methods used in phylogenetic analysis, such as bootstrap, jack-knifing, and symmetric analysis, all resample the characters (*110, 116, 121*) and involve perturbations of the data. Bremer support (*120, 122*) analysis is a completely different approach, which examines how many extra steps are needed to lose a branch in the consensus tree of near-most parsimonious trees. The method was first proposed by Farris et al. ( (*123*)) for distance cluster analysis and by Bremer ( (*122*)) for parsimony analysis. Since Bremer support analysis does not modify or perturb the original data matrix, it will not introduce arbitrary changes or false variation in the trees. Since it was proposed, Bremer support has become one of the two main methods for assessing the stability of phylogenetic trees and has been cited thousands of times. We used TNT (*117*) to calculate Bremer supports (Appendix 5 for script) to describe the stability of the phylogenetic results (S-figure 28). As the trees were generated based on both discrete and metric characters, only the absolute Bremer supports can be calculated, and the relative Bremer supports are not applicable. Bremer supports for most of the nodes are larger than 1, indicating that the tree is very stable.

**Effect of the equally-weighted character subgroups on phylogeny**

The characters in our phylogenetic analysis can be divided into several subgroups: 167 neurocranial, 160 visceral cranial, 145 mandibular, 175 dental, and 2 postcranial. Except for the postcranial characters, the other subgroups contain approximately the same number of characters. To test the robustness of the phylogeny when these character subgroups are changed, a phylogenetic analysis with one or more subgroups removed will certainly result in a different tree topology due to the reduction of informative characters. We did not test how robust the most parsimonious tree is when a subgroup is removed, because a tree based on cranial characters only or dental characters only certainly cannot be considered a tree supported by all available evidence.

Instead of removing some subgroups, we reweighted the characters so that all subgroups (except the postcranial, which has too few characters) have the same weight. The result is slightly different from the most parsimonious tree without adding weights, but all major clades identified in the most parsimonious trees remain unchanged (S-Figure 29). The consensus of the most parsimonious trees without adding weights is preferred, because it requires fewer assumptions.

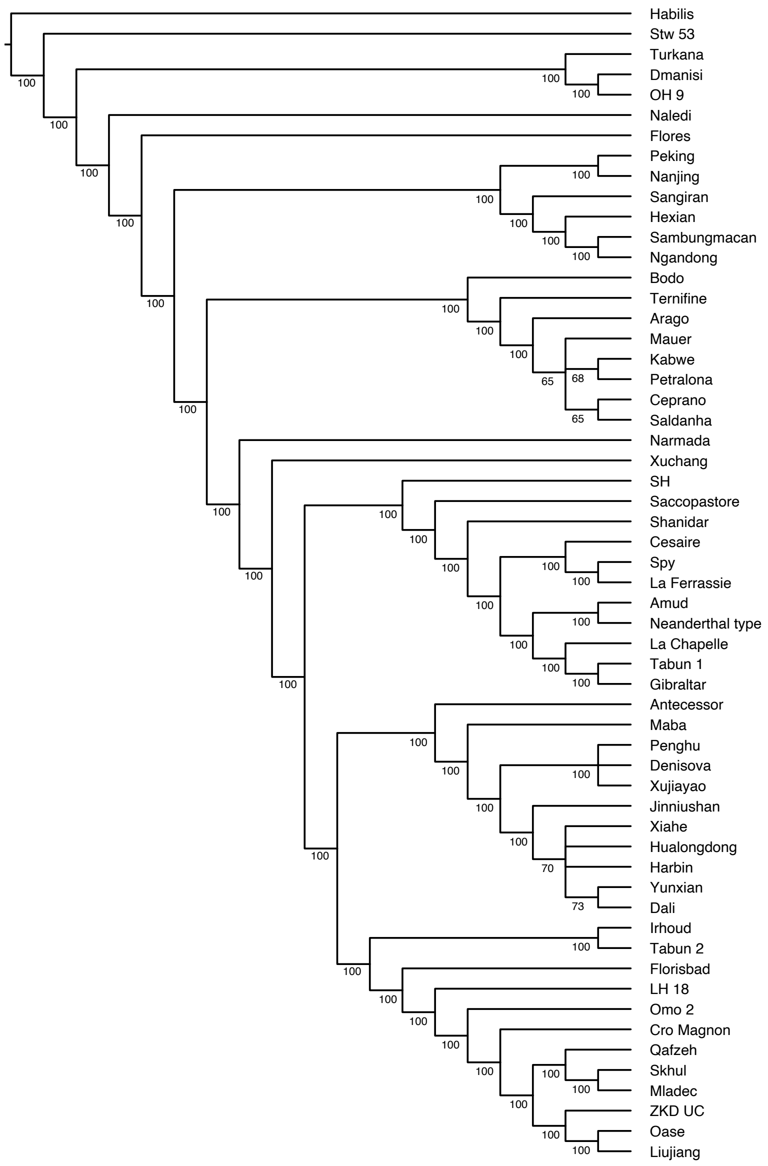


S-Figure 29. Majority rule consensus tree of the 41 most parsimonious trees, with neurocranial, visceral cranial, mandibular, dental, and postcranial character subgroups being equally weighted. The main topological feature of this tree is identical to the most parsimonious tree without reweighted the character subgroups (as in S-figure 22). The numbers near the internal nodes are the percentages of consensus.

**Bootstrap analysis of the phylogenetic position of Yunxian 2**

Reinstatement and reconstruction are based on morphological comparison of previously known Chibanian hominin crania and empirical judgement of the natural shape of the bones. Although multiple restorations and reconstructions were made and an average was selected for subsequent morphological study and phylogenetic analysis, it is impossible to estimate how different the reconstruction is from the actual undeformed cranium. However, we can use a resampling method to assess how reliable the phylogenetic position of Yunxian 2 is, when estimated from our reconstruction.

We can assume that some of the metric characters measured and discrete character states scored from the reconstruction of Yunxian 2 are inaccurate because of errors in the reconstruction. We do not know exactly how many of these inaccurate characters there are, but we can assume that they could reach up to 50% of the total characters. Therefore, we randomly removed 50% of the total metric and discrete scores of Yunxian 2, and then used TNT parsimony analysis to infer the phylogenetic position of Yunxian 2. This random removal of metric character scores from Yunxian 2 and the parsimony analysis were repeated 1000 times. Finally, all parsimony trees found during the 1000 iterations of phylogenetic analyses were summarized and evaluated using TBE (*118*).

This bootstrap resampling analysis indicated that the phylogenetic position of Yunxian 2 was strongly supported with a TBE of 89% (S-Figure 30), suggesting that errors in the reconstruction of Yunxian 2 (if any) do not affect the phylogenetic position. This means that our current reconstruction of Yunxian 2 is suitable for phylogenetic analysis.

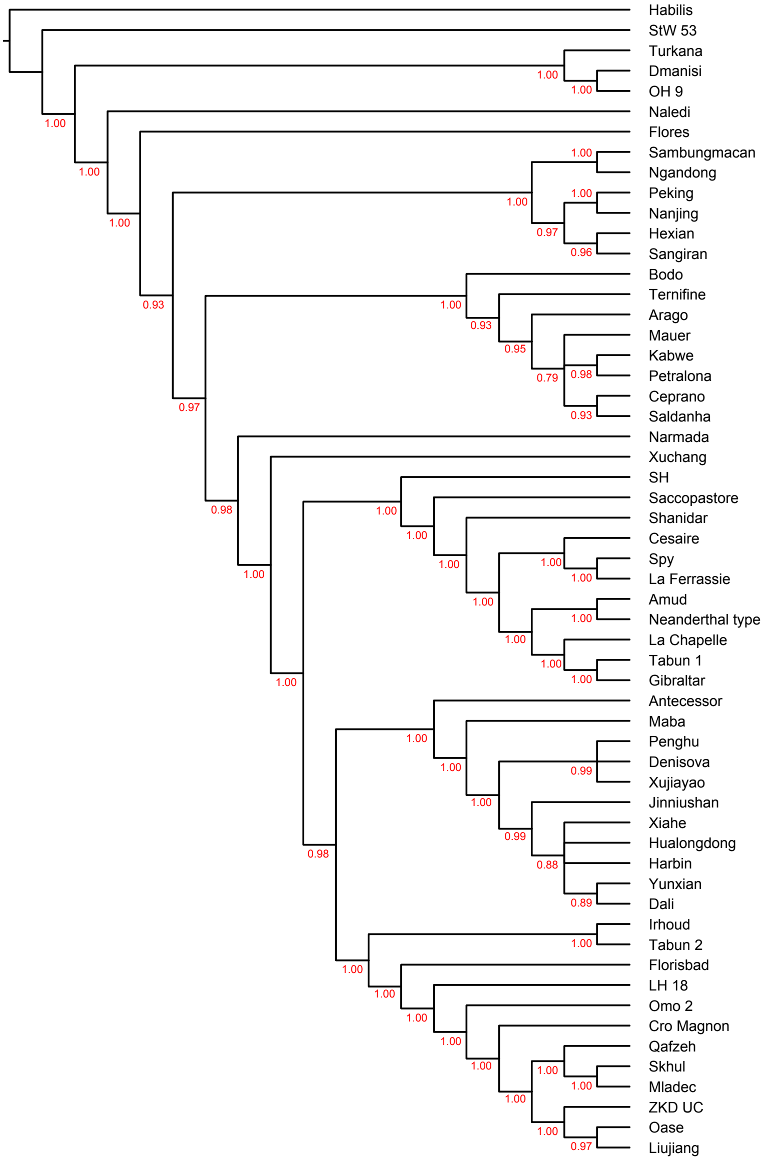

S-Figure 30. Bootstrap resampling analysis by randomly removing 50% of the total metric and discrete scores of Yunxian 2. Bootstrap trees were summarized on the preferred majority consensus tree of the most parsimonious trees (as in S-Figure 22). Numbers at the internal nodes are the TBE (*118*) of 1477 trees of the bootstrap analysis (*100*).

**Bayesian tip-dating analyses**

Bayesian inference for phylogeny reconstruction based on the current morphological data matrix of the genus *Homo* cannot provide as good a resolution as parsimony analysis. Therefore, we used the Bayesian tip-dating approach (*105, 124-126*) implemented in MrBayes 3.2.7 (*32*) to infer the divergence dates and evolutionary rates of the preferred majority consensus of the most parsimonious trees (as in S-Figure 22). Both fossil ages and morphological data were integrated in a coherent analysis. Discrete and continuous characters are treated as two data partitions. The Lewis Mk model with variable ascertainment bias correction (*127*) was used for the discrete data, and gamma rate variation across characters (*128*) (Mkv+Γ) was used for the likelihood calculation. 46 characters were defined as ordered and the rest of them (188 characters) were unordered (See the Characters for phylogenetic analysis section). Because MrBayes 3.2.7 cannot handle continuous characters directly and can deal with ordered characters only up to six states, all the continuous characters were discretised into six states. This is done by first dividing the range of 0 and 1 into six equal-length intervals (numbered as 0 to 5) and then converting each trait value into a state according to its interval assignment (Appendix 6). The discretised continuous characters were all defined as ordered to fit the nature of gradual change and modelled under Mkv+Γ. By default, the gamma shape of the Mkv+Γ model (*128*) was assigned an exponential (1.0) prior. The gamma shape models rate variation within each partition. The evolutionary rate variation among the two data partitions were accounted for using a uniform Dirichlet prior (*125*). The prior for the time-tree was modelled by the fossilized birth-death (FBD) process (*125, 129-131*). The process is conditioned on the time of the most recent common ancestor (root age) and has hyperparameters of speciation rate, extinction rate, fossil-sampling rate and extant-sampling probability. The root age was assigned an offset-exponential prior with a mean age of 3600 kyr and minimum age of 2800 kyr, referring to the potentially oldest fossil of *Homo habilis* (*132, 133*) and the beginning of the Late Pliocene epoch. The ages of the fossil tips were either fixed or given uniform distributions based on the corresponding stratigraphic ranges. The speciation, extinction and fossil-sampling rates were reparametrized for convenience (*125, 130*). The net diversification rate (speciation rate minus extinction rate) was assigned an exponential (200) prior with mean 0.005 which ranges from zero to infinity and puts less weight on higher rate. The turn-over rate (extinction rate over speciation rate, which is between 0 and 1) and relative fossil-sampling rate (fossil-sampling rate over the sum of extinction rate and fossil-sampling rate, which is also between 0 and 1) were both assigned a uniform (0,1) prior. The extant-sampling probability was fixed to 1.0 by default.

Apart from the time-tree, the relaxed clock model was used. With this model, the evolutionary rate varies along the branches in the tree. We used the white noise (WN) (*134*) model, in which the branch rates follow independent gamma distributions. The mean clock rate was assigned an exponential 100) prior (about 1 changes per 1000 characters per thousand years) and the variance parameter of the clock rate was exponential (2). As the discrete and continuous characters probably have distinct patterns of change through time, we unlinked the clock variance in these two partitions so that the evolutionary rate varies independently between partitions.

We executed two independent runs and 4 chains per run (1 cold and 4 hot chains with temperature 0.05) in the Markov chain Monte Carlo (MCMC) simulation. Each run was executed with 100 million iterations and sampled every 2000 iterations. The first 30% of samples were discarded as burn-in and the rest from two runs were combined. Good convergence and mixing were diagnosed by an effective sample size (ESS) (*135*) larger than 200 for all parameters and the average standard deviation of split frequencies (ASDSF) (*32*) smaller than 0.02. The posterior trees were summarized to both 50% majority-rule consensus tree and all-compatible consensus tree. The MrBayes commands are provided in Appendix 7. The analysis took about 55 hours using the parallel version of MrBayes on our computing cluster.

The Bayesian inference for the phylogeny of Yunxian and other *Homo* in this research does not provide clear resolution among some basal clades. Most of the clades discovered in parsimony analyses can also be discovered in our Bayesian inference, but the basal relationships among these clades cannot be resolved. In our previous Bayesian inference for phylogeny, we used parsimony backbone constraints, which directly treat continuous characters without discretization, to guide the tree topology search, increase the resolution of the phylogenetic relationships of basal diversification, and reduce polytomies in the majority-rule consensus tree (*85*). In our current analysis, we used Bayesian tip-dating to infer divergence dates only, and the tree topology was fixed to the preferred majority consensus of the most parsimonious trees, but we did not impose any date constraint in the Bayesian tip-dating analysis. This practice can be viewed as: given a limited number of proposed phylogenetic tree models (consistent with the results of parsimony analysis in this particular case), we use Bayesian tip-dating to estimate the divergence time of the internal nodes of these proposed trees.

It has long been known that some stochastic models of character change, when implemented in a maximum likelihood framework, provide a correspondence between the maximum parsimony method and the maximum likelihood method (*136-138*). The Zhang et al. 2020 method (*139*) was inspired by the linear relationship between parsimony scores and probability (*138*). Huelsenbeck et al. (*138*) found that the integrated likelihood is a rescaling of the parsimony score for a tree, and the marginal posterior probability distribution of the length of a branch depends on how the maximum parsimony method reconstructs the characters at the inner nodes of the tree. As a possible consequence of the linear relationship between parsimony scores and probability, they also found that trees sampled by the MCMC algorithm are similar to maximum parsimony trees. Zhang et al. (*139*) found that MCMC chains using parsimony-guided moves converge faster than chains using standard moves. They also exhibit better mixing and cover the most probable trees more quickly. This parsimony-guided tree proposal method was included in MrBayes 3.2 as a default set of tree moves based on promising preliminary experiments (*32, 139*). Although Bayesian inference and parsimony analysis are two phylogenetic methods based on two different sets of assumptions, previous research has shown that it is possible to mathematically link the methods. In practice, we think it makes sense to combine the two methods in order to obtain a more harmonious phylogenetic model.

Some recent analyses based on ancient DNA have produced relatively younger estimates of Neanderthal-*H. sapiens* divergence dates (*140*). The approach is also tip dating -- using ancient DNA sequences as data and their ages as tip dates. Although there were abundant molecular sequences in their analysis, it does not necessarily mean that their estimates are more reliable than ours. Theoretical studies have shown that even with infinitely long molecular sequences (or infinitely many morphological characters analogously) so that the branch lengths (distances measured by expected number of substitutions per site) can be inferred without error, the divergence times and evolutionary rates are confounded and rely on the information of fossil ages (or calibration priors) and clock models to get resolved (*141, 142*). They only included Neanderthals, Denisovans and *H. sapiens* with the oldest being the Sima de los Huesos early Neanderthals (~ 430 kyr), and lack information to inform divergences near the root of the genus *Homo*. We included all the major clades of the *Homo* genus, and thus we have more information from fossil ages to inform the divergence times of all the *Homo* clades. They also fixed the mutation rate (*140*) and thus put apparent certainty in the clock model, which might also bias the age estimates, while in our study, the clock rate was co-estimated with the divergence times from the tip-dating analysis. The FBD model that we used explicitly models the speciation, extinction and sampling processes and is more suitable for our data than the coalescent model (*143*) used by Posth et al. (*140*), which is better suited for a single population without population structure.

**Synapomorphies inferred from parsimony analysis**

The preferred majority consensus of the most parsimonious trees was described in TNT. The synapomorphies shared by the *longi* clade, *sapiens* clade and Neanderthal clade were calculated based on parsimony criterion . The results were listed in S-Table 5-7. Given a phylogenetic tree and using the maximum parsimony criterion, the state changes of a character across the tree are independent of other characters. For example, given a tree, the state changes of dental morphology are independent of the state changes of cranial characters, regardless of their states and whether they are missing or not. We emphasize that the phylogenetic tree proposed in this research is only a model based on currently available evidence. The synapomorphies of the lineages are also based on the current best understanding of the model.

S-Table 5. Nine synapomorphies shared by the *longi* clade (not including *Homo antecessor*)

| Characters | Plesiomorphic states | Apomorphic states |
|---|---|---|
| Character 0: Cranial capacity | 0.615-0.621 | 0.630 |
| Character 5: Glabella-bregma chord. g-b chord | 0.343-0.346 | 0.347 |
| Character 40: FRF. Nasion-subtense fraction. Frontal | 0.468-0.472 | 0.506 |
| Character 106: M49a. DKB. Interorbital breadth (d-d) | 0.648-0.670 | 0.597 |
| Character 126: M77a. NFA. Nasio-frontal angle (fm:a-n-fm:a) | 0.451-0.516 | 0.447 |
| Character 405: Frontal. Anterior view. Supraorbital tori arching in anterior view | gently arching | strongly arching |
| Character 420: Frontal. Lateral view. Supraorbital torus smoothly rolled | absent | present |
| Character 427: Frontal 4. Anterior view. Glabellar inflexion | Shallow | Deep |
| Character 435: Frontal. Double temporal line on the frontal | Absent | Present |

S-Table 6. Forty-four synapomorphies shared by the sapiens clade

| Characters | Plesiomorphic states | Apomorphic states |
|---|---|---|
| Character 13: M6. ba-sphba. Basilar length | 0.377 | 0.479 |
| Character 15: M8. XCB. Maximum cranial breadth | 0.354 | 0.300 |
| Character 16: M8c. Squama suture breadth | 0.425-0.442 | 0.345 |
| Character 17: Maximum cranial breadth at supramastoid crest. Cranial vault | 0.402 | 0.344 |
| Character 24: M11. AUB. Biauricular breadth | 0.372 | 0.355 |
| Character 25: M11b. Biradicular breadth | 0.395 | 0.382 |
| Character 26: M12. ASB. Biasterion breadth (ast-ast). Temporal | 0.478-0.520 | 0.417 |
| Character 34: Maximum bimastoid breadth | 0.396 | 0.370 |
| Character 35: M20. Porion-bregmatic projective height | 0.466-0.488 | 0.575 |

| | | |
|---|---|---|
| Character 36: AVH. Auriculare-vertex projective height | 0.451-0.472 | 0.504 |
| Character 45: M30. PAC. Parietal sagittal chord | 0.527-0.618 | 0.644-0.698 |
| Character 47: M30(3). Lambda-asterion chord (l-ast) | 0.312-0.322 | 0.273 |
| Character 49: M30c. Bregma-asterion chord (b-ast). Parietal | 0.604 | 0.746 |
| Character 58: M32(1). Frontal inclination angle (b-n-i). Frontal | 0.586-0.590 | 0.637-0.698 |
| Character 59: M32(2). Bregma angle (b-g-i). Frontal | 0.560-0.580 | 0.668-0.706 |
| Character 61: M33d. OCA. Occipital angle (degree) | 0.689-0.697 | 0.705-0.741 |
| Character 64: Temporal squama length (Martínez and Arsuaga, 1997) | 0.207-0.277 | 0.360-0.396 |
| Character 65: Temporal squama height (Martínez and Arsuaga, 1997) | 0.455 | 0.297 |
| Character 76: Postglenoid-ectoglenoid length | 0.493-0.533 | 0.577-0.784 |
| Character 79: Chord length of the parietomastoid suture. Incisura parietalis - asterion | 0.371-0.465 | 0.352 |
| Character 93: Supraorbital torus thickness medial. Frontal | 0.298-0.401 | 0.428 |
| Character 105: M48d. WMH. Cheek height | 0.282-0.315 | 0.230-0.271 |
| Character 115: Maxilloalveolar length | 0.366-0.370 | 0.327 |
| Character 125: Subspinale subtense | 0.171-0.181 | 0.159 |
| Character 137: Infraorbital plate angle in parasagittal plane | 0.392 | 0.375 |
| Character 138: Orbital incline angle in parasagittal plane | 0.685-0.738 | 0.760 |
| Character 147: Notch width | 0.341-0.346 | 0.443-0.669 |
| Character 149: Condylar neck length | 0.377-0.589 | 0.619-0.638 |
| Character 162: Mandibular corpus height at p4 | 0.582-0.616 | 0.680 |
| Character 442: Maxillary. Anterior view. Lateral view. Paranasal inflation. Maxilla superior lateral inflation of the bone surrounding the nasal aperture | Present | Absent |
| Character 462: Palatine. Maxillary. Ventral view. Palatine torus | Absent | Small, elevated for about 1-5 mm, limited to the palatine bones |
| Character 496: Temporal. Posterior inferior view. Mastoid process ventral projection relative to the temporal-occipital suture | Below the suture | Markedly below |
| Character 503: Temporal. Ventral view. Entoglenoid process | large | small |
| Character 516: Temporal. Ventral view. Tympanic plate orientation | More sagittally orientated | More coronally orientated |
| Character 517: Temporal. Lateral view. Tympanic plate thickness | moderate | thin |

| | | |
|---|---|---|
| Character 531: Mandible 1. Lateral view. Incurvatio mandibulae | absent | deep |
| Character 532: Mandible 2. Lateral view. Symphyseal profile. Relative to the lower dental border | Receding | Convex |
| Character 533: Mandible 3. Anterior view. Central part of the mental protuberance | absent | strong |
| Character 534: Mandible 4. Anterior view. Mental tubercle | absent | strong |
| Character 566: Mandible. Lateral view. Coronoid process orientation | posteriorly projecting | anteriorly projecting |
| Character 567: Mandible. Lateral view. Condyle height relative to the coronoid | lower | sub equal |
| Character 568: Mandible. Anterior view. Canine pillar | absent | moderate |
| Character 572: Mandible 37. Medial view. Internal coronoid pillar orientation | near vertical | oblique |
| Character 588: Mandible 52. Superior view, medial view. Symphysis planum alveolare (simian shelf) | present, small, or present, large | absent |

S-Table 7. Forty-eight synapomorphies shared by the Neanderthal clade

| Characters | Plesiomorphic states | Apomorphic states |
|---|---|---|
| Character 1: M1. GOL. g-op. Maximum cranial length | 0.326-0.363 | 0.185 |
| Character 7: Length of basal temporal | 0.339-0.353 | 0.281 |
| Character 9: Entire temporal bone length | 0.344 | 0.343 |
| Character 14: M7. FOL. Foramen magnum length (ba-o) | 0.504-0.575 | 0.268-0.374 |
| Character 27: M14. WCB. Minimum cranial breadth | 0.466-0.479 | 0.340-0.449 |
| Character 30: M13a. MDB. Mastoid width. Temporal | 0.228 | 0.216 |
| Character 31: Maximum mastoid width | 0.232 | 0.212 |
| Character 46: M30(2). Bregma-sphenion chord (b-sphn). | 0.481-0.600 | 0.402-0.450 |
| Character 49: M30c. Bregma-asterion chord (b-ast). Parietal | 0.578-0.604 | 0.567 |
| Character 53: Lambda-opisthocranion chord l-op | 0.445-0.642 | 0.419-0.433 |
| Character 56: Opisthocranion-opisthion chord | 0.377-0.389 | 0.509-0.558 |
| Character 57: Sphenobasion-opisthion length | 0.568-0.594 | 0.423 |
| Character 58: M32(1). Frontal inclination angle (b-n-i). Frontal | 0.586-0.590 | 0.539-0.570 |
| Character 68: Temporal muscle attachment length | 0.291-0.327 | 0.117-0.183 |
| Character 71: Tympanic axis angle | 0.502 | 0.540 |
| Character 73: Petrous axis angle | 0.589 | 0.606 |
| Character 77: Postglenoid-entoglenoid length | 0.451-0.546 | 0.350 |
| Character 102: M74(1). Clivus-alveolar plane angle | 0.491-0.549 | 0.554 |
| Character 124: M76a. SSA. Zygomaxillary angle (degree) | 0.268-0.294 | 0.109-0.193 |
| Character 125: Subspinale subtense | 0.171-0.181 | 0.206-0.245 |
| Character 130: Nasal bridge angle. Rightmire 1998 | 0.446-0.556 | 0.620-0.630 |

| | | |
|---|---|---|
| Character 139: bimandibular fossa breath | 0.507-0.541 | 0.499 |
| Character 145: Gonion condylar height | 0.503-0.696 | 0.479 |
| Character 165: Bimental breadth | 0.795-0.817 | 0.931 |
| Character 179: Internal breadth at canine along the alveolar margin | 0.601-0.650 | 0.667-0.696 |
| Character 198: Dental. Upper I1. Labial convexity. ASUDAS grades. UI1LC | 0.428 | 0.571 |
| Character 231: Dental. Upper M1. Buccolingual width | 0.223-0.299 | 0.127-0.173 |
| Character 239: Dental. Upper M2. Mesiodistal length | 0.252-0.411 | 0.101-0.149 |
| Character 275: Dental. Lower p3. Mesiodistal length | 0.161-0.268 | 0.109 |
| Character 280: Dental. Lower p4. Mesiodistal length | 0.245-0.482 | 0.069-0.100 |
| Character 285: Dental. Lower m1. Mesiodistal length | 0.177-0.376 | 0.054-0.090 |
| Character 286: Dental. Lower m1. Buccolingual width | 0.159-0.399 | 0.090 |
| Character 293: Dental. Lower m2. Mesiodistal length | 0.174-0.385 | 0.089-0.146 |
| Character 294: Dental. Lower m2. Buccolingual width | 0.196-0.299 | 0.192 |
| Character 302: Dental. Lower m3. Mesiodistal length | 0.151-0.160 | 0.090 |
| Character 405: Frontal. Anterior view. Supraorbital tori arching in anterior view | gently arching | strongly arching |
| Character 417: Frontal 6. Superior view. Supraorbital torus arching in superior view | gently arching | strongly arching |
| Character 420: Frontal. Lateral view. Supraorbital torus smoothly rolled | absent | present |
| Character 426: Frontal. Lateral view. Posttoral sulcus. Concavity of the region between supraorbital torus and frontal squama | moderate | deep |
| Character 428: Frontal. Mid-sagittal supraglabellary tubercle (Zeitoun, 2000) | Absent | Present |
| Character 438: Maxillary. Medial view. Lack of an ossified roof over the lacrimal groove | No | Yes |
| Character 440: Maxillary. Anterior view, medial view. Inferior concha covering the lacrimal groove | Present | Absent |
| Character 446: Maxillary. Ventral view. Anterior-posterior position of the root of the zygomaticoalveolar crest | M1-M2 | M2-M3 |
| Character 476: Occipital. Posterior inferior view. Superior nuchal line | Moderately developed | Highly elevated |
| Character 509: Temporal. Ventral view. Glenoid fossa overhang | Less than 50 percent | Equal or greater than 50 percent |
| Character 526: Nasal. Lateral view. Nasal root projecting | Deeply concave | At the same sagittal level as the glabella |
| Character 553: Mandible. Lateral view. Prominentia lateralis position relative to the tooth loci | m2-m3 | m3 |

| | | |
|---|---|---|
| Character 554: Mandible. Lateral view. Retromolar space shielded by the ramus | Completely shielded, shielding part of the last molar | No shielding, large space visible |
